# Optimizing *in vitro* Transcribed CRISPR-Cas9 Single-Guide RNA Libraries for Improved Uniformity and Affordability

**DOI:** 10.1101/2025.03.24.644170

**Authors:** Natanya K. Villegas, Yukiko R. Gaudreault, Abigail Keller, Phillip Kearns, James A. Stapleton, Calin Plesa

## Abstract

We describe a scalable and cost-effective sgRNA synthesis workflow that reduces costs by over 70% through the use of large pools of microarray-derived oligos encoding unique sgRNA spacers. These sub-pool oligos are assembled into full-length dsDNA templates via Golden Gate Assembly before *in vitro* transcription with T7 RNA polymerase. RNA-seq analysis reveals severe biases in spacer representation, with some spacers being highly overrepresented while others are completely absent. Consistent with previous studies, we identify guanine-rich sequences within the first four nucleotides of the spacer, immediately downstream of the T7 promoter, as the primary driver of this bias. To address this issue, we introduced a guanine tetramer upstream of all spacers, which reduced bias by an average of 19% in sgRNA libraries containing 389 spacers. However, this modification also increased the presence of high-molecular-weight RNA species after transcription. We also tested two alternative bias-reduction strategies: compartmentalizing spacers within emulsions and optimizing DNA input and reaction volumes. Both methods independently reduced bias in 2,626-plex sgRNA libraries, though to a lesser extent than the guanine tetramer approach. These advancements enhance both the affordability and uniformity of sgRNA libraries, with broad implications for improving CRISPR-Cas9 screens and optimizing guide RNA design for other CRISPR and nuclease systems.

## Introduction

CRISPR-Cas9 technology has revolutionized DNA manipulation, advancing high-throughput gene editing, functional genomics, genetic engineering, and large-scale functional screens. At the core of this technology are single-guide RNAs (sgRNAs) with 20-nucleotide spacer sequences that enable precise targeting of both coding and non-coding DNA, thanks to their high programmability (1). This versatility allows for multiplex functional genomics studies, where multiple genetic elements can be investigated simultaneously.

To fully harness this capability, these studies rely on sgRNA libraries containing thousands to hundreds of thousands of unique sgRNA spacers targeting diverse genomic regions (2). Libraries such as GeCKOv2 (3), Brunello (4), Sabatini human knockout (5), and human CRISPRi v2 (6) are commonly used for genome-wide CRISPR knockout (CRISPR-KO), interference (CRISPRi), and activation (CRISPRa) screens. These libraries typically feature sgRNA sequences synthesized as microarray-derived oligonucleotides and subsequently cloned into lentiviral vectors to transfect cell lines.

To reduce the costs associated with CRISPR screens using these large genome-wide sgRNA libraries such as GeCKOv2, numerous smaller, more cost-effective libraries have been developed (7). One such library contains four sgRNAs per gene (8), compared to the 10 used in the large genome-wide Sabatini library (5). Another is modular, consisting of three sublibraries with 2–4 sgRNAs per gene that are synthesized separately and can be combined to adjust sgRNA representation (9). However, even these smaller libraries are typically produced as distinct sublibraries rather than amplified from a larger pool.

In contrast, amplifying sgRNA subpools from microarray-derived oligos provides a more efficient and cost-effective approach for targeted screening. One parallel retrieval strategy generated 24 subpools, each containing 1,000 unique sgRNA sequences, with coverage ranging from 98.8% to 99.8% (10). Beyond its efficiency, this method leverages bulk pricing advantages, as a single pool of microarray-synthesized oligonucleotides can be subdivided into multiple libraries. For example, ordering five separate microarray-synthesized pools, each containing 1,000 oligos, costs $1,120 per pool, totaling $5,600. In contrast, ordering a single pool of 5,000 oligos costs $1,680, resulting in a 72% savings (Twist Bioscience). Furthermore, this approach enables the generation of multiple targeted sgRNA libraries from a single oligo pool. By subdividing microarray-derived oligo pools, researchers can design libraries tailored to specific cellular states or protein families, optimizing representation within individual subpools.

These examples outline a pathway for generating sgRNA libraries that are fully programmable and partitionable for targeted, cost-effective studies. While these methods offer substantial improvements, they typically rely on lentiviral vector delivery, posing challenges for certain cell types and contexts. While lentiviral vectors allow for stable sgRNA expression in human cell lines, their prolonged expression increases the risk of off-target editing (12). Additionally, random genomic integration can lead to mutagenesis and prolonged Cas9 expression, further exacerbating off-target effects (11, 12). In contrast, delivery of active ribonucleoprotein complexes (RNPs)—comprising of synthesized sgRNAs with Cas9 or catalytically inactivated Cas9 (dCas9)—has been shown to achieve higher editing efficiencies and reduced off-target effects compared to lentiviral plasmid expression, even in cells that are notoriously difficult to transfect (11–13). Additionally, RNPs can be produced from synthesized sgRNAs and dCas9 fused to small molecules or peptides, enhancing the recruitment of epigenetic proteins to DNA sites in mammalian cells (14). RNPs are also compatible with *in vitro* CRISPR-Cas9 applications, as seen in CasKAS, a method for identifying off-target sites via N3-kethoxal binding to single-stranded DNA. Synthetic sgRNA RNPs can be electroporated into cells or applied to DNA *in vitro*, highlighting their versatility (13).

To enable the use of RNPs, sgRNAs can be synthesized chemically or enzymatically with T7 RNA polymerase (T7 RNAP), allowing flexible design. Although chemical synthesis yields high-purity products, it is expensive and results in low yields, making it impractical for large-scale sgRNA libraries (12). Enzymatic synthesis is more affordable, produces high sgRNA yields, and is accessible using commercially available kits for *in vitro* transcription (IVT) by T7 RNAP. Although commercially available IVT kits provide an affordable means to synthesize individual sgRNAs, they are not a cost-effective solution for generating large-scale, multiplex sgRNA libraries. This limitation primarily arises from the need for specific DNA oligo inputs. Microarray-derived oligos yield only femtomole (fmol) to picomole (pmol) quantities per sequence, making them insufficient for direct large-scale IVT without amplification. While pooled, column-synthesized oligos (oPools) offer greater material yields, their cost is approximately tenfold higher than microarray-derived oligos, making them less practical for high-throughput applications. That said, Cas12a-Capture successfully used 11,438 full-length crRNAs compatible with Cas12a, ordered as two oPools (15), demonstrating that oPools are a viable input source for crRNA libraries. Finally, these IVT kits typically require single-stranded DNA (ssDNA) as input and are incompatible with double-stranded DNA (dsDNA) obtained from PCR amplification of sgRNA subpools (10).

In this work, we developed a highly customizable, scalable, and programmable *in vitro* method for producing sgRNA libraries using T7 RNAP. We first amplified subpools of sequences from thousands of unique microarray-synthesized DNA target oligos containing sgRNA spacers, taking advantage of affordable bulk pricing. To further reduce costs, we minimized oligo length by excluding the conserved sgRNA scaffold sequence required for Cas9 complexation (16). We then used Golden Gate Assembly (GGA) to ligate the dsDNA spacer fragment with the scaffold sequence, efficiently generating full-length templates for transcription (**Fig. 1a**).

**Figure. 1.**
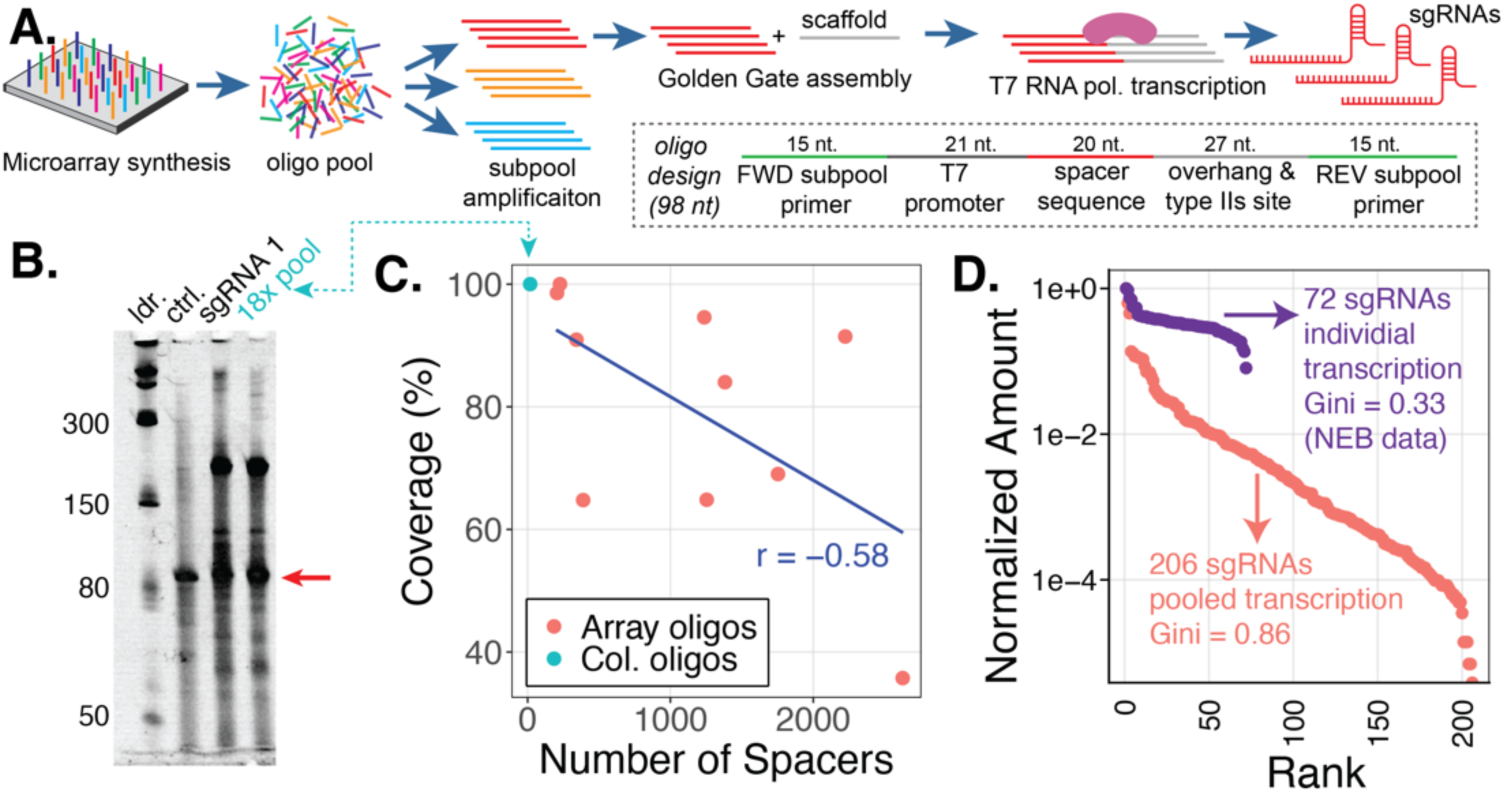
Experimental workflow and proof-of-concept for sgRNA library synthesis. **a.** We PCR-amplified microarray-derived 98-nt oligo subpools, each containing spacers corresponding to individual sgRNA libraries. Each oligo was designed with forward and reverse PCR primer sites, a T7 promoter followed by 1-2 guanines, unique 20 nucleotide spacer sequences, and a BsaI type IIs restriction site. Golden Gate Assembly (GGA) was used to add the conserved sgRNA scaffold sequence to the oligos. The resulting GGA products were used as templates for pooled IVT of sgRNA libraries using T7 RNAP. **b.** To evaluate the initial performance of our sgRNA synthesis workflow, we manually pooled 18 column-synthesized oligos with 18 unique spacer sequences, converting them to dsDNA by extending the reverse primer, followed by GGA and IVT. Successful synthesis of pooled and individual sgRNAs (100 bases) was confirmed by 10% TBE-urea denaturing electrophoresis, followed by SYBR-Gold post-staining. **c.** Ten sgRNA libraries containing 206 to 2,626 spacers were synthesized from microarray-derived (array) oligos. The percent coverage of the microarray-derived libraries (orange dots) was compared to the percent coverage of the 18-plex sgRNA library (teal dot) with column derived oligos. We find an inverse correlation between coverage and scale (Pearson correlation analysis, *r* = −0.58). **d.** Comparison of spacer distribution uniformity between a microarray-derived 206-plex sgRNA library and a library of 72 independently transcribed sgRNAs from New England Biolabs (NEB). Normalized abundance is shown, with spacers ranked in descending order.

GGA enables the ligation of multiple unique DNA fragments in a single reaction due to the activity of its type IIs restriction enzymes, which generate precise, user-specified overhangs (17). This approach offers distinct advantages over homology-based methods such as Gibson Assembly, uracil-specific excision reagent cloning (USER), and In-Fusion Cloning (Takara Bio). The DNA target oligos include a reverse primer site for subpooling that must be removed before assembly with the scaffold oligo. If left intact, this primer site remains adjacent to the scaffold oligo, displacing the spacer sequence that should be positioned at its 5’ end, thereby disrupting CRISPR targeting. To address this, a GGA restriction cut site was placed upstream of the reverse primer site, enabling a type IIs enzyme to cleave the priming site during DNA fragment assembly.

In contrast, homology-based methods would require multiple additional steps, including the digestion of unwanted primer sites, purification of DNA fragments, and hybridization of the two fragments. This increases workflow complexity and labor requirements, reducing the cost-effectiveness of adding the scaffold oligo separately. By introducing the conserved scaffold oligo after subpooling, we achieve an additional 14% cost savings. This contributes to the overall 72% reduction in cost achieved by subpooling microarray-derived oligos, as shorter oligo pools are more affordable. An additional advantage of GGA is that assembly is guided by short restriction sticky ends, whereas Gibson Assembly requires relatively long 20–40 nt homology regions for efficient assembly. Because the spacer region varies between sequences, designing consistent homology regions for Gibson Assembly would require extending the conserved sgRNA scaffold region beyond the spacer, significantly increasing oligo length and cost while also raising the risk of mis-hybridization (18, 19). While the USER method requires shorter overhangs for assembly, it is more complex and expensive compared to Gibson Assembly or GGA (18). Overall, these modifications led to substantial cost reductions while avoiding potential complications in our sgRNA synthesis process.

Once the DNA templates were assembled, we transcribed full-length sgRNAs *in vitro* using T7 RNAP and assessed metrics like spacer uniformity and coverage via RNA-seq analysis (**Fig. 1**). Our results validate the viability of this sgRNA synthesis workflow and reveal how IVT reaction conditions influence spacer distributions in sgRNA libraries. Since T7 RNAP preferentially transcribes certain sequences (20), we investigate how this bias affects spacer distribution. Specifically, spacer biases result in over-representation of certain spacers while under-representing others, potentially causing important functional hits to be missed. As a result, a significantly larger number of cells must be screened to achieve robust statistical power in CRISPR-based assays (21). Moreover, uneven sgRNA library coverage and uniformity can undermine the reproducibility and accuracy of these assays, limiting the reliability of their conclusions. To address this issue, we demonstrate the effectiveness of three distinct strategies to reduce this bias within sgRNA libraries containing 389 or 2,626 unique spacers and propose future directions for further optimization.

In summary, we present a scalable enzymatic sgRNA synthesis method optimized to improve spacer uniformity. This approach leverages T7 RNAP transcription to generate user-defined sgRNA libraries with reduced bias. As a result, our method provides a robust solution for *in vivo* CRISPR screening in human or mammalian cell lines and *in vitro* CRISPR assays.

## Results and Discussion

### Feasibility and Quality Assessment of sgRNA Library Synthesis

To establish an sgRNA library synthesis method independent of commercially available kits and protocols, we evaluated the feasibility of our custom workflow in two steps (**Fig. 1a**). First, we assessed IVT templates, produced by joining fragments containing 20-nucleotide (nt) sgRNA spacers with 83-nt scaffold fragments by Golden Gate Assembly (GGA), could reliably produce sgRNAs (**Fig. 1a**). This validation was crucial, as our T7 RNAP IVT protocol was developed by customizing existing methods, and its effectiveness required confirmation before scaling up to transcribe sgRNA libraries from microarray-derived subpools.

For our first proof-of-concept, we transcribed a pilot sgRNA library containing 18 unique spacer sequences. The 98-nt oligos were designed to contain a T7 promoter, 20-nt spacer, a type IIS restriction cut site (BsaI), and 15-nt forward and reverse primer sites (**Fig. 1a**). A single guanine was included at the 5’ end of spacers, downstream of the T7 promoter. This addition was made when the first nucleotide position was adenine, cytosine, or thymine, as it is necessary for T7 RNAP transcription initiation (22, 23). The 18 oligos were ordered as individual ssDNA oligos from Integrated DNA Technologies. The oligos were manually pooled to obtain an 18-plex oligo pool and duplexed by performing reverse single primer extension (RSPE). These dsDNA fragments were joined to the sgRNA scaffold by Golden Gate Assembly, producing 135 bp templates. The 18-plex sgRNA library was transcribed alongside a singleplex IVT of sgRNAs (sgRNA 1), also present in the 18-plex library as spacer 1. Successful synthesis of the 18-plex sgRNA library and sgRNA 1 was confirmed by 10% TBE-urea denaturing electrophoresis, as indicated by 100-nt bands corresponding to the size of our synthetic control sgRNA (**Fig. 1b**). We also noted the presence of 200-nt bands, indicating high molecular weight (HMW) dsRNA impurities from T7 RNAP RNA-templating activity (24). Overall, the 18-plex sgRNAs and sgRNA 1 show no differences between gel lanes, indicating successful synthesis of pooled and individual sgRNAs with our custom IVT protocol.

Next, we evaluated the feasibility of our complete sgRNA library synthesis workflow for generating libraries with hundreds to thousands of unique spacers. To reduce the cost of ordering microarray-derived oligos for large-scale production, we subpooled the oligo libraries (**Fig. 1a**). We obtained a single pool of ssDNA oligos containing 11,640 unique sgRNA spacers from Twist Bioscience. Each subpool primer pair was designed to amplify 206 to 2,626 spacer sequences corresponding to each sgRNA library.

We amplified each library using methods described in prior literature (25–27). Subpool amplification involved using quantitative PCR (qPCR) to determine the optimal number of cycles needed to reach the plateau phase while avoiding overamplification. For each qPCR and subsequent PCR amplification, we used 0.1 nanograms (ng) of diluted OLS pool. We then bulk-amplified each subpooled library using PCR, applying 0.01 ng of each subpool per reaction. We then used these amplified subpool fragments for the remainder of our sgRNA library synthesis workflow (**Fig. 1a**). To compensate for variable GGA product yields, we initially added a PCR and size-selection step following assembly to concentrate the transcription template for IVT. This additional step was performed before IVT to facilitate the production of large-scale sgRNA libraries and to re-transcribe the 18-plex sgRNA library for increased yield.

Following IVT, we evaluated the quality metrics of the sgRNA libraries to identify potential issues that could impact their downstream utility. We performed RNA-seq on all 10 microarray-derived sgRNA libraries and the re-transcribed 18-plex sgRNA library, based on a previous approach (28). Since the 5’ region of the sgRNAs contains the variable 20-nt spacer sequence, we employed 5’ Rapid Amplification of cDNA Ends (RACE) combined with template-switching to capture spacer diversity. The template-switched products were reverse-transcribed into cDNA, amplified, and nanopore sequenced, with a target of 1.5 million reads per sgRNA library, though the actual depth varied. The custom primer also incorporated unique molecular identifiers (UMIs) during cDNA amplification to account for potential PCR bias, errors, and facilitate consensus calling.

The percentage of expected spacers present in each sgRNA library (percent coverage) was 100% for the pilot 18-plex sgRNA library prepared from column-derived oligos. In comparison, percent coverage for microarray-derived sgRNA libraries ranged from 100% for the 227-plex library to 36% for the 939-plex library, with an average of 80% across all 10 libraries (SD ± 20%). A moderate negative correlation (Pearson correlation: *r* = −0.5787; **Table S1**) was observed between percent coverage and library scale for the microarray-derived libraries (**Fig. 1c**), indicating that coverage decreased as library scale increased. We also observed a strong positive correlation between coverage and nanopore sequencing depth (Pearson correlation: *R*² = 0.74, *r* = 0.87; **Fig. S1a**, **Table S1**), suggesting that sequencing depth strongly influences observed coverage. This finding indicates a non-uniform distribution within the library, as greater sequencing depth increases the likelihood of detecting low-abundance spacers in the tail of the distribution. Since nearly all microarray-derived libraries exhibited spacer coverage below 100%, addressing this limitation was essential to generate well-represented sgRNA libraries (**Fig. 1c**).

We evaluated the Gini Coefficient to assess the uniformity of sgRNA library representation. The Gini Coefficient measures inequality in sequence representation, where a value of zero indicates perfect uniformity and a value of one reflects perfect inequality (27, 29). Lower Gini Coefficients indicate more uniform spacer representation, reducing biases that could affect the identification of targeted hits in CRISPR screens or other assays. The 10 microarray-derived libraries had an average Gini Coefficient of 0.90 (SD ± 0.06), with values ranging from 0.81 for the largest library (2,626-plex) to 0.97 for the 342-plex library (**Fig. S1b**). We observed no correlation between Gini Coefficient values and library scale (Pearson correlation: *R*² = 0.04, *r* = −0.29; **Fig. S1b**, **Table S1).** Instead, the consistently high Gini Coefficients observed across individual sgRNA libraries likely result from differences in sequence composition within each library, rather than being a systematic effect of library size.

Given the consistently high Gini Coefficient values observed, we sought to establish a baseline for comparison. We extracted publicly available yield data for 72 individually transcribed sgRNAs produced by New England Biolabs using the EnGen® sgRNA Synthesis Kit. The normalized yield distribution for these individually transcribed sgRNAs served as a proxy for the expected read abundance of individual spacers within sgRNA libraries and had a Gini Coefficient of 0.33 (**Fig. 1d**). The large disparity between the Gini Coefficient of individually transcribed sgRNAs (0.33) and the much higher values observed in our pooled libraries raises the question of whether this increased inequality is due to the multiplexed nature of transcription or other factors. Notably, the smallest microarray-derived library (206-plex) exhibited a 160% increase in the Gini Coefficient (0.86) compared to individually transcribed sgRNAs, suggesting that multiplex sgRNA synthesis introduces substantial biases in spacer representation (**Fig. 1d**).

To investigate this further, we examined the relationship between the representation of an individual spacer in a multiplexed reaction and its yield when transcribed independently. Specifically, we compared the representation of sgRNA 1 within the pilot 18-plex sgRNA library to the yields of both the entire library and the singleplex transcribed sgRNA 1 (**Fig. 1b,c**). The 18-plex library produced 1.7 micrograms of total sgRNA, whereas sgRNA 1 yielded 0.33 micrograms, a 5.2-fold reduction in yield in the non-multiplexed setting. RNA-seq analysis revealed that sgRNA 1 accounted for only 0.0038 of the total reads in the 18-plex library, a 15-fold reduction from the expected 0.055 under perfect uniformity (**Fig. S2**). The more pronounced 15-fold reduction in read abundance suggests that multiplex transcription amplifies biases in spacer distribution, likely due to competition among templates within the same reaction.

Next, we evaluated the percentage of spacers with expected (perfect) and mutant sequences. Mutations are primarily introduced during the oligo synthesis, PCR amplification (2.8 x 10^−7^ errors/nt), and T7 transcription (5×10^−5^ errors/nt) (30). The 18-plex pilot sgRNA library had a significantly lower median percent perfects (82%, SD ± 5%) compared to the microarray-derived sgRNA libraries, with median values ranging from 89% (SD ± 9%) to 95% (SD ± 3%) (**Fig. S3**). A strong negative correlation (Pearson correlation: *r* = −0.90) was observed between median percent perfects and spacer coverage (**Table S1**). A similar trend is also observed between median percent perfects and sequencing depth (Pearson correlation: *R*² = 0.58, *r* = −0.8; **Fig. S1c**, **Table S1**). This correlation arises because spacers were included in the analysis only if they had at least 100 reads (including both perfects and mutants) in the RNA-seq data, ensuring sufficient representation. When spacer distributions are highly skewed, this threshold disproportionately includes overrepresented spacers, which tend to have a higher proportion of perfect sequences. PCR bias favors the perfect sequences which dominate, further enriching their representation. As a result, greater inequality in the distribution leads to a higher median percent perfects.

Consequently, the 18-plex sgRNA library, with 100% coverage, shows lower median percent perfects due to this dynamic (**Fig. 1c**). From these results, we concluded that our proof-of-concept sgRNA synthesis workflow could produce libraries with a high percentage of perfect spacers (**Fig. S3**). However, the microarray-derived sgRNA libraries exhibited reduced coverage (**Fig. 1c**) due to severe spacer inequalities as indicated by the Gini Coefficient values close to one (**Figs. 1d** and **S1b**). These biases could lead to important functional hits being missed during CRISPR-Cas9 screens, affecting the accurate identification of functional gene targets. While oversampling during screening can improve statistical power (21), such biases may still compromise reproducibility and accuracy. Heo and colleagues (2024) demonstrated that reducing biases in the DNA used to generate sgRNA libraries decreased the number of cells required in a CRISPRi screen by 10- to 20-fold. These findings underscore the importance of improving library quality (21).

### Reducing PCR Cycles in Microarray Oligo Amplification Fails to Improve sgRNA Library Uniformity

There are three potential sources of the observed spacer non-uniformities: oligo synthesis, PCR bias, or T7 RNA polymerase transcription bias. We first investigated how reducing the number of PCR cycles used to subpool and amplify microarray-derived libraries affected sgRNA representation, as observed through lower Gini Coefficients. PCR amplification is known to introduce biases into DNA libraries as cycle numbers increase, resulting in uneven sequence representation. These biases arise because DNA polymerase preferentially amplifies GC-rich priming motifs, leading to disproportionate enrichment of these sequences (31–34).

To test this hypothesis, we optimized the PCR cycle number for two sgRNA libraries: a large (1,382-plex) library and a smaller library (389-plex). PCR cycle optimization involved using qPCR to determine the ideal input template DNA concentration and the minimum number of PCR cycles needed for both subpool and bulk amplifications. Based on these results, we amplified the microarray-derived oligo libraries using 1 ng and 10 ng of template DNA—representing 10-fold and 1000-fold increases, respectively, over our initial protocol. This adjustment reduced the number of PCR cycles from 53 to 22 for the 389-plex sgRNA library and from 54 to 25 for the 1,382-plex sgRNA library. We also removed the amplification step following GGA, as it doubled the number of PCR cycles in our workflow and increased the risk of introducing mutations into the spacers.

As a comparison, we also obtained 135-nt column-synthesized oligos (oPools) from Integrated DNA Technologies, which included both the spacer sequence and scaffold. These oPool libraries were duplexed using RSPE with a single PCR cycle, serving as controls to establish a baseline spacer distribution with minimal PCR-induced bias. By comparing the Gini Coefficients of the reduced-PCR microarray-derived sgRNA libraries to those of the oPool-derived libraries, we aimed to determine whether severe spacer inequalities arose from excessive PCR cycles or the IVT process itself.

After IVT of the sgRNA libraries, we performed RNA-seq and analyzed the reduced-PCR microarray-derived libraries and the oPool control libraries to evaluate spacer distribution and overall sgRNA quality (**Fig. 2**). First, we compared the percent coverage of expected spacers across the reduced-PCR (22, 24, and 25 cycles) and excessive-PCR (53 and 54 cycles) microarray-derived libraries, as well as the oPool libraries amplified with a single PCR cycle (**Fig. 2a**). The 389-plex oPool library exhibited 80% target coverage (n = 1), while the 1,382-plex oPool library, transcribed in duplicate, exhibited 80% and 85% coverage (n = 2). The reduced-PCR microarray-derived libraries showed higher coverage, with two independently prepared 389-plex libraries demonstrating 97% and 100% coverage, and the 1,382-plex library reaching 94% coverage (**Fig. 2a**). In contrast, the excessive-PCR microarray-derived libraries exhibited substantially lower coverage, with only 65% for the 389-plex library and 84% for the 1,382-plex library (**Figs. 1c and 2a**).

**Figure. 2.**
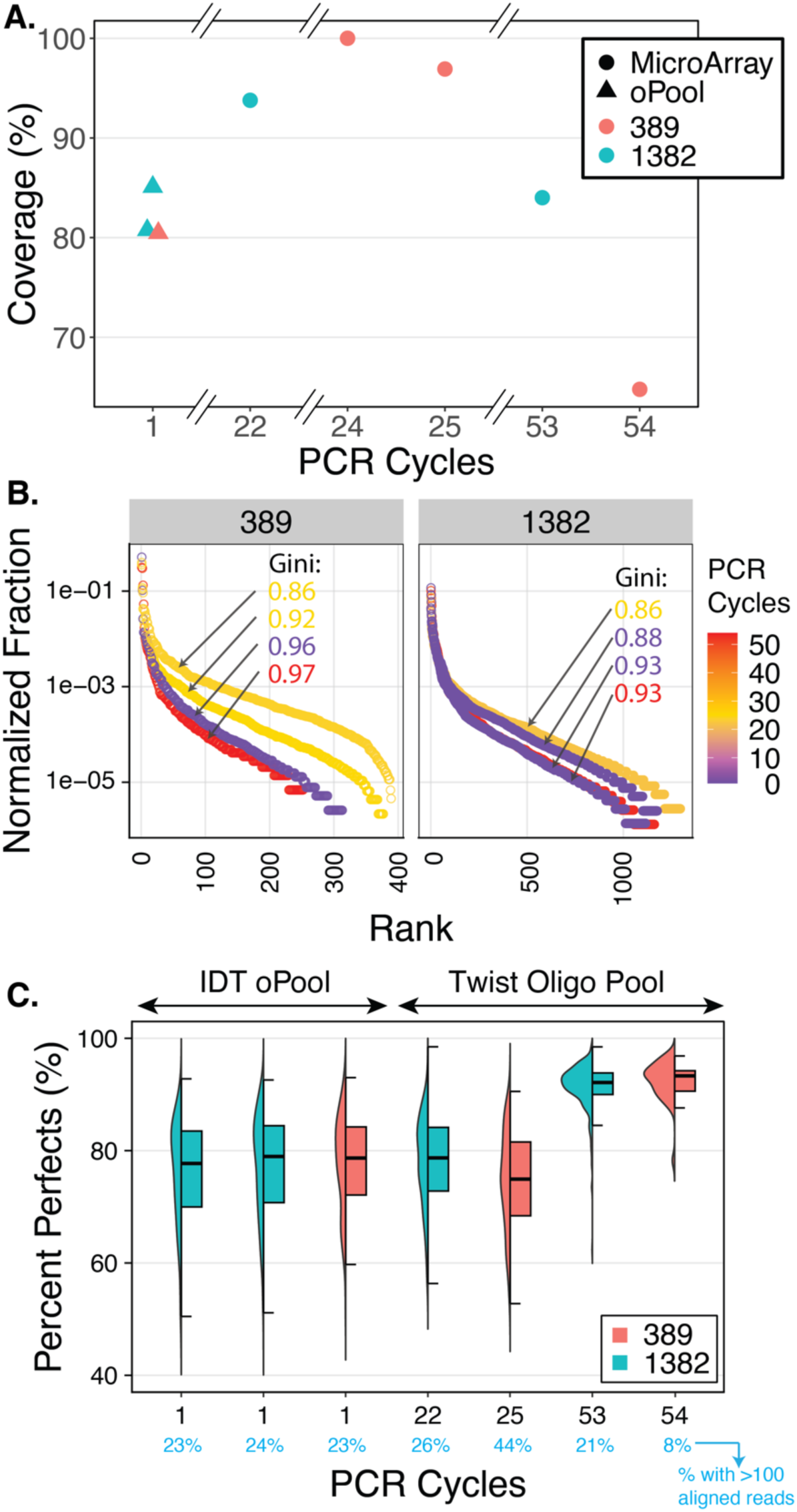
Comparison of sgRNA library metrics of two scales (389 and 1,382) with differing oligo sources and PCR cycles. **a.** Percent coverage of spacers across 389-plex (n = 4, orange) and 1,382-plex sgRNA libraries (n = 4, teal), prepared using microarray-derived oligos (Twist oligo pool; circles) or column-synthesized oligos (IDT oPool; triangles), as a function of the number of PCR amplification cycles. The x-axis describing the number of PCR cycles is broken between 1–22, 22–24, and 25–53**. b.** Normalized abundance (reads per spacer, relative to total reads) for 389-plex (n = 4) and 1,382-plex (n = 4) sgRNA libraries. Spacers are ranked in descending order of abundance, with PCR cycle numbers represented by a heatmap gradient, with purple corresponding to 0 cycles and red to 50 cycles. Gini Coefficients (Gini), displayed alongside each library’s rank-ordered curve, indicate the degree of inequality in spacer representation across different library scales and oligo sources**. c.** Percent perfects (proportion of perfectly synthesized spacer sequences) for 389-plex (orange) and 1,382-plex (teal) sgRNA libraries. Data for column-synthesized oligos (IDT oPool) and microarray-derived oligos (Twist oligo pool) are shown. Perfect spacers were only included if they had at least 100 reads in the RNA-seq data, ensuring sufficient representation for reliable analysis. The percentage of spacers with at least 100 reads is listed for each library in the x-axis (blue text). Each library is depicted using bifurcated plots: a half violin plot on the left (distribution of percent perfects) and a boxplot on the right (median percent perfect per library).

Initially, we expected oPool sgRNA libraries amplified with a single PCR cycle to achieve high coverage. We assumed that minimizing exponential PCR amplification would yield more uniform spacer representation, similar to microarray-derived libraries with low PCR cycles. However, the reduced-PCR microarray-derived libraries exhibited higher spacer coverage than the column-synthesized oPools (**Fig. 2a**). Despite this difference, spacer read counts were strongly correlated between the oPool and reduced-PCR microarray-derived 389- and 1,382-plex libraries (Pearson correlation: *R*² = 0.77, *r* = 0.99; *R*² = 0.68, *r* = 0.86; **Fig. S4a,b**). The strong correlation between spacer reads in two 1,382-plex oPool replicates (Pearson correlation: *R*² = 0.89, *r* = 0.79; **Fig. S4e**) supports the similarity in spacer distributions between oPool and reduced-PCR libraries. These results suggest that, despite differences in overall coverage, the relative spacer distributions remain highly similar, with variation in coverage likely arising from factors other than the oligos themselves.

We also expected the reduced-PCR libraries to achieve higher spacer coverage than the excessive-PCR libraries, and our results confirm this prediction. In the microarray-derived 389-plex sgRNA library, decreasing PCR cycles increased coverage by 32% and 35% (**Fig. 2a**). This increase in coverage is reflected in the moderate rather than strong correlation between spacer reads for the reduced- and excessive-PCR libraries (Pearson correlation: *R*² = 0.44, *r* = 0.32; **Fig. S4c**). A similar trend was observed in the 1,382-plex library, where increasing PCR cycles resulted in a 10% decrease in coverage (**Fig. 2a**). Due to the smaller magnitude of this effect, the correlation between spacer reads for the reduced- and excessive-PCR libraries remained stronger (Pearson correlation: *R*² = 0.72, *r* = 0.8; **Fig. S4d**). These results demonstrate that reducing PCR cycles effectively improves spacer coverage across two differently sized microarray-derived sgRNA libraries. We concluded that excessive PCR cycles reduce coverage due to inequality introduced by Kapa HiFi DNA polymerase bias (34).

Next, we evaluated the Gini Coefficients for the 389- and 1,382-plex sgRNA libraries, generated from DNA oligo libraries amplified with 1–54 PCR cycles. In the 1,382-plex libraries, the Gini Coefficient was 0.93 for the excessive-PCR library, 0.86 for the reduced-PCR library, and 0.93 and 0.88 for the column-derived oPool libraries (**Fig. 2b**). The 389-plex library followed a similar trend: the excessive-PCR library had a Gini Coefficient of 0.97, the reduced-PCR libraries yielded values of 0.92 and 0.86, and the oPool library showed a Gini of 0.96 (**Fig. 2b**). Overall, neither the reduced-PCR nor oPool libraries exhibited a consistent reduction in the Gini Coefficients compared to the excessive-PCR condition, suggesting that factors beyond PCR cycle number are the primary contributors to library uniformity.

In addition to percent coverage and Gini Coefficients, we evaluated the percentage of spacers with perfect sequences (percent perfects). We once again observed the trend of libraries with high inequality exhibiting higher percentage perfects. Low-PCR sgRNA libraries (1–25 PCR cycles) exhibited lower median percent perfects (mean: 78%, SD ± 1.7%, n = 5) compared to the 389- and 1,382-plex sgRNA libraries amplified with 53 and 54 PCR cycles, respectively (mean: 93%, SD ± 0.8%, n = 2; **Fig. 2c**). These observations suggest that DNA polymerase preferentially amplifies spacer sequences that are already overrepresented in the starting oligo libraries following excessive PCR cycles, while T7 RNA polymerase further enhances their abundance. These cumulative effects increase the representation of perfect spacers relative to less abundant mutant spacers or rare perfect spacers. While PCR, under ideal conditions, maintains the relative ratios of unique templates (in this case, spacer sequences) under ideal conditions, existing heterogeneity in the template pool may be exaggerated with increasing PCR cycles. This occurs due to competition between templates for limited PCR reagents such as primers and dNTPs (35). Excessive PCR cycling disproportionately amplifies high-abundance spacers while causing low-abundance spacers, including mutants, to drop below sequencing depth. Consequently, perfect spacers are enriched while overall spacer diversity decreases.

Pearson correlation analysis of the excessive-PCR microarray-derived sgRNA libraries further supports this observation. As mentioned previously, there is a strong negative correlation between the median percentage of perfect spacers and sequencing depth across all 10 libraries (Pearson correlation: *R*² = 0.58, *r* = −0.8; **Fig. S1c**). This indicates that perfect spacers are more abundant and more likely to be captured when sequencing depth is low, while more mutant spacers are included when sequencing depth is higher. This aligns with the strong positive correlation between spacer coverage and sequencing depth, showing that deeper sequencing captures both low-abundance perfect and mutant spacers (Peason’s correlation: *R*² = 0.74, *r* = −0.87; **Fig. S1a**, **Table S1**). Finally, our data directly support this prediction. In both the 389- and 1,382-plex sgRNA libraries, the fraction of reads for mutant spacers decreased with increasing PCR cycle number, while reads for target spacers increased (**Fig. S5**). Based on these findings, we recommend minimizing PCR cycles during oligo preparation to improve spacer coverage, consistent with prior reports (35).

Having established the effect of PCR cycle number on percent coverage (**Fig. 2a**) and percent perfects (**Fig. 2c**), we turned our attention to improving the Gini Coefficients. Our sgRNA synthesis workflow contains two steps mediated by polymerases: PCR with DNA polymerase and IVT with T7 RNAP (**Fig. 1a**). In addition to DNA polymerase, other enzymes, such as DNase and Tn5 transposase, compromise molecular genomics workflows due to their sequence preferences (36–38). Given our results we reasoned that T7 RNAP was the primary source of sequence bias in the sgRNA libraries, which would explain the persistence of spacer distribution bias following PCR cycle optimization (**Fig. 2b**). Since the sgRNA quality metrics were evaluated by reverse transcribing the sgRNAs into cDNA, it is possible that any reduction in the Gini Coefficient at the DNA level was masked by the subsequent T7 RNAP bias. This masking could occur because IVT is the final step in our workflow and would strongly influence the final Gini Coefficients obtained for each sgRNA library. On the other hand, improvements to the percent coverage of spacers were only due to the reduction in PCR cycles.

### T7 RNA Polymerase Favors Spacers Starting with Four Guanines

To identify spacers preferentially transcribed by T7 RNAP and understand their impact on spacer distribution, we analyzed position-dependent biases in sgRNA library spacer sequences produced by our workflow. Specifically, we assessed the log₂ fold change (FC) in the representation of each nucleotide at the first 10 positions of the 5′ end of all 20-nt spacers in each sgRNA library. We compared observed-to-expected ratios for each nucleotide at every position, visualizing the results as heatmaps for each library. The expected ratios were calculated based on the frequency of each nucleotide at each position in the designed spacer sequences, assuming a uniform distribution. Deviations from this baseline were captured by the log₂ fold change metric, as determined by RNA-seq data.

We first applied this analysis to the 389- and 1,382-plex oPool, PCR-reduced, and excessive-PCR sgRNA libraries (excluding duplicates) shown in **Fig. 2**, as these displayed pronounced spacer biases even after PCR optimization. The heatmap of the 389-plex oPool sgRNA library showed a mean log_2_ fold guanine enrichment of 2.1 within the first four nucleotide positions (mean: 2.1, SD ± 0.26). Conversely, adenines (mean: −1.89, SD ± 0.56), cytosines (mean: −2.25, SD ± 1.40), and thymines (mean: −2.97, SD ± 0.60) exhibited a log_2_ fold depletion of approximately 2.0 at these positions (**Figs. 3a** and **S6a**). Similar biases appeared in the reduced-PCR 389-plex library (**Fig. S6b**). However, in the excessive-PCR 389-plex library, guanine enrichment was limited to the first three positions (**Fig. S6c**), likely due to low spacer coverage (65%) caused by excessive PCR cycles (**Fig. 2a**). The 1,382-plex libraries showed similar but less pronounced bias trends (**Fig. S6d-f**), which may reflect library-to-library variation driven by unique spacer sequence compositions.

**Figure. 3.**
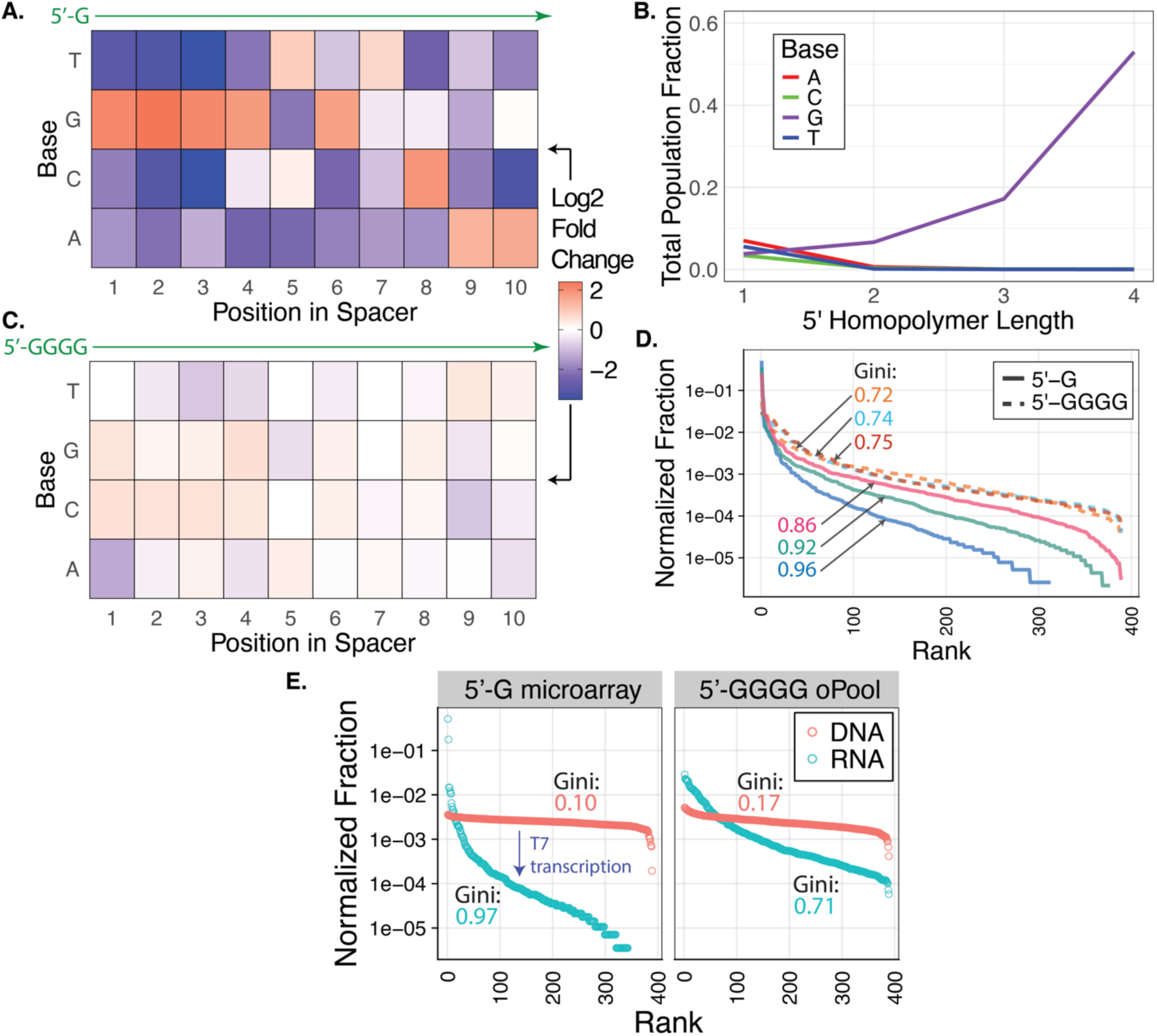
Influence of base composition on spacer abundance for 389-plex sgRNA libraries. **a.** Log₂ fold change of observed vs. expected spacer abundance for each nucleotide at the first 10 positions of 20-nt spacers in a 389-plex sgRNA library transcribed from oPool oligos. Observed values are based on spacer frequencies from RNA-seq data, while expected values assume uniform spacer distribution. Positive log₂ fold changes (orange) indicate enrichment, negative values (blue) indicate depletion, and white indicates no change. **b.** Fraction of spacers containing homopolymer stretches within the first four nucleotides at the 5’ end, analyzed from the same RNA-seq dataset as in **a. c.** Log₂ fold change of spacer abundance as in A, but for 389 spacers with a 5’ guanine tetramer added at positions +1 to +4 downstream of the T7 promoter. Nucleotide positions 1-10 along the spacer are immediately downstream of the 5′ tetramer. The same color scale applies as in A. **d.** Normalized fraction of spacer reads relative to total reads for 389 spacers with a single guanine at the first spacer position (solid lines, n = 3) versus 389 spacers padded with 5′ guanine tetramers (dashed lines, n = 3). Spacers are ranked by descending abundance. Each library, transcribed *in vitro* from 400 ng of template DNA in a 20 μL reaction volume, is represented by a distinct color. Rank-order curves with listed Gini Coefficients (Gini) quantify the inequality in spacer representation across library types. **e.** Gini Coefficients for 135 bp DNA libraries (coral) containing 389 spacers and their corresponding sgRNA libraries (teal), *in vitro* transcribed by T7 RNAP using 100 ng of input DNA in a 100 µL reaction. DNA libraries were generated from microarray-derived oligos with spacers starting with a 5’ guanine (left) or oPool-derived oligos with spacers padded with a 5’ guanine tetramer (right). Spacer distributions for each library format are shown as in **d**.

To investigate the guanine enrichment, we tested whether it was driven by the overall nucleotide composition of the first four positions or the presence of 5’ guanine homopolymers. In the 389-plex oPool sgRNA library, spacers containing 5’ homopolymers were enriched exclusively when containing 2- to 4-nt guanine homopolymer runs in the first four positions. In contrast, adenine, thymine, and cytosine 5’ homopolymer containing spacers were never enriched. Notably, spacers containing 2-, 3-, and 4-nt guanine homopolymers, represented 6.6%, 17%, and 53% of the total reads after transcription (**Fig. 3b**). Strikingly, spacers with 5′ guanine tetramers (5′ GGGG) were enriched by 3.1-fold compared to those with guanine triplets (5′ GGG) (**Fig. 3b**). These findings suggest that T7 RNAP strongly prefers transcribing sequences with guanine tetramers at the 5′ end, corresponding to positions +1 to +4 of the T7 promoter.

A prior study identified 5′ guanine triplets (5′ GGG) as a key determinant of T7 RNAP transcription efficiency. Using a library of randomized T7 promoter variants with an initiating guanine at position +1, they found that sequences with guanines at positions +2 and +3 were transcribed robustly. By randomizing positions +4 to +8 while keeping guanines fixed at +1 to +3, they demonstrated that guanine triplets enhanced transcription efficiency. This effect was observed regardless of the downstream sequence, likely due to the role of guanines in stabilizing the transcription bubble (39). However, the study did not assess the effect of guanines beyond position +3 or the impact of guanine tetramers on transcription efficiency.

In contrast, our study expanded the analysis to the first 10 nucleotides of sgRNA library spacers, providing a more comprehensive view of T7 RNAP’s sequence preferences. The 2.7-fold increased representation of 5′ guanine tetramers compared to triplets in the sgRNA libraries suggests that the additional guanine at position +4 enhances transcription efficiency (**Fig. 3b**). Given that a 5′ guanine triplet stabilizes the transcription bubble and prevents abortive initiation, we propose that the fourth guanine reinforces this effect, facilitating the transition from transcriptional initiation to elongation. These findings extend the model proposed by Conrad and colleagues (2020) and provide new insights into T7 RNAP behavior during sgRNA synthesis.

Our data confirm that T7 RNAP introduces extreme spacer bias within sgRNA libraries. Consequently, we recommend RNA-sequencing all sgRNA libraries prior to their use in CRISPR-Cas9 studies. To our knowledge, no studies have directly assessed spacer distribution within sgRNA libraries before their use in CRISPR applications. Typically, DNA libraries used as transcription templates and downstream CRISPR outputs, such as gene enrichment or phenotypic changes, serve as proxies for evaluating sgRNA library quality (15, 40, 41). Incorporating RNA-seq for direct quality control could improve accuracy, identifying biases early and saving time and resources during CRISPR screen optimization.

A common strategy for improving CRISPR-Cas9 screen reliability is redundancy, using multiple sgRNAs per gene and transduction replicates to enhance signal robustness (7). This approach helps mitigate variability in spacer activity and off-target effects, while also compensating for experimental noise and the limitations of prediction tools. However, unrecognized biases in sgRNA library composition, like those introduced by T7 RNAP, could further distort representation and impact screen performance. Incorporating RNA-seq for direct quality control would allow researchers to identify underrepresented spacers early, ensuring higher-quality libraries and potentially optimizing redundancy. Rather than using excessive sgRNA numbers to overcome unknown biases, well-characterized libraries improve redundancy, enhancing target coverage and streamlining experimental design and resource use. This approach would improve both the robustness and practicality of CRISPR screens and related assays.

### Reducing sgRNA Spacer Bias by Adding 5′ Guanine Tetramers

Given the strong bias of T7 RNAP for spacers with guanines in the first four positions (**Fig. 3a,b**), we hypothesized that adding a 5′ guanine tetramer to all spacers would reduce this bias. This modification could stabilize the T7 RNAP initiation-to-elongation transition and improve transcription of spacers otherwise disfavored when preceded by a single guanine. However, this approach raised some considerations. The four guanine homopolymer, corresponding to the first four positions after the TSS of the T7 promoter, could introduce 5′ sequence heterogeneity in sgRNAs, potentially complicating the production of single-species RNA for structural studies like NMR or X-ray crystallography. Nonetheless, correctly sized species can be isolated through gel electrophoresis or size-selection methods (42). Additionally, guanine tetramer padding might interfere with sgRNA spacer formation of R-loop complexes after Cas9 binding, a critical step in CRISPR-Cas9 target recognition. However, functionality tests in a cell-free system demonstrated a 2-fold higher enrichment of 12 DNA targets using a 12-plex sgRNA library containing a 5’ guanine tetramer complexed with dCas9 compared to using a 12-plex sgRNA library containing a single 5’ guanine (43). Thus, we concluded that the benefits of improved spacer uniformity outweigh potential 5′ heterogeneity, with no observed deficiencies in CRISPR-dCas9 binding or target recognition.

To test our hypothesis, we redesigned the 389-plex spacer library by adding a guanine tetramer to the 5′ end of each spacer. To evaluate whether the effects of this modification were dependent on scale, we grouped the 389 spacers into 12-, 60-, and 389-spacer subpools. As a control for the 12-plex 5′ GGGG subpool, we designed a second version of the 12-plex library by adding a single 5′ guanine (5′ G) only to spacers starting with A, C, or T. These were ordered as a single 473-oligo oPool from Integrated DNA Technologies. qPCR optimization showed that 10 ng of oPool input and no more than 12 PCR cycles yielded robust amplification across all subpools. Unlike microarray-derived oligos, which require separate amplification steps, a single subpool PCR generated sufficient DNA for GGA to append the sgRNA scaffold, producing 135-bp products for IVT.

Our first objective was to assess whether adding the 5′ guanine tetramer to spacers improves sgRNA library uniformity. To this end, we performed IVT with the GGA-produced DNA library containing 12 spacers padded with a 5′ GGGG sequence (library 12G4) or a single 5′ guanine (5′G, 12G). Resulting sgRNA libraries were reverse transcribed and nanopore sequenced using the 5′ RACE protocol. When comparing the spacer distributions between the two libraries, depicted as the normalized fraction of reads per spacer out of total reads, both showed similar trends. The Gini Coefficient values calculated from the spacer read distributions were 0.61 for 12G and 0.74 for 12G4 (**Fig. S7a**), suggesting slightly increased inequality in the 12G4 library. To further evaluate whether this sequence modification influenced spacer biases, we conducted Pearson correlation analysis of spacer reads between the 12G and 12G4 libraries. A lack of correlation would indicate that the modification exaggerated or reduced biases. Instead, we observed a strong positive correlation (*R*² = 0.76, *r* = 0.962; **Fig. S7b**), indicating that the relative abundance of spacers remained consistent across both libraries.

From these data, we inferred that adding a 5′ guanine tetramer did not significantly affect spacer biases at small scales. These findings prompted us to investigate whether this trend holds in larger sgRNA libraries containing 389 spacers padded with a 5′ guanine tetramer. To explore this, we performed IVT to generate an oPool-derived sgRNA library containing 389 spacers with a 5′ GGGG sequence (library 389G4). The resulting sgRNA library was sequenced as before and we determined the log₂ FC of observed vs. expected spacer abundance for each nucleotide within the first 10 positions of the 389G4 spacers.

The heatmap of the 389G4 sgRNA library showed a mean log_2_ fold enrichment of 0.40 for guanines within the first four nucleotide positions (SD ± 0.15), reflecting a 5.3-fold decrease compared to the 389G library at these positions (mean log_2_ fold change: 2.11, SD ± 0.25). Additionally, the depletion of adenines, cytosines, and thymines at these positions was less pronounced than in the 389G library (**Fig. 3a, c**). Pearson correlation analysis between 389-spacer microarray-derived sgRNA libraries with a single 5′ G and oPool-derived libraries padded with a 5′ GGGG showed no correlation in read counts (*R*² = 0.04, *r* = 0.05; **Fig. S8a**). A replicate of this comparison also showed no correlation (*R*² = 0.18, *r* = 0.14; **Fig. S8c**). Similarly, no correlation was observed between oPool-derived 5′ GGGG spacers and oPool-derived 5′ G spacers (*R*² = 0.02, *r* = 0; **Fig. S8b**). Analysis of multiple 389-plex sgRNA libraries showed lower Gini Coefficients for 5′ GGGG spacers (0.72, 0.74, 0.75; n = 3) compared to 5′ G spacers (0.86, 0.92, 0.96; n = 3). Overall, the 5′ GGGG libraries showed a mean 19.3% decrease in the Gini Coefficient (SD ± 2.81%) (**Fig. 3d**). We also evaluated the Gini Coefficients of 389-spacer sgRNA libraries transcribed with modified IVT conditions, using 100 ng input DNA in a scaled-up 100 μL reaction instead of the standard IVT setup, which used 400 ng input DNA in a 20 μL reaction. This adjustment produced a similar trend: a mean 22.7% decrease (SD ± 7.74%) in the Gini Coefficient for 5′ GGGG-padded spacers (n = 3) compared to spacers starting with a single guanine (n = 1) (**Fig. S9**). Combined, these results indicate that guanine tetramer padding substantially improved spacer representation, reducing inequality across the library while maintaining a more consistent and expected spacer distribution.

To determine whether the 5′ GGGG spacer modification affected overall sgRNA library yield, we analyzed the yields of multiple sgRNA libraries. A Wilcoxon rank-sum test revealed no significant difference in yield between libraries with 5′ GGGG spacers (n = 8, median: 0.805 μg) and those with 5′ G spacers (n = 13, median: 1.29 μg, p = 0.5002) (**Fig. S10**). Although there was a slight trend toward higher yields in the 5′ G libraries, the difference was not statistically significant. These results demonstrate that the 5’ GGGG spacer modification does not affect the overall sgRNA yield.

### Effect of Oligo Type on DNA Library Quality for IVT

We assessed whether oligo type influenced DNA library quality by sequencing the DNA templates and comparing microarray-derived oligos with 5′ G spacers and oPool-derived oligos with 5′ GGGG spacers. The percentage of perfect spacers was similar between the 389-plex microarray-derived library (median: 96.30%, n = 1) and the 389-plex oPool-derived library (median: 95.73%, n = 1). The distribution of percent-perfect spacers was also comparable between the two libraries, as shown in the violin plot in **Fig. S11a**. These results contrast with earlier reports of higher error rates for microarray-derived oligos compared to column-derived oligos (44, 45). Notably, the microarray-derived 5′ G library underwent 24 PCR cycles, while the oPool-derived oligos were amplified with only 7 cycles. All PCR amplification occurred during the subpool and bulk amplification stages, with no additional PCR performed after GGA of oligos to the conserved sgRNA scaffold. Despite this higher cycle count, both libraries exhibited similarly high percentages of perfect spacers (**Fig. S11a**). Given the vendor-reported error rate for microarray oligo synthesis (one error per 3,000 nt), it is likely that microarray technology now produces oligos with error rates comparable to column-derived oligos. Furthermore, while Mighell and colleagues (2022) recommend using column-derived oligos due to their lower reported synthesis error rates, our results support the use of modern microarray-derived oligos as a high-quality and cost-effective alternative.

We also compared spacer inequality between libraries by analyzing the Gini Coefficients of normalized read fractions. The oPool-derived 389-plex 5′ GGGG DNA library showed a more skewed spacer distribution (Gini = 0.17, n = 1) compared to the microarray-derived 5′ G library (Gini = 0.10, n = 1), as determined by the Wilcoxon rank-sum test (p = 3.97 × 10⁻⁸; **Fig. S11b**). Furthermore, there was no correlation between the two libraries (Pearson correlation: *R*² = 0, *r* = 0.04; **Fig. S11c**). The observed inequalities could be due to the 5′ GGGG modification, DNA polymerase bias, or the oligo synthesis method. Yet, despite undergoing only 7 PCR cycles compared to the 24 PCR cycles for the 5’G library, the 5′ GGGG library exhibited greater skew, which could be caused by variable amplification efficiencies or PCR polymerase bias associated with the homopolymer region. Still, both libraries maintained high rates of spacer percent perfects rates and good overall quality, reinforcing that the primary source of inequality in sgRNA libraries arises from T7 RNAP sequence preferences (**Fig. 3e**).

We also examined the impact of oligo source (microarray vs. oPool) on the distribution of transcribed sgRNA libraries. Neither the 5′ GGGG nor 5′ G DNA library spacer reads correlated with those of their transcribed sgRNA libraries (**Fig. S12a,b**). This indicates that T7 RNAP biases are present in both library formats and contribute to skew. More specifically, the microarray-derived 5’G sgRNA library had a Gini Coefficient of 0.97 (n = 1), an 89.7% increase from the Gini of the 5’G DNA library that encodes it. In contrast, the oPool-derived 5′ GGGG sgRNA library yielded a 13.6% lower Gini coefficient of 0.71 (n = 1), reflecting a 76.1% increase from its corresponding DNA library (**Fig. 3d**).

Notably, this difference stems from T7 RNAP sequence preferences, as the difference in Gini coefficients between the 5’ G and 5’ GGGG sgRNA libraries is 3.7-fold larger than the difference observed between their DNA libraries (**Fig. 3d**). These findings suggest that the type of oligo used to construct the DNA library had less influence on spacer distribution than the 5′ guanine modification’s impact on the resulting sgRNA libraries. Together, these data demonstrate that padding the 389 spacers with a 5′ guanine tetramer effectively reduced T7 RNAP transcription biases, though this effect was not observed at the smaller, 12-spacer scale.

Both microarray-derived and oPool-derived oligos demonstrated excellent quality, with fewer than 5% mutant spacers and minimal spacer inequality. Though the oPool-derived library with 5′ GGGG spacers showed slightly greater spacer distribution skew than the 5′ G microarray-derived library, the difference was small but statistically significant. These inequalities were decoupled from the inequalities that we have observed after transcription due to the behavior of T7 RNAP. While we initially used oPool-derived oligos to quickly optimize libraries and IVT conditions with small pools, our experimental design offers a more cost-effective strategy.

### Emulsion IVT (eIVT) Enhances sgRNA Library Uniformity

Designing sgRNA spacers with a 5′ GGGG sequence substantially improved sgRNA library uniformity. However, this approach still has substantially more inequality than the Gini (0.33) observed for the 72 sgRNAs individually transcribed by NEB (**Fig. 1d**). We hypothesized that modifying the IVT process itself could further improve library uniformity, either independently or in combination with the 5′ GGGG spacer design. To this end, we developed an emulsion-based IVT (eIVT) workflow that isolates sgRNA template DNA molecules within droplet microcompartments, reducing competition for IVT reagents and T7 RNAP. This strategy draws from the successful use of oil-water emulsions in DropSynth gene synthesis, where compartmentalization prevents cross-hybridization of DNA oligos and reduces competition during both DNA assembly and amplification (25, 27). A similar approach is used in whole-genome amplification (WGA) for single-cell sequencing, where emulsions minimize competition among genome fragments during amplification, reducing bias (46). Given these effects, we predicted that eIVT would yield sgRNA libraries with lower Gini Coefficients compared to bulk IVT. To accommodate our existing emulsification protocol, we increased IVT reagent concentrations fivefold while varying DNA input, scaling a standard 20 μL IVT reaction up to 100 μL, which is the aqueous phase volume used for emulsification.

The aqueous phase was combined with commercial fluorinated oil and emulsified by vortexing (25). To assess whether compartmentalization in emulsions slowed transcription, we prepared two 12-plex sgRNA libraries by incubating eIVT reactions at 37°C for either 2 hours, which is the standard IVT time, or 16 hours. The 2-hour incubation condition showed a 2-fold increase in yield compared to the 16-hour incubation condition [data not shown], confirming that the transcription process was complete within 2 hours within emulsions. Additionally, IVT incubation time did not affect spacer distribution for both conditions (**Fig. S13a**), with nearly identical normalized read fractions and a strong positive one-to-one linear relationship (Pearson correlation: *R*² = 0.99, *r* = 0.99; **Fig. S13b**). Based on these results, all subsequent eIVT experiments used a 2-hour incubation at 37°C.

After incubation, we broke the emulsions with chloroform, treated the aqueous phase with DNase, and purified the sgRNAs via column purification (**Fig. 4a**). Since emulsions can function as compartmentalized bioreactors on a microscale, we evaluated whether eIVT affects the total yield of sgRNA libraries compared to bulk IVT. We also assessed how library scale and spacer sequence modifications (5’ G vs. 5’ GGGG) influence IVT performance under both conditions. Bulk IVT controls were prepared identically to the aqueous phase of each eIVT, using the same DNA input and 100 μL reaction volume. By maintaining equivalent volumes, we controlled for any potential effect of reaction volume on sgRNA yield and spacer uniformity. A Wilcoxon rank-sum test showed no significant difference (p > 0.05) in sgRNA yields between libraries transcribed with 100 ng of input DNA using bulk IVT (median: 1.8 μg, SD = ± 0.98, n = 14) and those transcribed with eIVT (median: 1.8 μg, SD = ± 0.98 n = 13; **Fig. 4b**). These results indicate that eIVT is a viable method for sgRNA transcription, as it maintains comparable yields to bulk IVT.

**Figure. 4.**
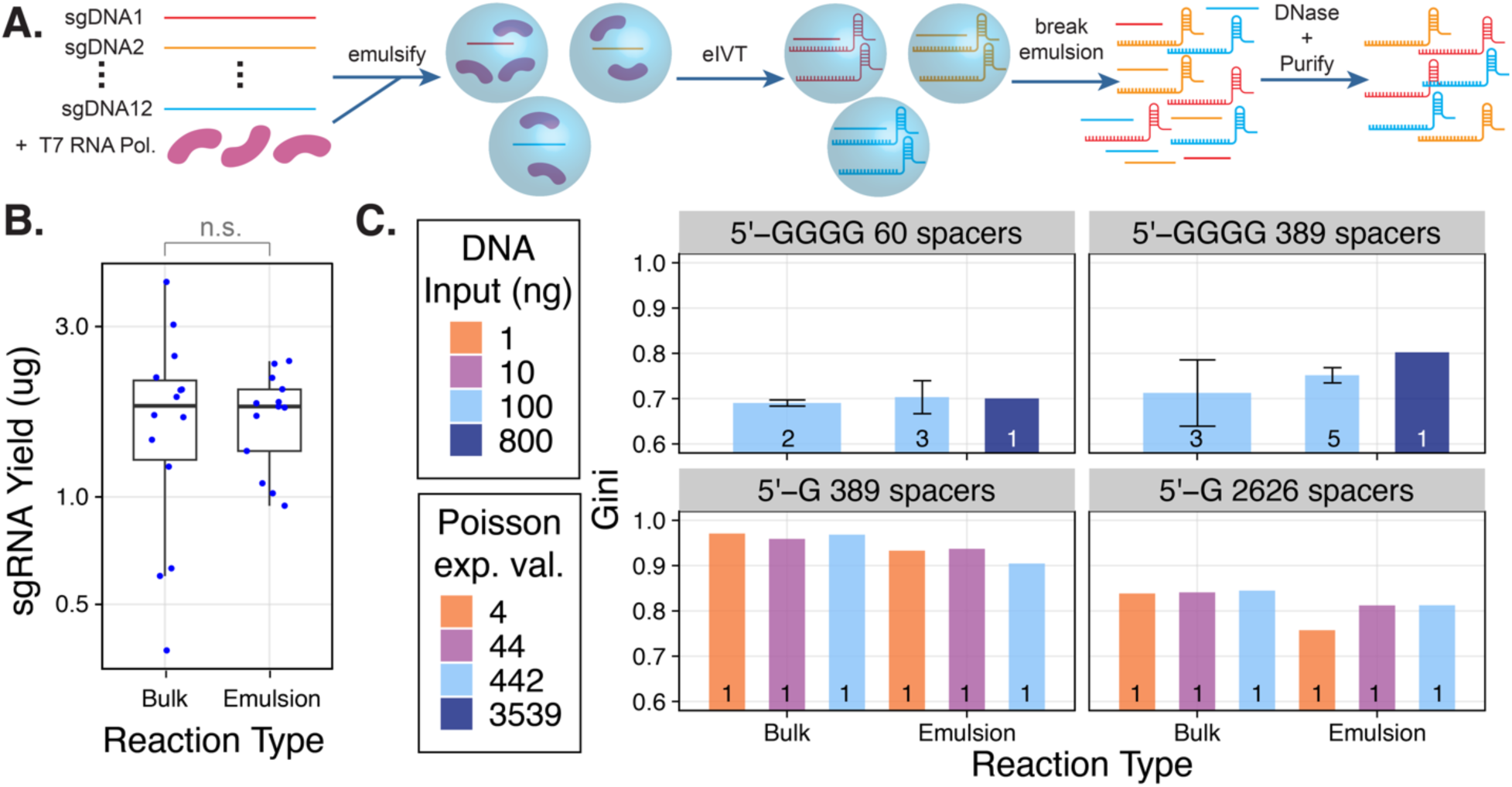
Evaluating *in vitro* transcription in emulsions (eIVT) as a novel approach to synthesize sgRNA libraries. **a**. A 100 μL IVT reaction was prepared by scaling up a standard 20 μL IVT reaction fivefold and adding Golden Gate-assembled library DNA templates to the aqueous phase. The aqueous phase was emulsified in a fluorinated oil by vortexing, generating stable droplets that function as microreactors for IVT. After transcription, emulsions were broken, and sgRNA library products were purified via DNase treatment and column purification. **b.** Comparison of sgRNA yields for various sgRNA library scales transcribed with 100 ng of input DNA using bulk IVT (n = 14) or eIVT (n = 13). sgRNA libraries generated by bulk IVT serve as controls for each sgRNA library produced by eIVT. Yields (μg) are further categorized by library scale: 12-plex (n = 5), 18-plex (n = 1), 60-plex (n = 5), 389-plex (n = 12), and 2,626-plex (n = 3). A Wilcoxon rank-sum test found no significant difference in sgRNA yields between bulk IVT and eIVT (p > 0.05). **c.** Libraries contained spacers starting with either a single 5’ guanine (5’ G) or a 5’ guanine tetramer (5’ GGGG). The top panel displays the median Gini coefficients for 5’ GGGG sgRNA libraries synthesized using bulk eIVT versus bulk IVT with 800 ng (navy, Poisson = 3,539) or 100 ng (aqua, Poisson = 442) of input DNA. The Poisson value represents the estimated mean number of input DNA molecules per emulsion droplet. Individual bars represent each sgRNA library type, with sample sizes labeled. Bar heights indicate the median Gini coefficient per category, and black bars represent standard deviation (SD). The bottom panel shows median Gini coefficients for 5’ G sgRNA libraries synthesized using bulk eIVT versus bulk IVT with 100 ng (aqua, Poisson = 442), 10 ng (purple, Poisson = 44), or 1 ng (orange, Poisson = 4) of input DNA. Sample size is one for all sgRNA libraries in the bottom panel.

Previously, we observed that adding a 5’ GGGG sequence to a 389-plex sgRNA library reduced the Gini Coefficient by a mean of 19.3%, improving uniformity compared to the equivalent library with 1G spacers (**Fig. 3d**). We hypothesized that eIVT of a library DNA template with the same 5’ GGGG modification would further enhance spacer uniformity, lowering Gini Coefficient values even more. To test this, we transcribed 60- and 389-plex sgRNA libraries with 5’ GGGG spacers (60G4 and 389G4) using eIVT and bulk IVT with either 100 ng or 800 ng of input DNA. These DNA amounts were chosen to assess the impact of DNA input on eIVT, given that DNA partitioning within emulsion droplets follows a Poisson distribution. To maximize compartmentalization efficiency, reducing this mean toward one molecule per droplet is ideal, but too little DNA may compromise sgRNA library yield for downstream applications. Therefore, we compared 800 ng (Poisson = 3,539) and 100 ng (Poisson = 442) DNA inputs. At these high DNA inputs, multiple molecules will co-localize within droplets. For instance, in emulsions with a 5 µm mode droplet diameter (25), 500 ng of 135 bp DNA yields a Poisson mean of 942 DNA molecules per droplet.

Following RNA-seq analysis of transcribed sgRNA libraries, the 60G4 sgRNA library had median Gini Coefficients of 0.69 (SD ± 0.0068, n = 2) for 100 ng bulk IVTs, 0.70 (SD ± 0.036, n = 3) for 100 ng eIVTs, and 0.70 (n = 1) for the 800 ng eIVT. The 389G4 sgRNA library had median Gini Coefficients of 0.71 (SD ± 0.073, n = 3) for 100 ng bulk IVTs, 0.75 (SD ± 0.017, n = 3) for 100 ng eIVTs, and 0.80 (n = 1) for the 800 ng eIVT. No bulk IVT controls were included for the 800-ng input condition. Across all comparisons, eIVT did not reduce median Gini Coefficients relative to bulk IVT sgRNA libraries (**Fig. 4c**, upper panel). These findings suggest that a Poisson distribution of 442 DNA molecules per droplet (from 100 ng input DNA) is too high for effective template partitioning, potentially allowing intermolecular competition for reagents.

To further investigate the effects of Poisson loading in emulsions on library uniformity, we tested input DNA amounts of 100 ng (Poisson 442), 10 ng (Poisson 44), and 1 ng (Poisson 4) while performing eIVT of 389- and 2,626-plex 5’ G sgRNA libraries. The 2,626-plex library was included to assess whether eIVT offers greater benefits at larger library scales, where competition among spacers for reagents is expected to be more pronounced. No sgRNA libraries transcribed with 1 ng of input DNA, nor the two bulk IVTs with 10 ng of input DNA, yielded detectable RNA by fluorometer quantification. However, we performed 5’ RACE-RT and successfully sequenced all sgRNA libraries using nanopore sequencing, regardless of yield. Each sgRNA library condition was transcribed once (n = 1). To assess statistical significance, we applied the Kolmogorov–Smirnov (KS) test to determine whether the spacer distributions of eIVT and bulk IVT sgRNA libraries were drawn from the same population at a given scale. While the KS test evaluates whether observed differences arise by chance or reflect a real effect, the Gini Coefficient indicates the directionality of changes in distribution uniformity.

The 389-plex sgRNA library produced with eIVT and 1 ng of input DNA exhibited a significantly lower Gini Coefficient (0.93) compared to the bulk IVT control (0.97) (**Fig. 4c**). This difference was statistically significant according to the KS test (D = 0.20352, p = 3.53 × 10⁻⁷; Table S2). To assess whether position-dependent biases of T7 RNAP persisted following eIVT, we evaluated the Log₂ FC for each nucleotide within the first 10 positions at the 5′ end of all 20-nt spacers. The eIVT library showed a global reduction in overall nucleotide bias compared to the bulk library, with FC values closer to the expected baseline of zero (**Fig. S14a,b**). This indicates that the strong bias toward guanine within the first four positions was reduced in the eIVT library relative to the bulk IVT control.

This bias reduction was even more pronounced in the 2,626-plex library, where the Gini coefficient decreased from 0.84 (bulk IVT) to 0.76 (eIVT) (**Fig. 4c**). The KS test confirmed the statistical significance of this result (D = 0.22022, p = 2.20 × 10⁻¹⁶; **Table S2**). Additionally, the Log₂ FC heatmap showed a global decrease in nucleotide bias, with a distinct pattern of enrichment for guanine at position 2 and cytosine at position 5 (**Fig. S15a,b**). This pattern likely reflects library-to-library variation driven by unique spacer sequence compositions, as previously observed in the 1,382-plex library (**Fig. S6d-f**).

With 10 ng of input DNA, the 389-plex eIVT library did not exhibit a lower Gini Coefficient (0.94) compared to the bulk IVT control (0.96) (**Fig. 4c**), as indicated by the lack of statistical significance (KS test: D = 0.0977, p = 0.054; **Table S2**). However, the FC heatmap showed a global reduction in bias (**Fig. S14c,d**). In contrast, the 2,626-plex eIVT library exhibited a significantly lower Gini Coefficient (0.81) than the bulk IVT control (0.84) (**Fig. 4c**). The KS test confirmed significance (D = 0.0984, p = 7.35 × 10⁻¹⁰; **Table S2**), accompanied by a consistent reduction in positional bias (**Fig. S15c,d**).

With 100 ng of input DNA, the 389-plex eIVT library exhibited a lower Gini Coefficient (0.90) than the bulk IVT control (0.97) (**Fig. 4c**). The KS test confirmed statistical significance (D = 0.3626, p = 2.20 × 10⁻¹⁶; **Table S2**), and the FC heatmap indicated a global reduction in nucleotide bias (**Fig. S14e,f**). Similarly, the 2,626-plex eIVT library exhibited a significantly lower Gini Coefficient (0.81) than the bulk IVT control (0.84) (**Fig. 4c**), with the KS test confirming significance (D = 0.0811, p = 3.19 × 10⁻⁷; **Table S2**). The FC heatmap reflected decreased guanine overrepresentation and a broader recovery of A, T, and C levels toward the expected log₂ FC values of zero. (**Fig. S15e,f**).

Together, these results show that for both the 389-plex and 2,626-plex libraries, the eIVT method consistently reduced Gini Coefficients and improved uniformity at 1 ng and 100 ng DNA input amounts. At 10 ng input, the 389-plex library showed a decrease in Gini Coefficient under eIVT, though not statistically significant, while the 2,626-plex library showed a significant reduction. These findings suggest a possible scale-dependent effect.

We next examined how spacer representation differs between eIVT and bulk IVT. To assess whether varying the DNA input reduces guanine-driven bias at the 5’ end of spacers, we analyzed guanine representation at the first five positions. The percent decrease in guanine representation served as a simple proxy for bias in spacer distribution. For each sgRNA library transcribed with 1 ng, 10 ng, or 100 ng input DNA, this value was calculated as the change in Log₂ FC between bulk IVT and eIVT, normalized by the bulk IVT Log₂ FC. This approach was based on our previous finding that guanines within the first four positions drive overrepresentation (**Fig. 3**).

For the 389-plex library, the percent decrease in guanine representation at spacer positions 1 to 4 remains at or below 25% across all DNA input amounts. At position 5, guanine representation decreases by 48.2%, 23.6%, and 47.6% for the 1 ng, 10 ng, and 100 ng DNA input conditions, respectively. Notably, the 1 ng and 100 ng conditions roughly twice the reduction seen at 10-ng (23.6%) (**Fig. S16a**), consistent with the lack of a significant Gini Coefficient decrease at the intermediate input (**Fig. 4c**, **Table S2**). For the 2,626-plex sgRNA libraries, the percent decrease in guanine representation at positions 1 to 3 remained below 86.3% across all DNA input amounts. At position 4, reductions were more pronounced: 113% for the 100-ng input condition, 166% for the 10-ng input, and 253% for the 1 ng input. In contrast, at position 5, guanine representation increased slightly for the 1 ng (25.8%) and 10 ng (19.8%) inputs, while the 100-ng input showed a modest 16.7%, decrease (**Fig. S17a**).

These results show that eIVT effectively reduces the overrepresentation of guanine-containing spacers at nucleotide positions previously identified as biased toward guanine. In the 389-plex library, the greatest relative reduction in bias occurred at position 5, immediately following positions 1 to 4 (**Fig. S16a**). In contrast, the 2,626-plex library exhibited a stronger bias toward guanine-containing spacers at positions 2 and 4, with the most substantial reduction at position 4 (**Fig. S17a**). Across all DNA input amounts, percent decreases in guanine bias were consistently lower for the 389-plex library compared to the 2,626-plex library (**Figs. S16a** and **S17a**). For the 389-plex library, the 1 ng and 100 ng eIVT conditions produced the most notable reduction in bias, while in the 2,626-plex library, the largest decrease occurred at 1 ng, supporting the prediction that lower DNA input enhances effective compartmentalization. As expected, the 10 ng and 100 ng conditions showed more modest reductions (**Fig. S17a**). The smaller scale of the 389-plex library may have contributed to variability in guanine bias reduction due to stochastic effects. Finally, this analysis focuses exclusively on changes in guanine bias and does not capture broader nucleotide composition bias.

Given the narrow scope of this analysis, we evaluated overall changes in spacer representation for bulk IVT and eIVT sgRNA libraries using Pearson correlation analysis. Correlation linear fits were compared to a unity line, which represents a perfect 1:1 relationship between spacer read fractions in each condition. Steeper linear fit slopes indicate greater reductions in bias using emulsion compartmentalization. Red dashed lines denote the median read fraction per spacer from the input DNA library transcription template, reflecting a highly uniform spacer distribution (Gini = 0.102; **Fig. S11b**). Spacers that decreased in representation due to eIVT appear as blue dots above the unity line, while those that increased appear as red dots.

The 389-plex sgRNA libraries exhibited strong positive correlations across 1 ng (*R*² = 0.9, *r* = 0.95), 10 ng (*R*² = 0.85, *r* = 0.99), and 100 ng (*R*² = 0.79, *r* = 0.98) DNA input amounts (**Fig. S16b-d).** With eIVT, overabundant spacers decreased in representation for the 1 ng and 10 ng input conditions compared to bulk IVT (**Fig. S16b,c**). In contrast, the 100-ng condition showed an increase in the representation of low-abundance spacers (**Fig. S16d**). Notably, the two most overrepresented spacers decreased in the 10 ng and 100 ng input conditions, whereas only one of these spacers showed a reduction in the 1 ng condition (**Fig. S16b-d**). Since these changes are plotted on a log_10_ scale, reductions in the most overrepresented spacers are substantial. The 2,626-plex sgRNA libraries also showed positive correlations across 1 ng (*R*² = 0.66, *r* = 0.61), 10 ng (*R*² = 0.76, *r* = 0.91), and 100 ng *(R*² = 0.76, *r* = 0.92) DNA input conditions. However, differences between individual input amounts were less pronounced. The most overrepresented spacer decreased in the 1 ng and 10 ng conditions but remained largely unchanged in the 100-ng condition (**Fig. S17b-d**).

Overall, these findings suggest that reducing the most overrepresented spacers has the greatest impact on uniformity metrics. The more pronounced effects observed in the smaller 389-plex library likely reflect stochastic variation, whereas the larger 2,626-plex library exhibits more consistent behavior across conditions. Paradoxically, the pronounced stochastic effects in the smaller 389-plex library offer valuable insights into how DNA input amount influences transcription efficiency in emulsions. Across both libraries, the lower Gini Coefficients and greater reduction in guanine overrepresentation support our hypothesis that the 1 ng DNA input condition (Poisson 4) enhances compartmentalization and uniformity. However, despite these promising proof-of-concept results, the 1 ng condition did not yield a quantifiable sgRNA output, limiting its practical utility for library generation. In contrast, the 100-ng input condition (Poisson 442) produced the most substantial reduction in guanine bias and a significantly lower Gini Coefficient for the 389-plex library. This suggests that higher DNA template concentrations enhance T7 RNAP transcription efficiency and improve uniformity by increasing substrate availability. The 10-ng input condition (Poisson 44) showed moderate effectiveness in reducing T7 RNAP bias in both libraries but did not significantly lower the Gini Coefficient for the 389-plex library. This may indicate that the 10-ng input condition fails to strike a balance between sufficient DNA concentration for transcription efficiency and the compartmentalization benefits observed at lower inputs.

Given the limited sample size and the application of the KS test across all conditions, our ability to fully resolve the relationship between DNA input and Gini Coefficients may be restricted. Nonetheless, these findings show that eIVT significantly reduces bias in the spacer distributions of sgRNA libraries with 389-plex or greater complexity. Further optimization will be required to achieve comparable or greater bias reduction for 5’ GGGG libraries relative to bulk IVT protocols.

### Evaluation of High Molecular Weight Byproducts in sgRNA Libraries

We investigated the presence of double-stranded RNA (dsRNA) byproducts in sgRNA libraries to better understand why eIVT improves spacer uniformity in libraries with 5’ G spacers but not in those modified with a 5’ GGGG sequence. We hypothesized that the 5’ GGGG motif may interfere with the uniformity gains observed in eIVT for libraries containing 5’ G spacers. To explore this, we performed polyacrylamide gel electrophoresis of 389-plex sgRNA libraries transcribed under various DNA input conditions using either eIVT or bulk IVT (**Fig. S18**). Only libraries with quantifiable yields were analyzed. The gel revealed that all sgRNA libraries contained the expected 100-nt sgRNA products, confirmed by band alignment with a synthetic sgRNA control from IDT (**Fig. S18a**). However, most libraries also showed a prominent high molecular weight (HMW) 200-nt dsRNA byproduct, likely generated by the residual RNA-dependent activity of T7 RNAP (47). We did not analyze these byproducts beyond this point since we cannot capture information about these dsRNA byproducts in our RNA-seq data. The residual RNA-dependent activity of T7 RNAP causes cis-primed extension from small hairpin loops within transcribed sgRNAs, producing looped-back dsRNA products (48, 49). These dsRNA impurities are known to trigger innate immune responses when IVT-produced RNA is used in therapeutic applications (50–52). As a result, they are often removed by high-performance liquid chromatography, cellulose-based chromatography, or native purification methods (53–55). The formation of these byproducts is associated with high-yield conditions during T7 RNAP IVT batch synthesis (28, 49), so their presence in both eIVT and IVT libraries is unsurprising.

Unexpectedly, the eIVT condition prepared with 10 ng of input DNA produced minimal dsRNA byproducts, as observed in the gel image (**Fig. S18a**). ImageJ quantification of band intensities revealed that this condition showed almost no detectable dsRNA signal, with levels even lower than the synthetic sgRNA control, while still producing clear sgRNA products. In contrast, the corresponding IVT reaction with 10 ng of input DNA displayed low band intensities for both sgRNA and dsRNA products (**Fig. S18b**). Analysis of relative dsRNA band intensity confirmed that most of the total RNA signal in the eIVT condition corresponded to sgRNA products, whereas the IVT condition showed a higher proportion of dsRNA byproducts (**Fig. S18c**). These results suggest that emulsions combined with a low DNA input of 10 ng may help prevent the formation of dsRNA byproducts by promoting low-yield rather than high-yield conditions. However, we could not directly compare this observation to band intensities from eIVT prepared with 1 ng of input DNA due to insufficient yield for analysis by gel electrophoresis. These promising results warrant further replication to confirm whether eIVT with 10 ng or less input DNA enhances sgRNA production while minimizing dsRNA byproducts. If validated, this approach could offer a valuable strategy for optimizing sgRNA synthesis conditions.

In addition to the 200-nt dsRNA byproducts, the gel also revealed that all sgRNA libraries containing spacers starting with 5’ GGGG had multiple HMW RNA bands longer than 300 nt (**Fig. 18Sa**). Padding spacers with a 5’ GGGG sequence offers advantages, such as improved spacer uniformity compared to 5’ G sgRNA libraries when both are transcribed in bulk IVT (**Figs. 3d and 4d**). However, this approach also produces additional HMW RNA species, which are a clear drawback. Although we do not observe a statistically significant difference in overall RNA yield between 5’ G and 5’ GGGG sgRNA libraries (**Fig. S10**), a higher proportion of the quantified RNA from the 5’ GGGG libraries consists of non-functional HMW RNAs rather than functional sgRNAs. These excessive HMW byproducts appear regardless of whether the libraries were transcribed using eIVT or standard IVT (**Fig.18Sa**). Furthermore, since we first began transcribing them, we have consistently observed greater levels of HMW byproducts in 5’ GGGG libraries compared to 5’ G libraries (**Fig. S18d**). Thus, when the 5’ guanine tetramer is added, the functional sgRNA species are a smaller relative fraction of the total RNA population.

Despite the presence of these HMW products, the 5’ GGGG libraries remain suitable for *in vitro* CRISPR applications (43). Moreover, these byproducts can be reduced or eliminated through purification methods and the use of engineered T7 RNAP variants (52, 56), making this approach likewise potentially viable for producing high-quality messenger RNA libraries for mammalian cells or *in vivo* applications.

### Future Directions for Optimizing eIVT

Comparing the NEB dataset on individually transcribed sgRNAs to our eIVT results (**Fig. 4c**) suggests substantial room for further optimization within our compartmentalization approach. In our current method, we were only able to reduce the Poisson expectation value to 4, indicating that multiple spacers were still co-transcribed within individual droplet compartments. One potential strategy to overcome this limitation is using barcoded beads to isolate spacers into individual microreactions, similar to the DropSynth gene synthesis method (56, 57). In this approach, beads decorated with unique barcodes corresponding to short barcodes on the 5′ end of DNA oligos could selectively capture DNA templates encoding single spacers. By scaling the number of beads with the library size, this method could improve spacer representation and reduce bias. For example, 1,536 unique beads could transcribe a 1,536-plex sgRNA library. Concentrating individual spacer sequences within droplets would likely enhance library uniformity and scalability, making this a promising direction for future development. Moreover, using barcoded beads ensures a high local concentration of template DNA in each droplet, which should avoid the reduced yields observed when Poisson loading was minimized without bead capture.

Another potential improvement involves reducing droplet size without decreasing input DNA amounts. Our current eIVT emulsions, generated by vortexing, produce droplets with a typical diameter of 5 µm, which is relatively large compared to the DNA template input. Reducing the diameter to 700 nm while maintaining 100 ng of input DNA would yield a Poisson expectation value of 0.76, substantially decreasing the probability of multiple spacers co-localizing within a single droplet. This reduction in droplet size could improve spacer uniformity while preserving sufficient sgRNA yield for CRISPR applications. Although uniform droplets of this size can be produced using microfluidic methods (57, 58), they require specialized chips and equipment, increasing operational complexity. Similarly, generating very small vesicles to reduce the Poisson loading is technically challenging and may offer limited additional benefits. In contrast, a bead-based barcoding approach would be relatively straightforward to implement, making it the most practical option for optimizing eIVT.

### Effects of IVT Reaction Conditions on sgRNA Library Uniformity

Due to optimizations made to IVT reactions for compatibility with an emulsions-based transcription approach, we investigated how varying DNA input and total IVT reaction volume affect sgRNA library uniformity in the absence of emulsions. To do this, we transcribed sgRNA libraries using 100 μL and 20 μL reaction volumes with varying DNA input amounts. The 100 μL reaction was included as it matched the aqueous phase conditions used in eIVT. Following RNA-seq, we assessed the Gini Coefficients of sgRNA libraries across different scales (12-, 60-, 389-, and 2,626-plex) using DNA input amounts ranging from 1 to 400 ng and spacers beginning with either a 5’ G (1G) or 5’ GGGG (4G) (**Fig. 5**). Given the limited sample size, with most libraries represented by a single sample, we applied the KS test to determine whether spacer distributions deviated significantly from the input DNA population.

**Figure. 5.**
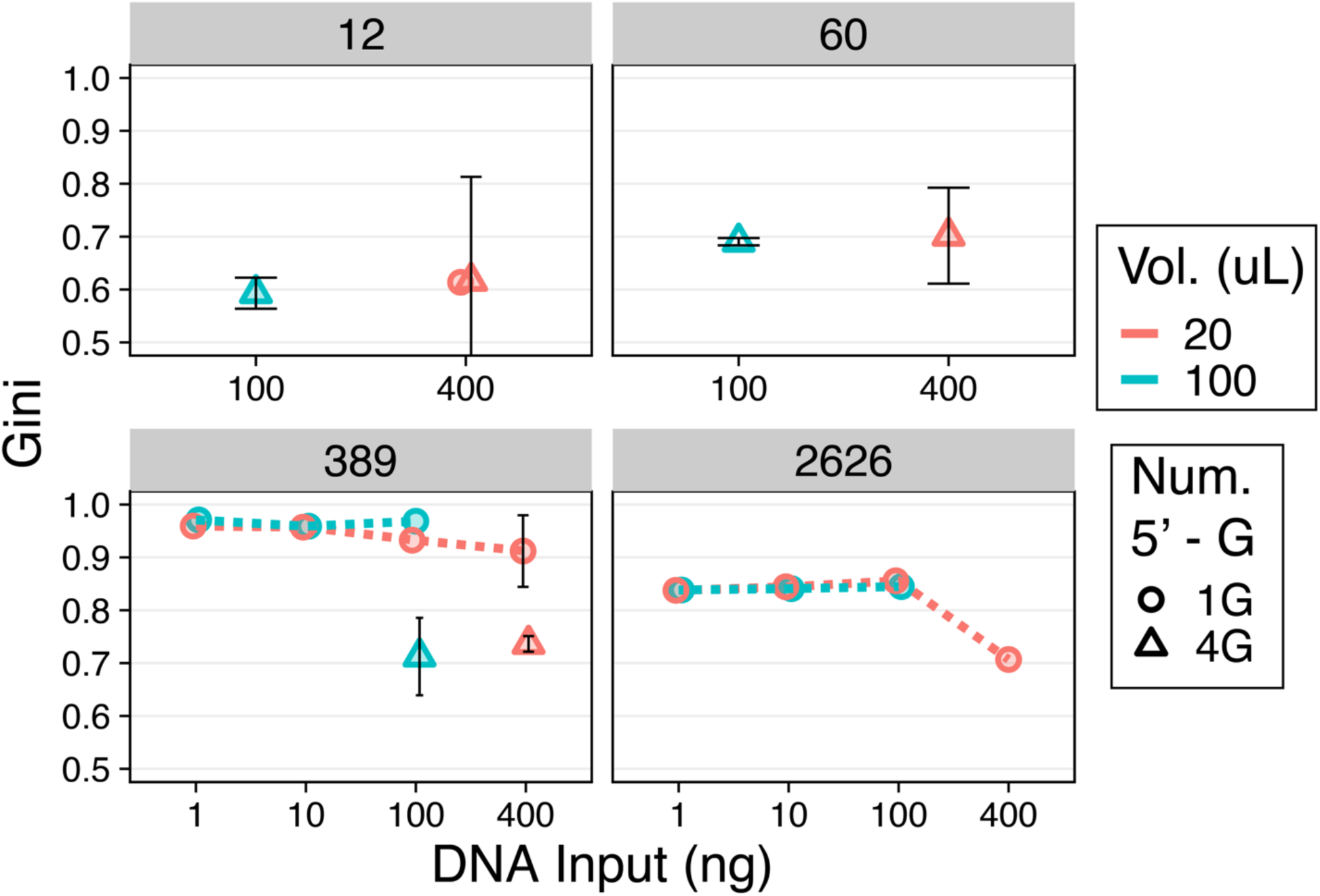
Gini Coefficients of sgRNA libraries across different scales, DNA input amounts, and reaction volumes. Libraries were transcribed in either a standard 20 μL reaction volume (orange) or a scaled-up 100 μL reaction volume (blue). Spacers were designed to start with either a single 5’ guanine (1G, shown as open circles) or four 5’ guanines (4G, shown as open triangles), except for the 2,626-plex library, which only contains 1G spacers. Libraries were generated at four different scales: 12-plex, 60-plex, 389-plex, and 2,626-plex, using DNA input amounts ranging from 1 ng to 400 ng. Most libraries were transcribed with a sample size of one. However, the following sgRNA libraries contain a sample size of two: 12-plex (4G, 100 ng, 100 μL), 60-plex (4G, 100 ng, 100 μL), 60-plex (4G, 400 ng, 20 μL), and 389-plex (1G, 400 ng, 20 μL). The following sgRNA libraries contain a sample size of three: 12-plex (4G, 400 ng, 20 μL), 389-plex (4G, 100 ng, 100 μL), and 389-plex (4G, 400 ng, 20 μL). For libraries with n > 2, the median Gini Coefficient is plotted with bars representing standard deviation.

For the 12- and 60-plex libraries, Gini Coefficients remained consistent across different reaction volumes and DNA input amounts. In the 12-plex libraries, the median Gini Coefficient for 4G spacers was 0.59 (SD ± 0.029, n = 2) for a 100 μL reaction with 100 ng DNA, compared to 0.62 (SD ± 0.20, n = 3) in 20 μL reactions with 400 ng DNA. Similarly, the 60-plex libraries maintained Gini Coefficients between 0.69 to 0.70 across all tested conditions. Variability was more pronounced in libraries transcribed with 400 ng DNA in 20 μL reactions (**Fig. 5**), though this was less evident in larger libraries, likely due to limited replication conditions.

In the 389-plex libraries, small yet statistically significant changes in Gini Coefficients were observed across DNA input conditions in 100 μL reactions (KS test, p < 0.05; **Table S3**). The 10-ng input showed a slight reduction in Gini (0.96) compared to the 1-ng and 100-ng inputs (both 0.97) (**Fig. 5**). More substantial changes were observed in 20 μL reactions, where increasing DNA input from 1 ng to 400 ng progressively and significantly reduced the Gini Coefficient from 0.97 to 0.91 (Fig. 5, KS test, p < 0.05, **Table S3**). This suggests that higher DNA input can improve library uniformity in smaller reaction volumes. The 4G version of the 389-plex library consistently outperformed the 1G version, with lower Gini values (0.71 to 0.74) across both reaction volumes (**Fig. 5**), consistent with earlier results (**Fig. 3**) demonstrating that the 5’ GGGG modification reduces T7 RNA polymerase transcriptional bias relative to 5’ G spacers.

For the 2,626-plex libraries, there was minor but statistically significant variation in Gini Coefficients across DNA input amounts in both 20 μL and 100 μL reactions (KS test, p < 0.05, **Table S3**). At this larger scale, the effect of DNA input on uniformity became more pronounced. In 100 µL reactions, libraries prepared with 1 ng and 100 ng DNA showed similar Gini Coefficients (approximately 0.84), while the 10-ng condition showed a slight but statistically significant improvement (Gini = 0.81) (KS test, p < 0.05, **Table S3**). For the 20 μL reactions, the trend was more clear: increasing DNA input from 1 ng to 400 ng progressively decreased the Gini Coefficient from 0.84 to 0.71, representing the best uniformity observed in this experiment. Notably, the change from 100 ng to 400 ng DNA in the 20 μL condition resulted in a substantial drop in Gini from 0.84 to 0.71 (**Fig. 5**).

Given that the largest improvement in the Gini Coefficient was observed when increasing the input DNA amount from 100 ng to 400 ng in a 20 µL reaction volume for the 2,626-plex 5’ G sgRNA libraries, we evaluated this condition more closely. Notably, the 400-ng library showed a clear reduction in nucleotide bias compared to the 100-ng library, with log_2_ fold decrease of 1.3 in guanine overrepresentation at position 2 and a broader recovery of A, T, and C levels toward baseline FC values (**Fig. 19a,b**). To further investigate whether increasing the DNA input from 100 ng to 400 ng reduces bias at the first five spacer positions which are most susceptible to bias, we examined the percent decrease in guanine representation. The percent decrease at positions 1 to 3 was at or below 53.6% and increased to 331% at position 4 (**Fig. S19c**). This represents a 78% reduction in guanine bias at position 4 compared to the 253% decrease in guanine representation observed for the 2,626-plex library produced by eIVT with 1 ng of input DNA (**Figs. S19c** and **S17a**). However, since in the baseline condition guanines are not as overrepresented at position 4 compared to positions 1 and 2, any decrease in guanine representation at this position results in larger percent decreases.

We hypothesize that the improvements in Gini Coefficient for libraries transcribed with 400 ng input DNA in 20 µL result from higher sgRNA template concentration. This likely increases T7 RNAP substrate availability, leading to more efficient and uniform transcription. While higher DNA input improves uniformity, the most consistent enhancement comes from using spacers that start with a 5′ GGGG sequence rather than 5′ G. This benefit was observed across various library sizes, including the 389-plex (**Figs. 3** and **5**), and likely applies to larger libraries as well. Therefore, we conclude that incorporating a 5′ GGGG sequence before spacer regions is the most effective strategy for improving sgRNA library uniformity, especially for libraries of 389 spacers or more.

When redesigning spacers with a 5′ GGGG sequence is not feasible, we recommend using 400 ng of input DNA instead of 100 ng for IVT of large-scale sgRNA libraries in a 20 µL reaction volume. Our findings may also inform future optimization of our eIVT method, as we observed that increasing DNA input to 100 ng reduced bias in both the 389- and 2,626-plex libraries. While we tested up to 800 ng of input DNA for smaller libraries (12- and 60-plex) without observing bias reduction, we did not test this approach at larger scales, where higher DNA input may still yield additional improvements in uniformity.

## Conclusions

Large-scale sgRNA libraries are essential tools in CRISPR-Cas9 research, enabling functional screens and a wide range of *in vitro* assays. However, there is a growing demand for user-defined, programmable sgRNA libraries that can be prepared using simple and cost-effective methods. In this work, we developed a workflow that meets this need while addressing a critical challenge often overlooked in *in vitro* transcription of heterogeneous templates: the spacer distribution biases introduced by T7 RNA polymerase. We found that T7 RNAP strongly favors spacers containing guanines in their first four positions, resulting in the systematic underrepresentation of other nucleotide sequences. This bias was further confirmed by comparing the severe spacer inequalities in our sgRNA libraries to the highly uniform DNA templates used for transcription.

The bias we identified in sgRNA libraries can compromise the accuracy and reproducibility of CRISPR-Cas9 screens by underrepresenting certain spacers, potentially leading to the loss of important functional hits. Beyond CRISPR screens, T7 RNAP biases may also affect methods that rely on *in vitro* transcribed RNA baits for hybridization to exons or ancient DNA in targeted sequencing, although this remains to be assessed (59, 60). Consequently, these biases could impact a broad range of CRISPR-based and RNA library-based studies.

In this study, we explored three independent strategies to mitigate T7 RNAP-driven bias and improve sgRNA library uniformity. The most effective was the addition of a 5’ guanine tetramer to all spacer sequences, which consistently and substantially reduced bias. This simple design stage modification has the potential to enhance the accuracy and reproducibility of CRISPR-Cas9 screens by enhancing the detection of functional hits that might otherwise be missed due to poor spacer representation. However, we also found that 5’GGGG modified templates generate substantially more high-molecular-weight (HMW) RNA byproducts than those starting with a single 5’G posing a key limitation to this approach.

In addition to spacer sequence redesign, we explored two alternative strategies to reduce transcription bias: compartmentalizing transcription within emulsion droplets and varying DNA input and reaction volumes during bulk *in vitro* transcription. Our results demonstrate that emulsions improve library uniformity in a scale-dependent manner, particularly at lower DNA inputs. Future optimizations, such as ligating input DNA templates to barcoded beads prior to emulsification, could enhance this effect. Notably, an sgRNA library transcribed in emulsions with a low DNA input amount (e.g., 10 ng) lacked HMW RNA byproducts, which are known to trigger inflammatory responses in human cell lines and *in vivo*. Although this requires further validation, it suggests that emulsion-based transcription could offer additional, unexpected benefits. We also found that higher DNA concentrations (20 ng/μL) significantly reduced bias compared to lower concentrations (5 ng/μL or below). When spacer redesign or emulsion-based transcription is not feasible, we recommend maximizing DNA input (400 ng) in small reaction volumes (20 μL) under standard IVT conditions to achieve optimal library uniformity.

Beyond addressing T7 RNAP biases, our results show that modern microarray-derived oligos have low mutation rates and comparable quality compared to column-derived oligos. This contrasts with earlier reports of poor microarray oligo fidelity and reflects improvements in synthesis technologies. By ordering a pool of 11,640 unique oligos and subpooling them via PCR into 10 sgRNA libraries of varying sizes, we achieved up to 72% cost savings compared to ordering individual libraries. Coupled with our *in vitro* transcription workflow, this approach eliminates the need for commercial kits requiring specific formats, making large-scale sgRNA library production more accessible and affordable for high-throughput studies.

Finally, this study opens avenues for future improvements in sgRNA design and synthesis. The principles used to mitigate T7 RNAP biases could be extended to guide RNA libraries for alternative Cas proteins, such as Cas12a and Cas13a. In particular, CRISPR-Cas12a guide RNA (crRNA) libraries may naturally exhibit greater spacer uniformity due to their sequence architecture where the conserved crRNA scaffold lies at the 5’ end downstream of the T7 promoter (15), unlike Cas9 sgRNAs, which place the variable spacer region at the 5’ end. By combining emulsion-based synthesis with careful reaction optimization, our approach represents an important step toward producing high-quality, uniformly distributed sgRNA libraries. We hope this work inspires new methodologies and enhances the precision and reliability of CRISPR-based applications.

## Supporting information

Supplementary Data 1 - microarray and oPool oligo sequences

## Acknowledgements

This work was supported by the National Science Foundation MCB-2032259 grant. Research reported in this publication was supported by the National Institute of General Medical Sciences of the National Institutes of Health under award number T32GM149387 (to NV). The content is solely the responsibility of the authors and does not necessarily represent the official views of the National Institutes of Health.

## Data and Materials Availability

Raw nanopore sequence reads for sgRNA libraries were submitted to the NCBI Sequence Read Archive under BioProject accession PRJNA1237881 (https://www.ncbi.nlm.nih.gov/bioproject/1237881).

Supplementary Data 1 contains all microarray and oPool oligo sequences. Supplementary Data 2 contains processed data sequences and counts can be found at https://doi.org/10.6084/m9.figshare.28635950

## Materials and Methods

### Sourcing and Pooling 18 Oligos for the Pilot sgRNA Library

Eighteen individual 98nt oligos were ordered from Integrated DNA Technologies (IDT, Coralville, USA) as single-stranded DNA (ssDNA) oligos. The oligo sequences are listed in **Table S4**. To pool the oligos, 2 µL of each 100 µM target oligo was combined to a total volume of 36 µL. Subsequently, at least 10 reverse single primer extension (RSPE) reactions were performed to convert the pooled target oligos into double-stranded DNA. Each RSPE reaction contained 1.25 µL of pooled target oligos, 2.5 µL of 100 µM reverse primer (skpp15-1-R filt15-1181), 12.5 µL of 2X Q5 Hot-Start High-Fidelity Master Mix (NEB), and nuclease-free water to a final volume of 25 µL. The thermocycler RSPE program consisted of an initial denaturation at 98°C for 30 sec, followed by one cycle of 98°C for 30 sec, 64°C for 30 sec, and 72°C for 60 min. DNA Cleanup Columns capable of binding five micrograms of DNA (NEB) were used to purify the double-stranded DNA oligonucleotides (dsDNA oligos).

### Preparing Microarray-Derived Oligos for Large-Scale sgRNA Synthesis

The pool of 11,640 microarray-derived oligos (chip9) that was resuspended to 4.3 ng/µL following manufacturer guidelines (Twist Bioscience, South San Francisco, USA). The 98 nt chip9 oligos included 10 unique forward and reverse primer pair sites for PCR subpooling, enabling the generation of templates for 10 sgRNA libraries.

### Subpool Amplification of 10 Microarray-Derived Libraries Prior to PCR Optimization

First, chip9 OLS were diluted 10-fold to a concentration of 0.43 ng/µL. Subpooling qPCRs were prepared using library-specific primer pairs and annealing temperatures as listed in **Table S5**. Each qPCR included 0.1 ng of diluted chip9 OLS, 1.25 µL each of 10 µM forward and reverse primers, 0.25 µL of 100X Biotium Thiazole Green (Thermo Fisher Scientific), 12.5 µL of 2X Kapa HiFi HotStart ReadyMix (Roche Sequencing Solutions Inc, Pleasanton, USA), and nuclease-free water to a total volume of 25 µL. The qPCR program included: initial denaturation at 98°C for 45 seconds, followed by 50 cycles of 98°C for 10-15 seconds, annealing for 15 seconds (see **Table S5** for temperatures), and extension at 72°C for 15 seconds. PCRs were prepared without Thiazole Green, and each subpool was amplified with the number of PCR cycles corresponding to the amplification plateau observed via qPCR (**Table S5**). The ten subpooled oligo libraries were purified using 5 µg DNA Cleanup Columns (NEB).

### Bulk Amplification Prior to PCR Optimization

Bulk amplification qPCR was prepared similarly to subpooling qPCR, except that 0.01 ng of subpooled DNA oligos was added per reaction. This adjustment was made to prevent the deckchair effect observed with larger template inputs. Additionally, the qPCR initial denaturation and denaturation times were reduced to 30 and 10 seconds, respectively. PCR was performed with the number of qPCR cycles corresponding to the amplification plateau for each subpool (**Table S5**), and a final 1-minute extension at 72°C. The ten bulk-amplified libraries were purified using 5 µg DNA Cleanup Columns (NEB).

### Subpool Amplification Optimization of 389-, 1,382-, and 2,626-Plex Microarray-Derived Libraries

To minimize the number of PCR cycles required for subpooling chip9 389-, 1,382-, and 2,626-plex libraries, qPCRs were performed with varying amounts of OLS template. For the 389-plex library, qPCR reactions included 0.1 ng, 0.25 ng, 0.5 ng, or 1 ng of 0.43 ng/µL chip9 OLS, along with 1.25 µL of 10 µM forward primer (skpp15-23-F filt15-577), 1.25 µL of 10 µM reverse primer (skpp15-23-R filt15-1596), 0.25 µL of 100X Biotium Thiazole Green (Thermo Fisher Scientific), 12.5 µL of 2X Kapa HiFi HotStart ReadyMix (Roche Sequencing Solutions Inc., Pleasanton, USA), and nuclease-free water to a final volume of 25 µL.

The qPCR program consisted of an initial denaturation at 98°C for 30 seconds, followed by 50 cycles of 98°C for 10–15 seconds, annealing at 53°C for 15 seconds (for the 389-plex subpool), and extension at 72°C for 15 seconds. The lowest cycle number was achieved using 1 ng of chip9 OLS, which was selected for subpooling the 389-plex library.

To determine the optimal amplification cycles for subpooling the 1,382-plex library, qPCR was performed with 1 ng of template using skpp15-6 primers and annealing at 53°C for 15 seconds (**Table S5**). The same approach was applied to the 2,626-plex library using the skpp15-27 primer pair with an annealing temperature of 51°C for 15 seconds. All three oligo libraries (389-, 1,382-, and 2,626-plex) were subpooled using the qPCR-determined cycle numbers (**Table S5**) with a 1-minute extension at 72°C. Subpooled oligos were purified using 5 µg DNA Cleanup Columns (NEB).

### Bulk Amplification Optimization of 389-, 1,382, and 2,626–Plex Microarray-Derived Libraries

To reduce PCR cycles for bulk amplification of chip9 389-, 1,382-, and 2,626-plex oligo libraries, qPCR was performed with 1 ng, 10 ng, or 20 ng of subpooled DNA and the same primers as used for subpooling. Amplification with 10 ng of template resulted in the least PCR cycles. PCR was performed with the number of qPCR cycles corresponding to the amplification plateau for each library (**Table S5**). A final 1-minute extension at 72°C was also added. All three bulk-amplified libraries were purified using 5 µg DNA Cleanup Columns (NEB).

### Column-Synthesized (oPool) DNA Library Templates Prepared with One PCR Cycle

Two column-synthesized oPools containing 120-nt target oligos with a T7 promoter, 20-nt spacer, and sgRNA scaffold sequence, were obtained from Integrated DNA Technologies (IDT, Coralville, USA). The oPools were resuspended in Tris-EDTA (TE) buffer, pH 8.0 according to the manufacturer’s guidelines. The S2 oPool contained oligos with 1,382 unique spacers from chip9, while S4 oPool contained oligos with 389 unique spacers from chip9.

S2 and S4 oPools were converted to dsDNA by performing RSPE as follows. Three reactions were prepared per oPool, each containing 2.5 µL of µM reverse primer (sgRNA_GGA_oligo_REV_NV; **Table S6**), 500 ng of S2 or S4 oligos, 12.5 µL of 2X Q5. The thermocycler RSPE program consisted of an initial denaturation at 98°C for 30 sec, followed by one cycle of 98°C for 10 sec, 70°C for 30 sec, and 72°C for 60 min. DNA Cleanup Columns capable of binding five micrograms of DNA (NEB) were used to purify the dsDNA oligos.

### Column-Synthesized (oPool) sgRNA library templates with 5’ Guanine Tetramer Spacers

One oPool (e13sgRNA) containing 473 oligos that were all 98 nt in length was obtained from Integrated DNA Technologies (IDT, Coralville, USA) and was resuspended in Tris-EDTA (TE) buffer, pH 8.0 according to the manufacturer’s guidelines. The oligos included four unique forward and reverse primer pair sites for PCR subpooling four target oligo libraries. Three subpools included a 5’ GGGG sequence at positions +1 to +4 after the T7 promoter, with 12 (12G4), 60 (60G4), or 389 (389G4) unique spacers. The fourth subpool had the same 12 spacers, but each started with 5’ G (12G) at the +1 position.

### Subpool Amplification of oPool Oligos

qPCR was performed with varying amounts of oPool template to optimize subpooling of the 12G and 12G4 libraries. Subpool-specific primer pairs and corresponding annealing temperatures are in **Table S7**. Each subpool qPCR included 0.1 ng, 1 ng, or 10 ng of oPool template, 1.25 µL of 10 µM forward and reverse primers, 0.25 µL of 100X Biotium Thiazole Green (Thermo Fisher Scientific), 12.5 µL of 2X Kapa HiFi HotStart ReadyMix (Roche Sequencing Solutions Inc, Pleasanton, USA), and nuclease-free water to a total volume of 25 µL. The qPCR program included the following steps: initial denaturation at 95°C for 3 min, followed by 40 cycles of 98°C for 15 seconds, annealing for 15 seconds (see **Table S7** for temperatures), and extension at 72°C for 15 seconds. Since 10 ng of oPool template resulted in the least number of cycles for amplification, qPCRs were also performed for 60G4 and 389G4 with this amount.

Subpool amplification of 12G, 12G4, 60G4, and 389G4 target oligos was accomplished by preparing 32-42 reactions with 10 ng of oPool template. Each subpool was amplified with the number of PCR cycles corresponding to the amplification plateau observed via qPCR (**Table S7**). The subpooled target oligo libraries were purified using 5 µg DNA Cleanup Columns (NEB).

### Golden Gate Assembly of Double-Stranded DNA Templates for IVT

Double-stranded DNA templates for IVT were prepared using a two-fragment Golden Gate Assembly (GGA). For each sgRNA library, 3-10 GGA reactions were performed. Each assembly reaction contained 2.0 pmol of duplexed 83 bp sgRNA scaffold oligos combined with 1.0 pmol of subpooled 98 bp oligos (either oPool or microarray-derived in origin) in an approximate 2:1 ratio. The sgRNA scaffold oligo (sequence listed in **Table S6)** was initially duplexed by annealing it to its reverse complement. However, this step was streamlined by ordering the sgRNA scaffold oligo already duplexed from Integrated DNA Technologies (IDT, Coralville, USA). The reaction mixture also included 2.5 µL of 10X T4 DNA ligase reaction buffer supplemented with 2.5 µL ATP to a final concentration of 1 mM (New England Biolabs, NEB, Ipswich, USA), 0.25 µL T4 DNA ligase (400,000 units/mL) (New England Biolabs, NEB, Ipswich, USA), 0.75 µL of BsaI-HF®v2 (20,000 units/mL) (New England Biolabs, NEB, Ipswich, USA), and nuclease-free water to a final volume of 25 µL.

The thermocycler GGA protocol consisted of 100 cycles, with each cycle containing the following steps: 37°C for 5 min and 16°C for 5 min. A heat inactivation step of 80°C for 20 min was added to denature T4 DNA ligase and BsaI-HF®v2. A 5 µg DNA Cleanup Column (NEB) was used to purify the assemblies. Verification of assemblies was performed with 2% E-Gel™ EX Agarose Gels (Thermo Fisher Scientific, Waltham, USA) with a 50 bp ladder (New England Biolabs, NEB, Ipswich, USA).

For the 18-plex and microarray-derived sgRNA libraries shown in **Fig. 1** (main text), the GGA products corresponding to the IVT DNA template (135 bp) were amplified with PCR using the SgRNA_GGA_oligo_FWD_NV and sgRNA_GGA_oligo_REV_NV primers (**Table 6)**. The resulting PCR product was size selected from agarose gels and used as the input DNA for IVT. However, this GGA product amplification step was eliminated from our workflow due to spacer coverage reductions due to excessive PCR cycles (**Figs. 1C** and **2A**).

### IVT of Pooled sgRNAs with T7 RNA Polymerase

IVT was performed using sgRNA template dsDNA oligos (135 bp) prepared with GGA. One to three IVT reactions were prepared per individual sgRNA library or subpooled library. For each standard 20 µL reaction, components were added in the following order: 0.5 µL of Murine RNase inhibitor (40,000 units/mL) (NEB), nuclease-free water, 0.01-4.5 pmol (1-400 ng) of template DNA, 2 µL of 100 mM dithiothreitol (NEB), 2 µL of 10X RNA polymerase reaction buffer (NEB), 2 µL of 25 mM ribonucleotide solution mix (NEB), 2 µL of T7 RNA polymerase (50,000 units/mL) (NEB), and nuclease-free water to a final volume of 20 μL. To prepare 100 µL reactions, all reagent volumes were scaled up 5-fold in volume, except for the input DNA amount. A negative control lacking template DNA was also prepared for all experiments. Reactions were mixed by pipetting, spun down, and incubated in a thermocycler at 37°C for two hours.

### DNase Treatment of Transcribed sgRNAs

The transcribed sgRNAs were treated with 5 µL of 10X DNase I reaction buffer (NEB), 0.5 µL of DNase I (2,000 units/mL) (NEB), and nuclease-free water to bring the total volume to 50 µL. Reactions were mixed by pipetting, spun down, and incubated at 37°C for 10 min. A 50 µg RNA Cleanup Column (NEB) was used to purify the sgRNA libraries.

### Gel Verification of sgRNA Synthesis Products

The synthesized sgRNAs were evaluated by Tris-borate-EDTA 10% polyacrylamide urea denaturing gel electrophoresis (Bio-Rad Laboratories, Hercules, USA). A single-stranded low-range RNA ladder (NEB) and an *S. pyogenes* control sgRNA sequence from the NEB EnGen® sgRNA Synthesis Kit—ordered as a synthetic sgRNA from Integrated DNA Technologies (IDT, Coralville, USA) (**Table S6**)—were included for size comparison. Samples were initially prepared by adding 2 μL of 2X TBE-urea sample loading buffer containing 7 M urea (Boston BioProducts, Milford, USA). However, due to product discontinuation, samples were later prepared with 2X RNA loading dye containing 47.5% Formamide (New England Biolabs, NEB, Ipswich, USA).

### IVT of sgRNA Libraries in Emulsions (eIVT)

First, IVT reactions were prepared using sgRNA template DNA generated through GGA. One IVT reaction was prepared per sgRNA target oligo subpool. First, components were added in the following order to prepare a 100 µL aqueous phase reaction: 2.5 µL of Murine RNase inhibitor (40,000 units/mL), nuclease-free water, 0.01-9 pmol (1-800 ng) of template DNA, 10 µL of 100 mM dithiothreitol, 10 µL of 10X RNA polymerase reaction buffer, 10 µL of 25 mM ribonucleotide solution mix, 10 µL of T7 RNA polymerase (50,000 units/mL), and 10 µL of 20 mg/mL recombinant albumin (NEB). A negative control lacking template DNA was also prepared.

An oil phase for each IVT was prepared by adding 600 µL of QX200™ Droplet Generation Oil for EvaGreen to non-stick 1.5 mL tubes (Bio-Rad Laboratories, Hercules, USA) as described by Plesa and colleagues (2018). This was later switched to Pico-Surf® (2% (w/w) in Novec™ 7500) from Sphere Bio (Cambridge, UK). A pipette was used to transfer the 100 µL aqueous phase to the bottom of the tube containing the oil phase. This process was repeated for each reaction corresponding to a unique sgRNA library. The tubes were sealed with parafilm and vortexed in Vortex Genie 2 (Scientific Industries) at 3000 rpm for 3 min. Then, the IVT reactions in emulsions (eIVT) were incubated in a thermomixer at 37°C for two hours.

To begin breaking the emulsions following incubation, eIVTs were mixed thoroughly by pipetting, and each reaction was evenly distributed between two 1.5 mL tubes, with approximately 350 µL per tube. To each tube, 175 µL of Monarch® DNA Elution Buffer (New England Biolabs, NEB, Ipswich, USA) and 612.5 µL of chloroform were added. The tubes were sealed using parafilm and then vortexed by Vortex Genie 2 at 3000 rpm for one min. Vacuum grease (phase lock gel) was applied inside the tubes, which were then centrifuged at 15,500 × g for 10 min. The 44.5 µL of upper aqueous phase pertaining to each eIVT tube was aliquoted into 0.2 mL PCR tubes and combined with 5 μL of 10X DNase I reaction buffer (NEB), 0.5 μL of DNase I (2,000 Units/mL) (NEB), and nuclease-free water to bring the total volume to 50 µL. DNase I treatment of transcribed sgRNAs was incubated at 37°C for 10 min. A 50 µg RNA Cleanup Column (NEB) was used to purify the sgRNA libraries. The sgRNA libraries were evaluated as described in the section “***Verification of sgRNA synthesis*”.**

### RNA-Seq validation of sgRNA libraries using 5’ RACE protocol

A 5’ Rapid Amplification of cDNA Ends (RACE) protocol was used for RNA-seq of sgRNA libraries, consisting of first strand synthesis, template switching, quantitative PCR (qPCR) validation, bulk PCR amplification, and product verification. We selected this 5’ RACE protocol to capture the variable spacer sequence located on the 5’ end of the sgRNAs.

For first-strand synthesis, a custom gene-specific 49 nt RT primer (RT_primer_YG, 49 nt) was annealed to the 3’ end of sgRNAs to add an overhang with a priming region and an 18 nt unique molecular identifier (UMI). Each annealing reaction included 0.5 µL Murine RNase inhibitor (40,000 units/mL) (New England Biolabs, NEB), 100 ng to 1 µg of sgRNA library, 1 µL 10 µM of RT_primer_YG (**Table S8**), 1 µL 10 µM Deoxynucleotide (dNTP) Solution Mix (NEB), and nuclease-free water to a total volume of 6 µL. Reactions were mixed by pipetting, spun down, and incubated at 70°C for 5 min.

RT and template switching were accomplished by combining one RT reaction per annealing reaction for a total volume of 10 µL. Each 4 µL RT reaction was prepared by combining 2.5 µL of 4X Template Switching RT Buffer (New England Biolabs, NEB, Ipswich, USA), 0.5 µL of 75 µM template switching oligo (TSO_YG; **Table S8**), and 1 µL of 10X Template Switching RT Enzyme Mix (NEB). The combined RT and annealing reactions were mixed by pipetting, spun down, and incubated in a thermocycler for 90 min at 42 C, followed by 5 min at 85°C.

Diluted template-switched cDNA (10 µL RT reactions diluted 2-fold) was then used for qPCR. Each qPCR included 1 µL diluted cDNA, 1.25 µL 10 µM TSO-specific primer (TSO_primer_YG; **Table S8**), 1.25 µL 10 µM gene-specific primer (Gene_specific_primer_YG; **Table S8**), 0.25 µL 100X Biotium Thiazole Green (Thermo Fisher Scientific), 12.5 µL 2X Q5 Hot-Start High-Fidelity Master Mix (NEB), and nuclease-free water to a total volume of 25 µL. The thermocycler PCR program consisted of four stages: Stage one included an initial denaturation at 98°C for 30 sec. Stage two consisted of five cycles with denaturation at 98°C for 10 sec and annealing/extension at 72°C for 5 sec per kilobase (30 sec/kb). Stage three involved five cycles with denaturation at 98°C for 10 sec and annealing at 70°C for 5 sec (30 sec/kb). Stage four comprised 40 cycles with denaturation at 98°C for 10 sec, annealing at 65°C for 15 sec, and extension at 72°C for 5 sec (30 sec/kb). To prevent over-amplification of cDNA products, the number of cycles corresponding to the amplification plateau was selected for each sgRNA library. RT samples with high cDNA yields that amplified in less than 10 cycles were further diluted twofold and re-amplified via qPCR.

Bulk PCR setup followed that of the qPCR validation, excluding the addition of Thiazole Green. Amplification followed the same PCR protocol as described earlier, except that a final extension step at 72°C for 5 min was added. Additionally, the number of cycles specified for stage four of the thermocycler PCR program was determined individually for each template based on prior qPCR analysis. PCR products were cleaned and concentrated using DNA Cleanup Columns (NEB) or GeneJET PCR clean-up columns (Thermo Fisher Scientific). The amplified cDNA products were verified on 2% or 4% E-Gel™ EX Agarose Gels (Thermo Fisher Scientific) with a 50 bp ladder (NEB).

Following verification, the cDNAs were initially extended from 186 bp to 1.049 kbp (see “***Lengthening cDNAs for Nanopore Sequencing***”) and submitted to Plasmidsaurus (South San Francisco, USA) for long-read sequencing. This was done only for the 18-plex and 10 microarray-derived sgRNA libraries shown in **Fig. 1**. The remaining sgRNA libraries were submitted without extension once Plasmidsaurus began offering short-read nanopore sequencing.

### Lengthening cDNAs for Nanopore Sequencing

Gibson assembly was used to add a pUC19 fragment to the 186 bp cDNA products of the 18-plex and microarray-derived sgRNA libraries shown in **Fig. 1**, extending their length to over 600 bp to meet the long-read nanopore sequencing requirements at the time.

To amplify a linear fragment of pUC19 for Gibson Assembly, each PCR reaction was prepared with 200 ng of pUC19 (Bayou Biolabs, Metairie, USA), 1.25 µL of 10 µM forward primer (sgRNA_cDNA_gibson_FWD_YG; **Table S8**), 1.25 µL of 10 µM reverse primer (sgRNA_cDNA_gibson_REV_YG; **Table S8**), and 12.5 µL of 2X Q5 Hot-Start High-Fidelity Master Mix (NEB). Nuclease-free water was added to bring the total reaction volume to 25 µL. The thermocycler PCR program consisted of an initial denaturation at 98°C for 30 sec, followed by 25 cycles of 98°C for 10 sec, 67°C for 30 sec, and 72°C for 45 sec, with a final extension at 72°C for 2 min. The resulting 1.576 kilobase pairs (kbp) PCR product was purified with 5 µg DNA Cleanup Columns (NEB) and digested with DpnI to remove any remaining pUC19 templates. Each DpnI digest contained 1 µg of the amplified linear pUC19 fragment, 5 µL of 10X rCutSmart Buffer (NEB), 1 µL DpnI (20,000 units/mL) (NEB), and nuclease-free water to a total volume of 50 µL. The digests were incubated in a thermocycler at 37°C for 15 min and then purified with 5 µg DNA Cleanup Columns (NEB).

Gibson Assembly was then performed to connect the linear fragment amplified from pUC19 to cDNA inserts. Each Gibson Assembly reaction included 100 ng of 1.576 kbp linear pUC19 fragment, 42.07 ng of 186 bp cDNA insert, 10 µL 2X Gibson Assembly® Master Mix, and nuclease-free water to a total volume of 20 µL. A positive control was prepared with 10 µL of NEBuilder® Positive Control and a no-template control was prepared with nuclease-free water. The assemblies were incubated in a thermocycler at 50°C for 15 min and then purified with 5 µg DNA Cleanup Columns (NEB).

PCR Amplification of Extended cDNA Product. To prepare the extended cDNA products for nanopore sequencing, a 1.049 kbp section of the 1.762 kbp Gibson Assembly product was PCR-amplified. Each PCR was prepared with 2.97 ng of the Gibson Assembly product, 1.25 µL of 10 µM forward primer (sgRNA_cDNA_1kbp_ext_FWD_YG; **Table S8**), 1.25 µL of 10 µM reverse primer (sgRNA_cDNA_1kbp_ext_REV_YG; **Table S8**), 12.5 µL of 2X Q5 Hot-Start High-Fidelity Master Mix (NEB), and nuclease-free water to a total volume of 25 µL. The thermocycler PCR program consisted of an initial denaturation at 98°C for 30 sec, followed by 25 cycles of 98°C for 10 sec, 65°C for 30 sec, and 72°C for 25 sec, with a final extension at 72°C for 2 min. The PCR products were purified with 5 µg DNA Cleanup Columns (NEB).

