## Supplementary Information for "Optimizing *in vitro* Transcribed CRISPR-Cas9 Single-Guide RNA Libraries for Improved Uniformity and Affordability"

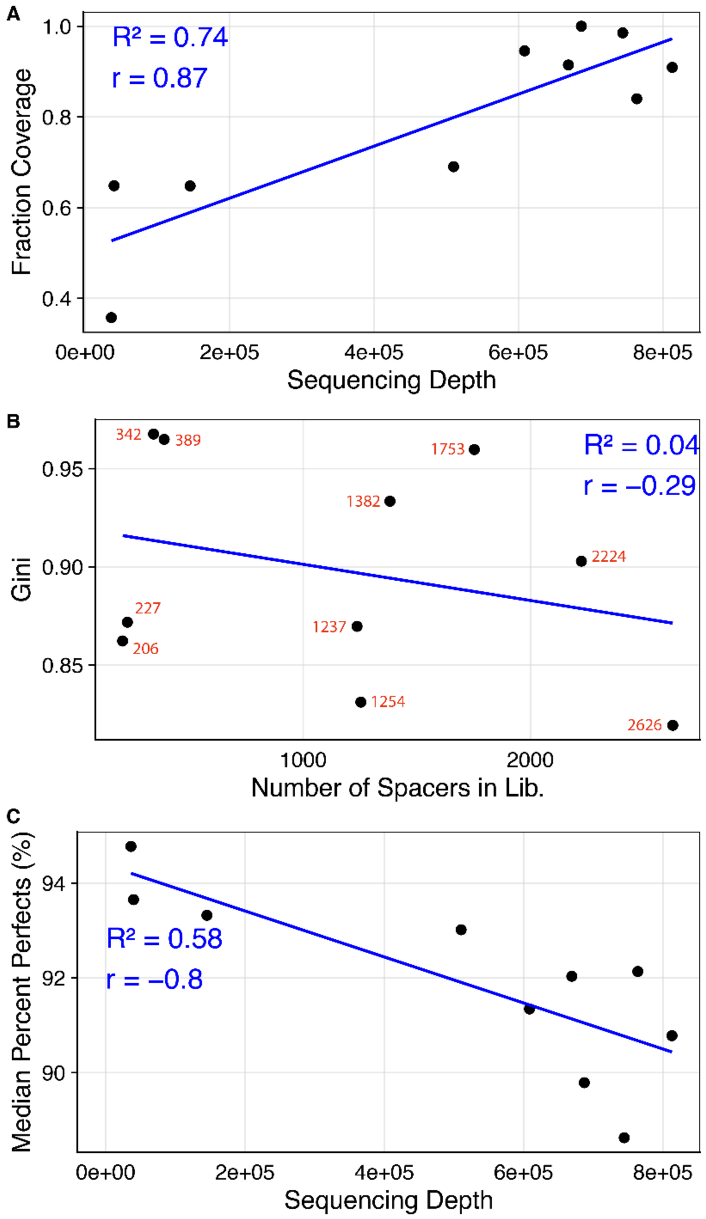


**Figure. S1. Trends in quality metrics for 10 microarray-derived sgRNA libraries. a.** Fraction of spacers observed out of expected (coverage) versus sequencing depth ($R^2 = 0.74$, $r = 0.87$). **b.** Gini Coefficients versus library scale (number of spacers in lib) ($R^2 = 0.04$, $r = -0.29$). The exact number of spacers per library are shown in red. **c.** Median percent of spacers with perfect or expected spacer sequences (median percent perfects) versus sequencing depth ($R^2 = 0.71$, $r = -0.9$).

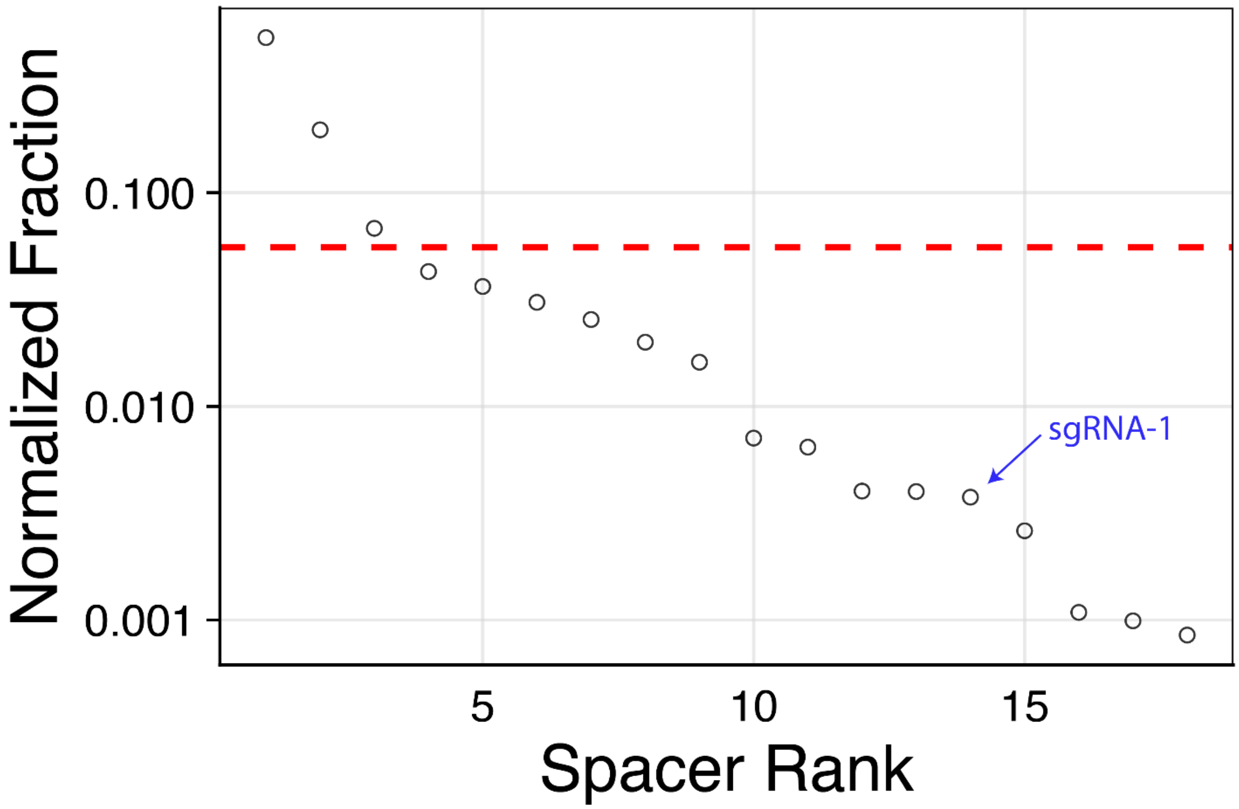


**Figure. S2. Distribution of 18 spacers transcribed as a single sgRNA library, shown by the normalized fraction of reads for perfect sequences (y-axis, log scale).** Each open circle represents an individual spacer, ranked by decreasing abundance. The dashed horizontal line indicates the expected read distribution under perfect uniformity.

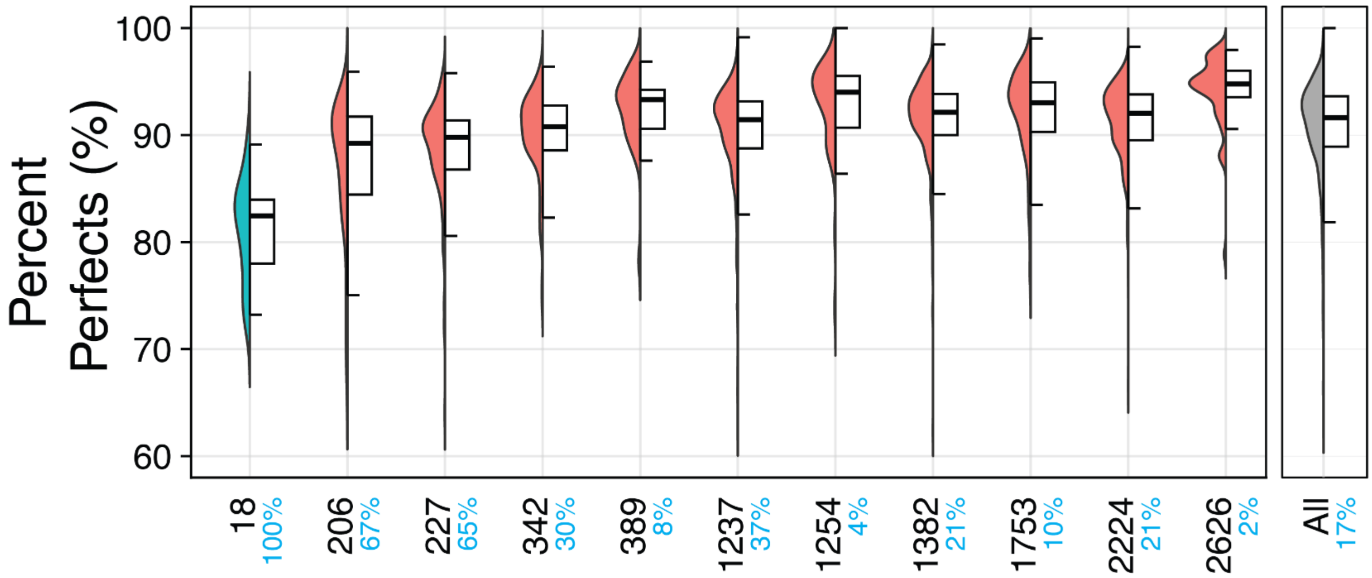


**Figure. S3. Comparison of percent perfect spacer sequences across sgRNA libraries.** Perfect spacer sequences correspond to the expected spacers programmed into each library. Each library is represented by a bifurcated plot: the left half shows a half-violin plot (distribution of percent perfects), and the right half displays a boxplot (median percent perfect per library). Only spacers with at least 100 reads in the RNA-seq data were included to ensure reliable analysis. The percentage of spacers meeting this threshold is indicated on the x-axis (blue text). The column-synthesized 18-plex sgRNA library is shown in teal, microarray-derived libraries in orange, and the overall median percent perfects across all 11 libraries is represented by the gray bar.

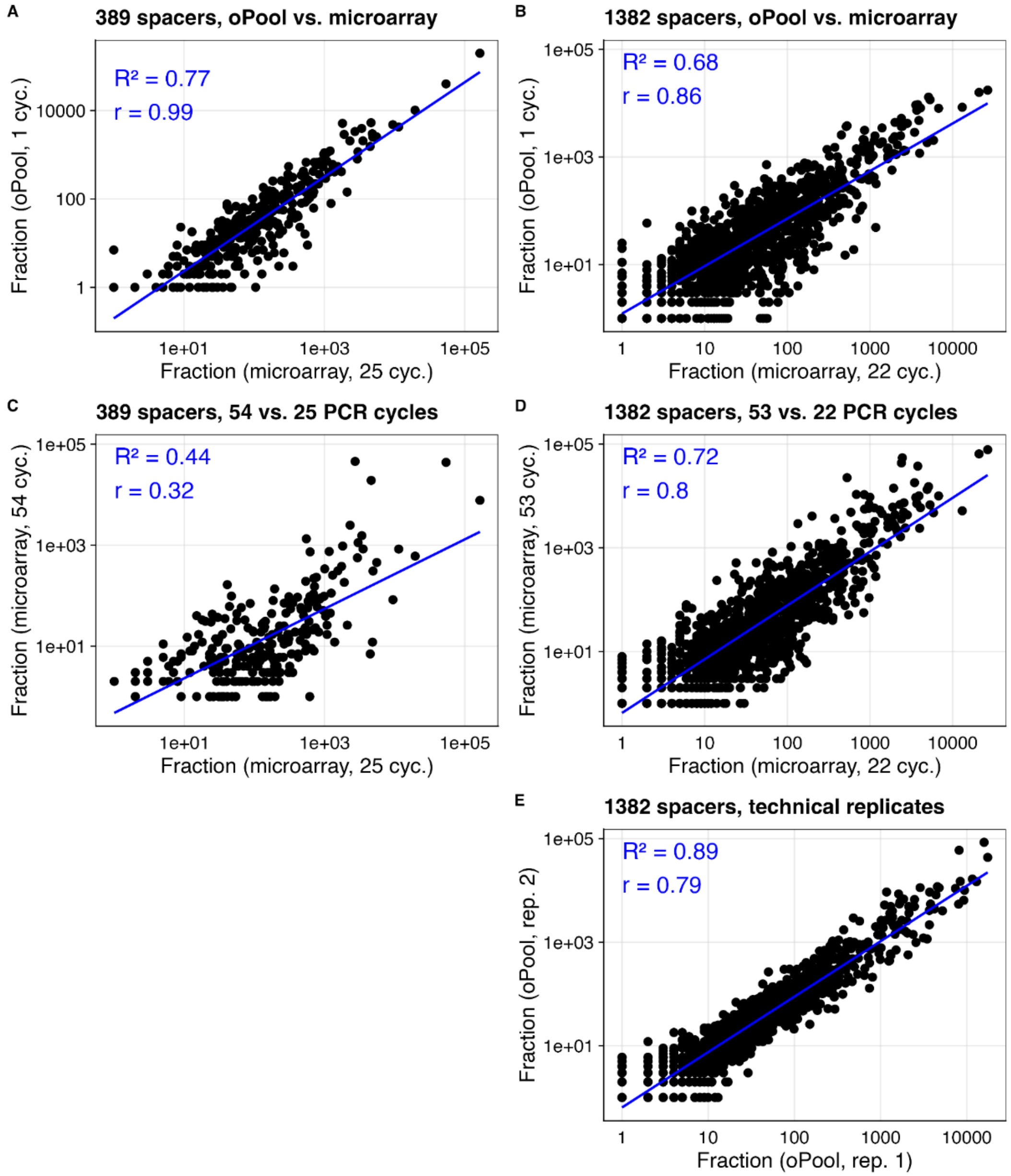


**Figure. S4. Pearson correlations of spacer reads for oPool- and microarray-derived sgRNA libraries prepared with varying PCR cycle numbers. a.** oPool-derived 389-plex sgRNA library spacer reads (1 cycle) versus a microarray-derived version prepared with 25 PCR cycles ($R^2 = 0.77$, $r = 0.99$). **b.** oPool-derived 1,382-plex sgRNA library spacer reads (1 cycle) versus a microarray-derived version prepared with 22 PCR cycles ($R^2 = 0.68$, $r = 0.86$). **c.** Microarray-derived 389-plex sgRNA library spacer reads (52 cycles) versus a reduced-PCR version prepared with 25 PCR cycles ($R^2 = 0.44$, $r = 0.32$). **d.** Microarray-derived 1,382-plex sgRNA library spacer reads (53 cycles) versus a reduced-PCR version prepared with 22 PCR cycles ($R^2 = 0.72$, $r = 0.80$). **e.** Comparison of oPool-derived 1,382-plex sgRNA library spacer reads between two replicates ($R^2 = 0.89$, $r = 0.79$).

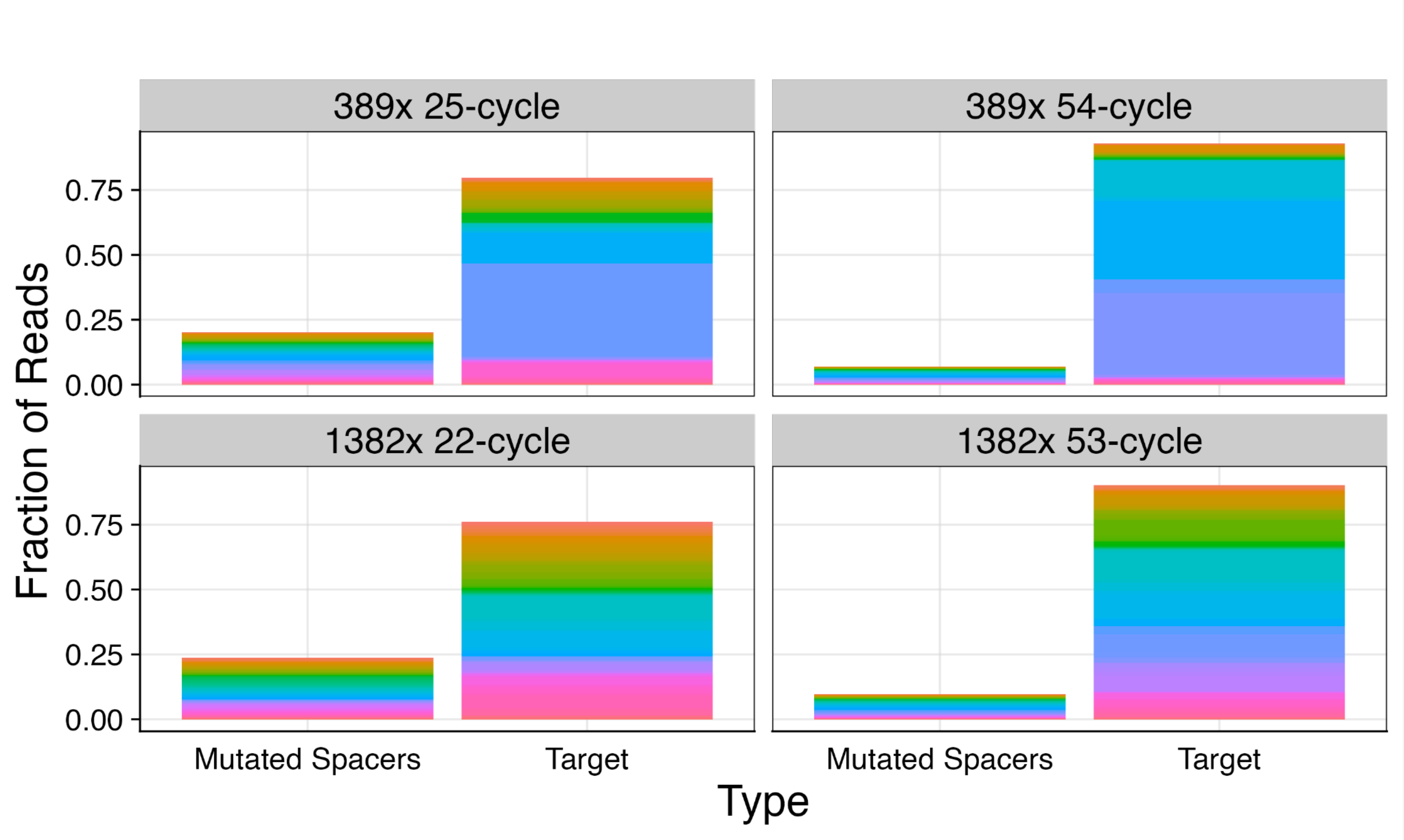


**Figure. S5. Effect of PCR cycle number on the fraction of mutant and target spacers in 389- and 1,382-plex microarray-derived sgRNA libraries.** Fraction of total reads, scaled to one, representing mutant spacers that deviate from the expected sequence compared to target spacers with the correct expected sequences. The top panel shows these metrics for the 389-plex sgRNA library prepared with reduced PCR using 25 cycles and excessive PCR using 54 cycles. The bottom panel shows the same for the 1,382-plex sgRNA library, comparing reduced PCR with 22 cycles to excessive PCR with 53 cycles. Each color within the rainbow pattern represents a unique spacer sequence, whether mutant or target.

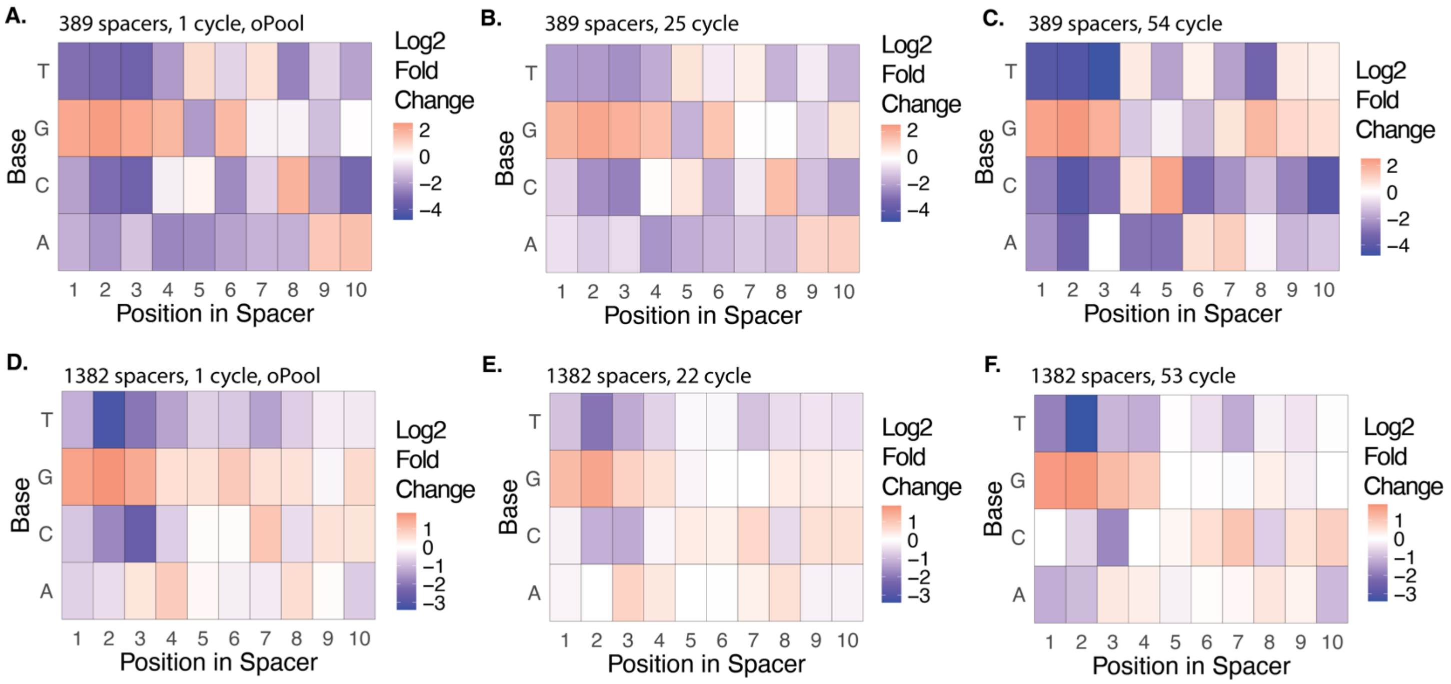


**Figure. S6. Influence of base composition on spacer abundance within 389- and 1,382-plex sgRNA libraries prepared with differing oligo sources and PCR cycle numbers.** $Log_2$ fold change (FC) of observed vs. expected spacer abundance across all nucleotide identities within the first 10 nucleotides of 20-nt spacers. "Observed" refers to the fraction of spacers identified from RNA-seq data that meet the sequence criteria, while "expected" represents the fraction assuming perfect uniformity across the population. Positive $Log_2$ FC values (orange) within the heat maps indicate increased spacer abundance compared to expected, while negative values (blue) indicate decreased abundance. Zero change is shown in white. The $Log_2$ fold change scale ranges from -4 to 2 for the 389-plex library and -3 to 1 for the 1,382-plex library. **a.** oPool-derived 389-plex spacer library prepared with one PCR cycle. **b.** Microarray-derived 389-plex spacer library prepared with reduced PCR cycles (25 total). **c.** Microarray-derived 389-plex spacer library prepared with excessive PCR cycles (54 total). **d.** oPool-derived 1,382-plex spacer library prepared with one PCR cycle. **e.** Microarray-derived 1,382-plex spacer library prepared with reduced PCR cycles (22 total). **f.** Microarray-derived 1,382-plex spacer library prepared with excessive PCR cycles (53 total).

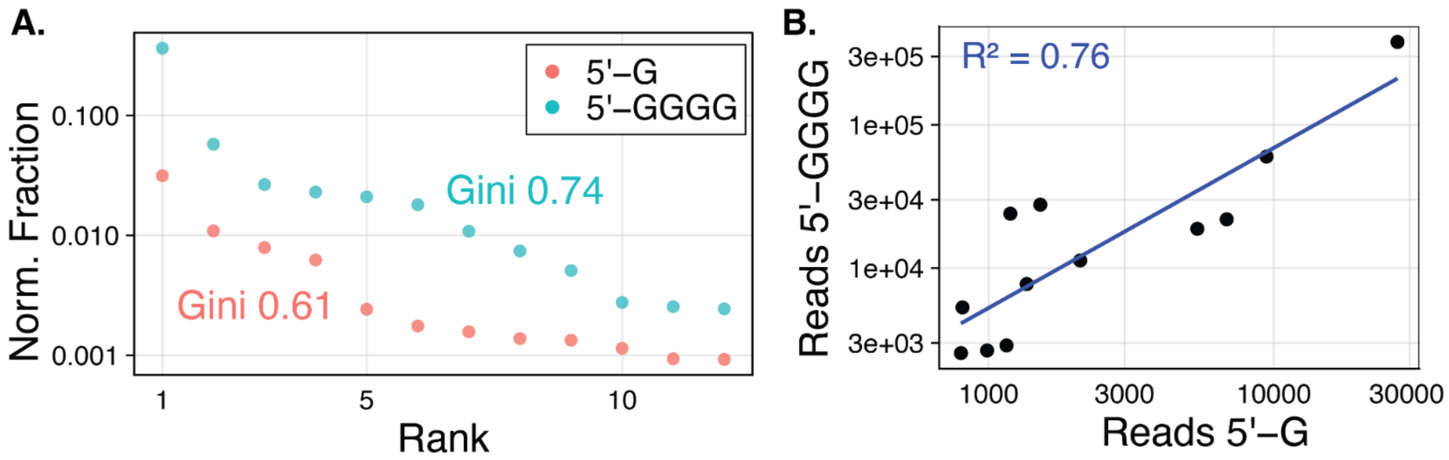


**Figure. S7. Distribution of 12 sgRNA spacers starting with a 5' G and 5' GGGG (12G and 12G4 libraries). a.** Comparison of spacer distribution uniformity between two 12-plex sgRNA libraries containing identical spacers, except that the spacers in one library start with a single 5′ guanine (5′ G, orange), while the spacers in the other library are padded with a 5′ guanine tetramer (5’ GGGG, teal). Normalized spacer abundance is shown, with each dot representing a unique spacer, ranked in descending order. Gini Coefficients (Gini), shown alongside each library's spacer distribution, measure the inequality in spacer representation between the two library types. **b.** Pearson correlation ($R^2$ = 0.76, r = 0.962) of spacer abundance between the two sgRNA libraries presented in **a.**

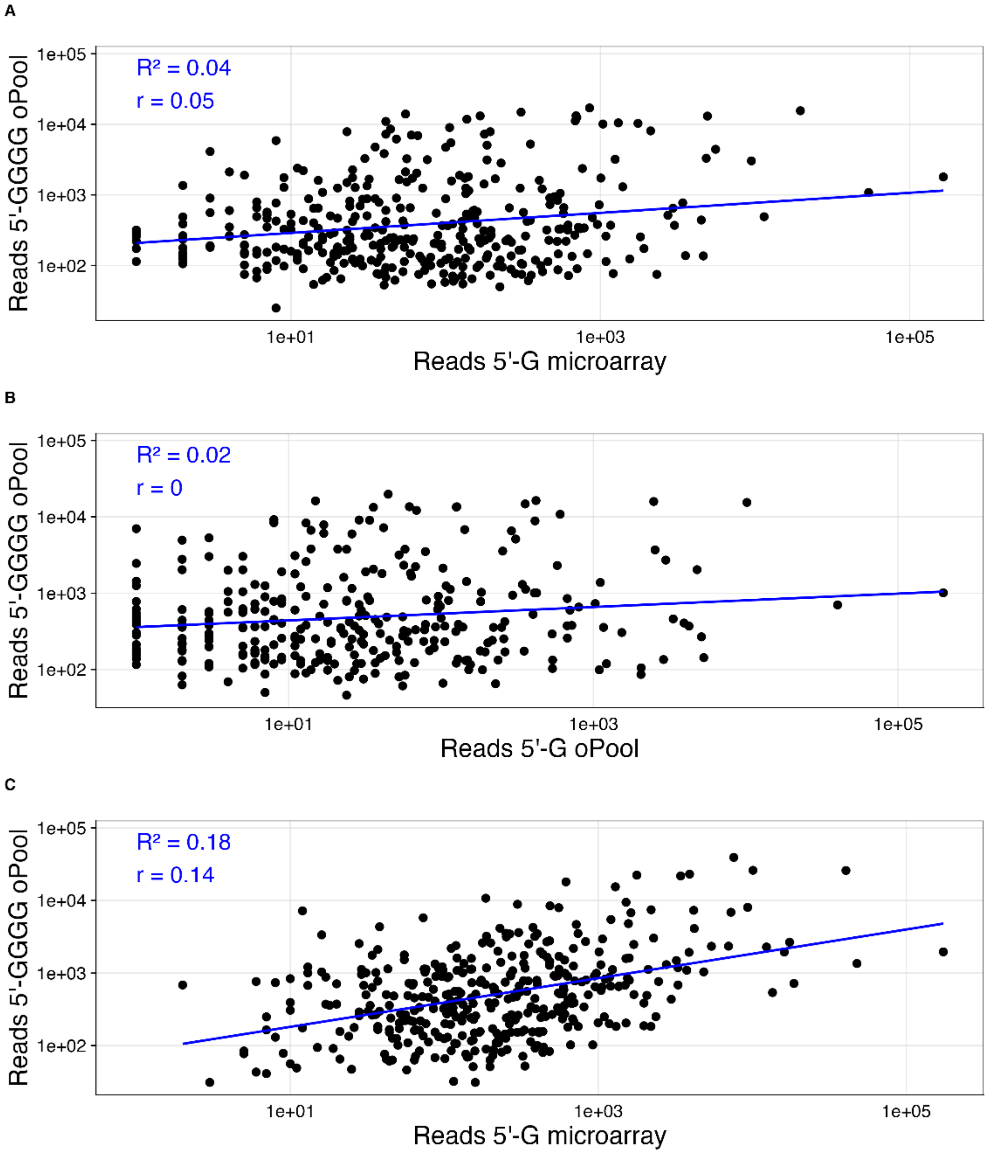


**Figure. S8. Pearson correlations of spacer reads for sgRNA libraries containing 389 spacers starting with 5' G and 5' GGGG (389G and 389G4 libraries). a.** oPool-derived sgRNA library (7 PCR cycles) with four guanines compared to a microarray-derived sgRNA library (25 PCR cycles) with a single guanine ($R^2 = 0.04$, $r = 0.05$). **b.** oPool-derived sgRNA library with four guanines (7 PCR cycles) compared to an oPool-derived sgRNA library with a single guanine (1 PCR cycle) ($R^2 = 0.02$, $r = 0$). **c.** Replicate of the condition shown in A using different sgRNA libraries ($R^2 = 0.18$, $r = 0.14$).

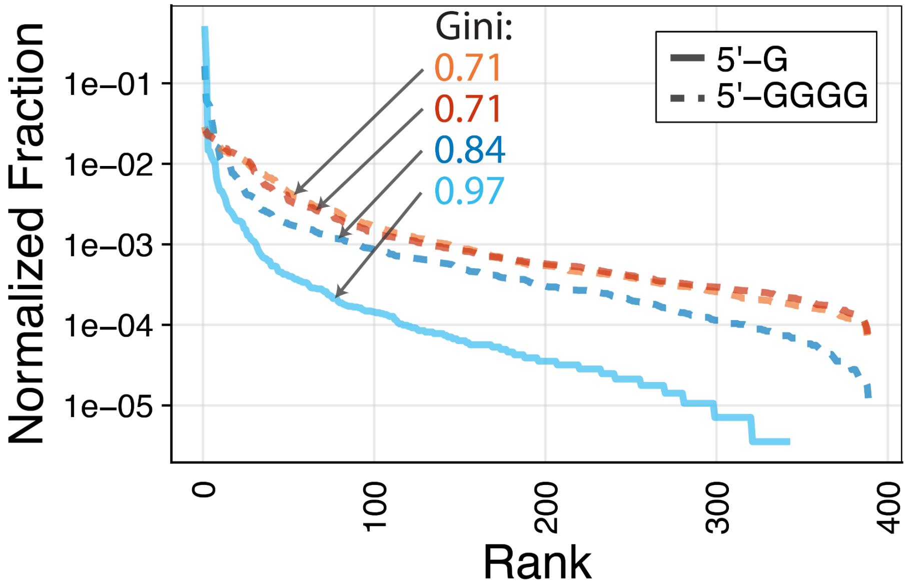


**Figure. S9. Gini Coefficients of sgRNA libraries containing 389 spacers starting with 5' G and 5' GGGG (389G and 389G4 libraries) transcribed under modified *in vitro* transcription (IVT) conditions.** Normalized fraction of spacer reads relative to total reads for sgRNA libraries containing 389 spacers, either starting with 5’ G (solid line, n = 1) or 5’ GGGG (dashed lines, n = 3). Spacers are ranked by decreasing abundance. Each library, transcribed *in vitro* from 100 ng of template DNA in a 100 µL reaction volume, is represented by a distinct color. Gini Coefficients (Gini) are listed next to each library’s rank-ordered curve to quantify inequality in spacer representation across library types and scales.

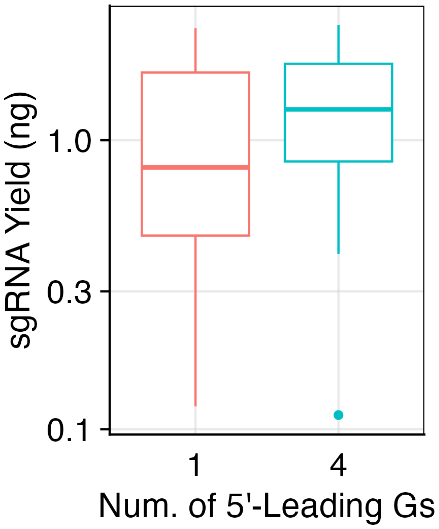


**Figure. S10. Comparison of sgRNA yields between libraries with 5’ G and 5’ GGGG spacers.** Comparison of sgRNA yields (µg) for sgRNA libraries containing spacers that start with 5’ G (median: 0.805 µg, n = 13) versus 5’ GGGG (median: 1.29 µg, n = 8). Differences in yield were not statistically significant (Wilcoxon rank-sum test, p = 0.5002).

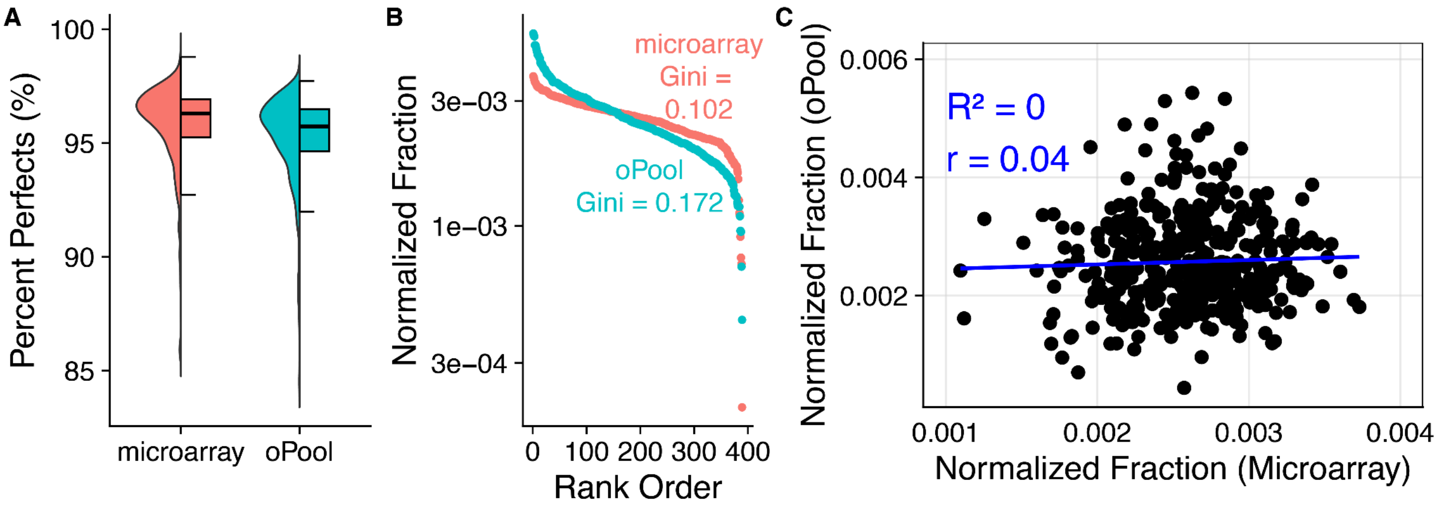


**Figure. S11. Comparison of quality metrics for 135 bp DNA libraries containing 389 spacers, prepared from microarray- and oPool-derived oligos. a.** The percentage of perfect spacers, defined as spacers without mutations, was assessed in microarray-derived (coral, median: 96.30%) and oPool-derived (teal, median: 95.73%) DNA libraries. Each library is represented as a bifurcated plot: the left half shows a half violin plot (distribution of percent perfect spacers), while the right half displays a boxplot (median percent perfect for each library). Only spacers with at least 100 reads in the RNA-seq data were included to ensure reliable analysis. **b.** Normalized abundance (reads per spacer, relative to total reads) for the same libraries as shown in **a.** Spacers are ranked in descending order of abundance. Gini coefficients (Gini), listed next to each library's rank-ordered curve, indicate inequality in spacer distribution, showing a significant difference in distribution between libraries (Wilcoxon rank-sum test, $p = 3.97 \times 10^{-8}$). **c.** Pearson correlation analysis of normalized fraction of spacer reads between the microarray- versus oPool-derived DNA libraries ($R^2 = 0$, $r = 0.04$).

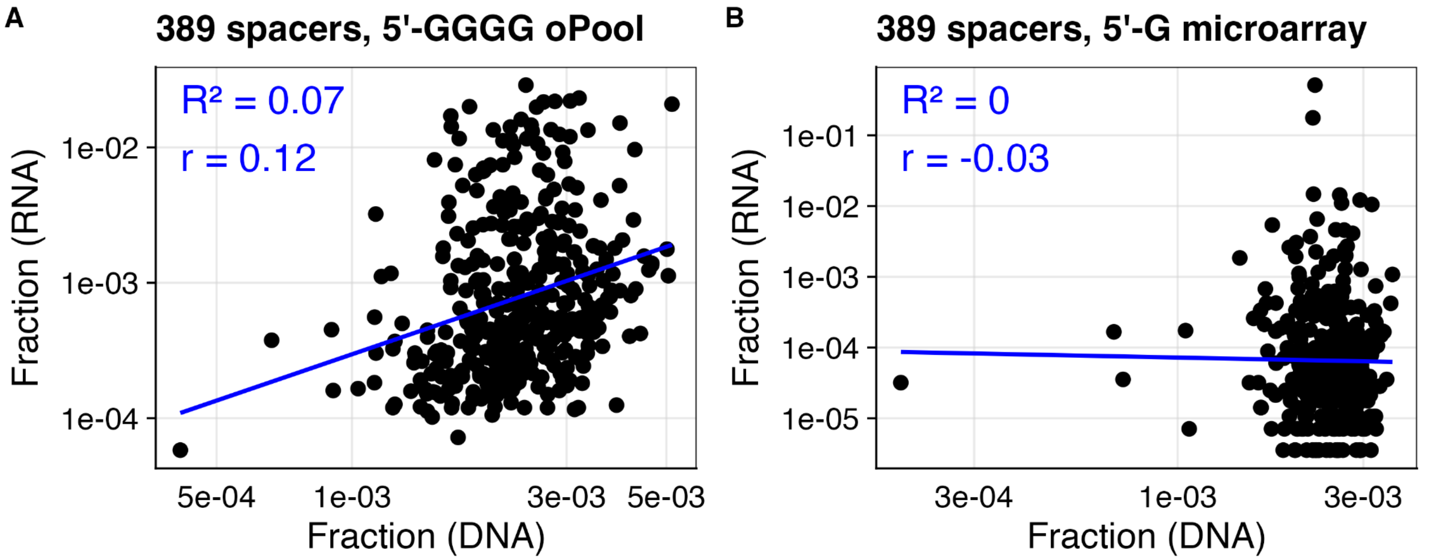


**Figure. S12. Pearson correlations of spacer reads between 135 bp DNA libraries containing 389 spacers and their corresponding *in vitro* transcribed sgRNA libraries. a.** Comparison of an oPool-derived DNA library and its corresponding sgRNA library, transcribed using 100 ng input DNA in a 100 µL IVT reaction volume. Both libraries contain spacers that start with 5' GGGG ($R^2 = 0.07$, $r = 0.12$). **b.** Comparison of a microarray-derived DNA library and its corresponding sgRNA library, transcribed using 100 ng input DNA in a 20 µL IVT reaction volume. Both libraries contain spacers that start with 5' G ($R^2 = 0$, $r = -0.03$).

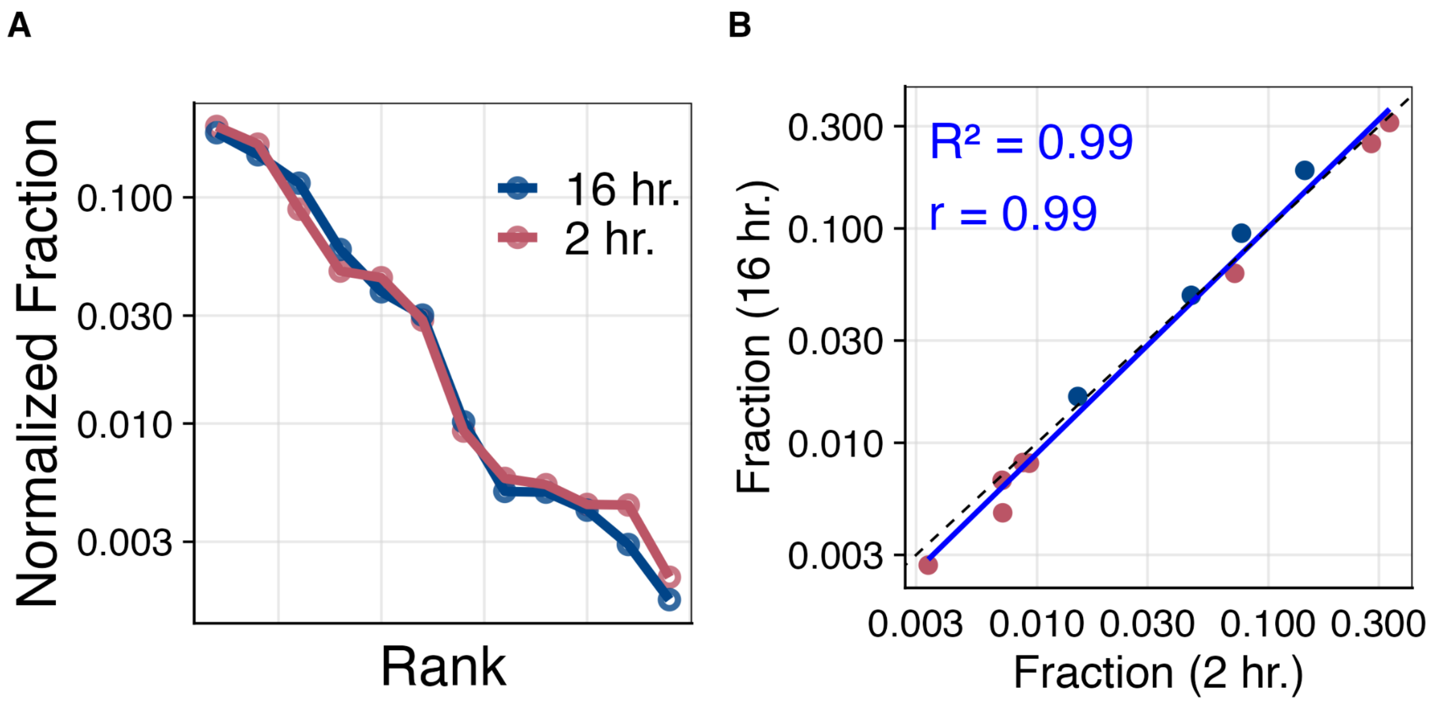


**Figure. S13. Comparison of incubation times for transcribing a 12-plex sgRNA library in emulsions. a.** Spacers ordered by decreasing normalized fraction of reads (rank) for libraries transcribed at 37°C for 2 hours (red) and 16 hours (blue). **b.** Pearson correlation analysis of the fraction of spacer reads for sgRNA libraries transcribed for 2 hours (red) and 16 hours (blue) ($R^2 = 0.99$, $r = 0.99$).

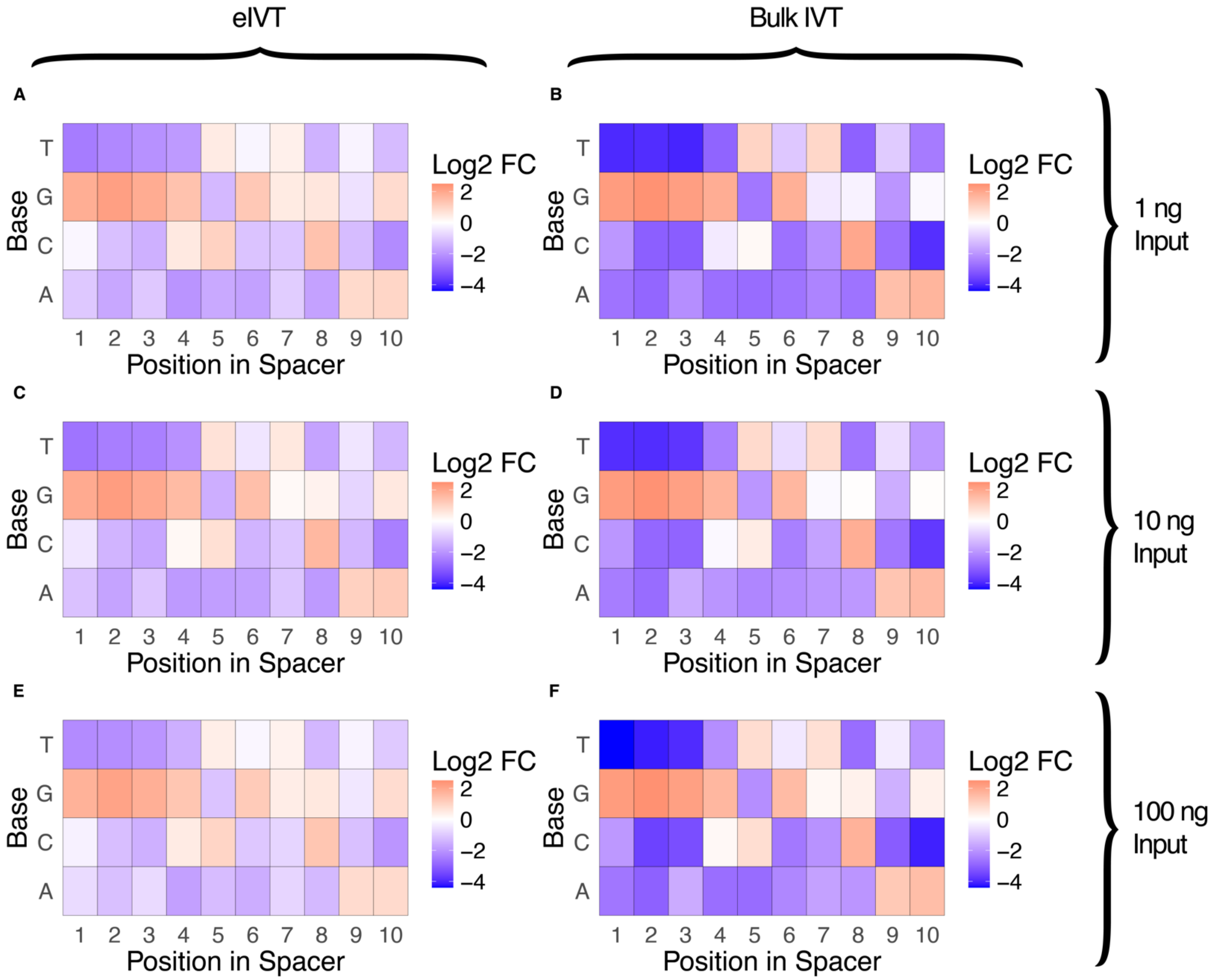


**Figure. S14. Effect of base composition on spacer abundance in 389-plex sgRNA libraries with 5' G (389G library), transcribed via bulk IVT or emulsion IVT (eIVT).** $Log_2$ fold change (FC) of observed vs. expected spacer abundance across all nucleotide identities within the first 10 nucleotides of 20-nt spacers. "Observed" refers to the fraction of spacers identified from RNA-seq data that meet the sequence criteria, while "expected" represents the fraction assuming perfect uniformity across the population. Positive $Log_2$ FC values (orange) indicate increased spacer abundance compared to expected, while negative values (blue) indicate decreased abundance. Zero change is shown in white. The FC scale ranges from -4 to 2 across all heatmaps. **a.** eIVT with 1 ng input DNA **b.** Bulk IVT with 1 ng input DNA **c.** eIVT with 10 ng input DNA **d.** Bulk IVT with 10 ng input DNA **e.** eIVT with 100 ng input DNA **f.** Bulk IVT with 100 ng input DNA

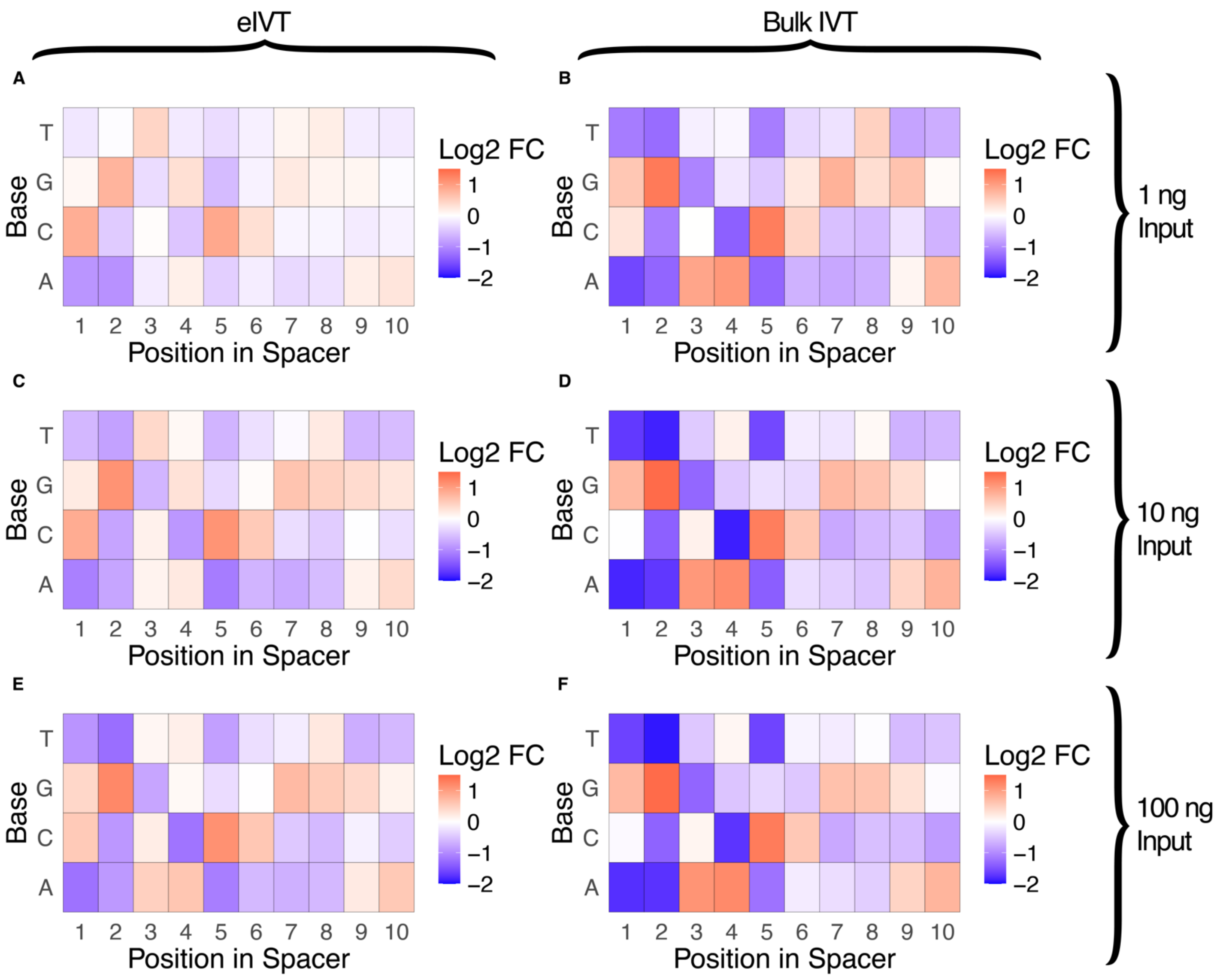


**Figure. S15. Effect of base composition on spacer abundance in 2,626-plex sgRNA libraries with 5' G, transcribed via bulk IVT or emulsion IVT (eIVT).** $Log_2$ fold change (FC) of observed vs. expected spacer abundance across all nucleotide identities within the first 10 nucleotides of 20-nt spacers. "Observed" refers to the fraction of spacers identified from RNA-seq data that meet the sequence criteria, while "expected" represents the fraction assuming perfect uniformity across the population. Positive $Log_2$ FC values (orange) indicate increased spacer abundance compared to expected, while negative values (blue) indicate decreased abundance. Zero change is shown in white. The FC scale ranges from -2 to 2 across all heatmaps. **a.** eIVT with 1 ng input DNA **b.** Bulk IVT with 1 ng input DNA **c.** eIVT with 10 ng input DNA **d.** Bulk IVT with 10 ng input DNA **e.** eIVT with 100 ng input DNA **f.** Bulk IVT with 100 ng input DNA

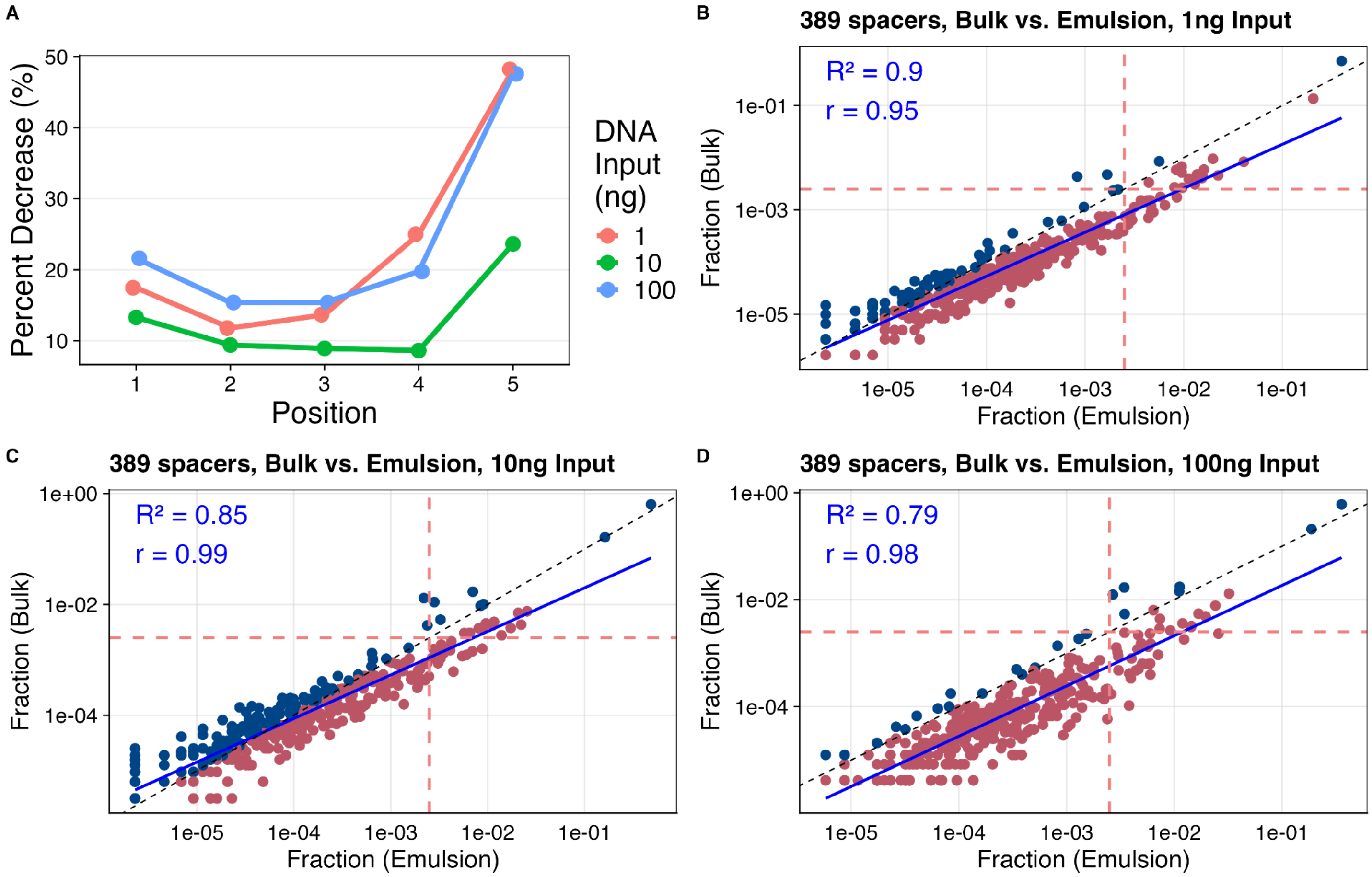


**Figure. S16. Spacer representation changes in 389-plex sgRNA libraries with 5' G, transcribed via eIVT with varying DNA input amounts. a**. The percentage decrease in $Log_2$ fold change (FC) of observed vs. expected abundance reflects the reduction in spacer position-dependent guanine representation in eIVT compared to bulk IVT, calculated using the formula (FC_bulk - FC_eIVT) / FC_bulk. This analysis was performed for sgRNA libraries prepared with 1 ng (coral), 10 ng (green), and 100 ng (aqua) input DNA **b-d.** Pearson correlation analysis of spacer representation across input DNA amounts: **b.** 1 ng ($R^2 = 0.9$, $r = 0.95$), **c.** 10 ng ($R^2 = 0.85$, $r = 0.99$), and **d.** 100 ng ($R^2 = 0.79$, $r = 0.98$).The solid blue line represents the linear fit of spacer reads, while the dashed black line indicates the unity fit (1:1 relationship) between bulk IVT and eIVT. Blue dots represent overrepresented spacers that decrease in eIVT, while red dots represent underrepresented spacers that increase in eIVT. The red dashed lines represent the median read fraction per spacer from the input DNA library, which had a highly uniform spacer distribution and served as the template for sgRNA transcription.

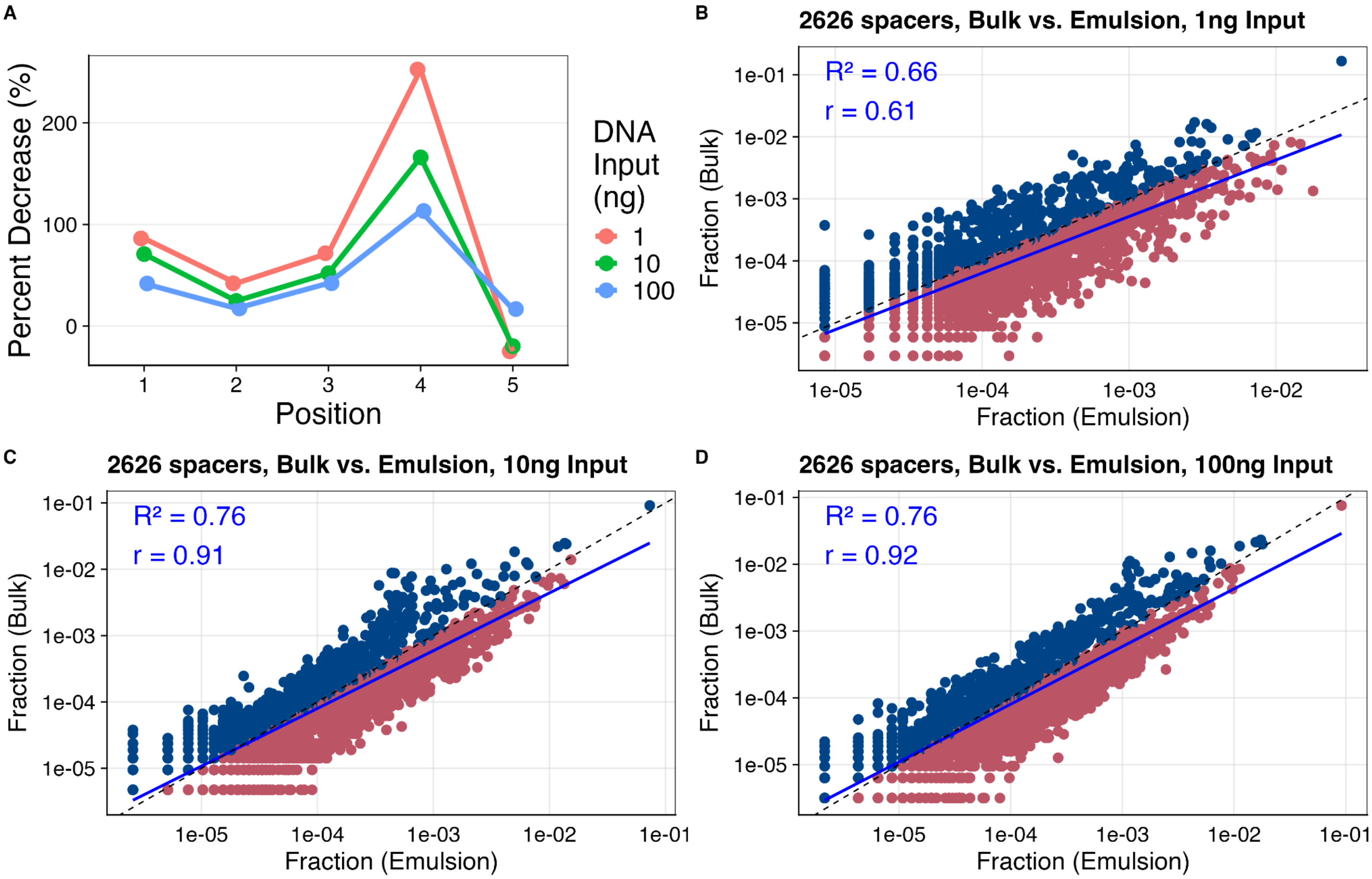


**Figure. S17. Spacer representation changes in 2,626-plex sgRNA libraries with 5' G, transcribed via eIVT with varying DNA input amounts. a.** The percentage decrease in $Log_2$ fold change (FC) of observed vs. expected abundance reflects the reduction in spacer position-dependent guanine representation in eIVT compared to bulk IVT, calculated using the formula (FC_bulk - FC_eIVT) / FC_bulk. This analysis was performed for sgRNA libraries prepared with 1 ng (coral), 10 ng (green), and 100 ng (aqua) input DNA. **b-d.** Pearson correlation analysis of spacer representation across input DNA amounts: **b.** 1 ng ($R^2 = 0.66$, $r = 0.61$), **c.** 10 ng ($R^2 = 0.76$, $r = 0.91$), and **d.** 100 ng ($R^2 = 0.76$, $r = 0.92$).The solid blue line represents the linear fit of spacer reads, while the dashed black line indicates the unity fit (1:1 relationship) between bulk IVT and eIVT. Blue dots represent overrepresented spacers that decrease in eIVT, while red dots represent underrepresented spacers that increase in eIVT.

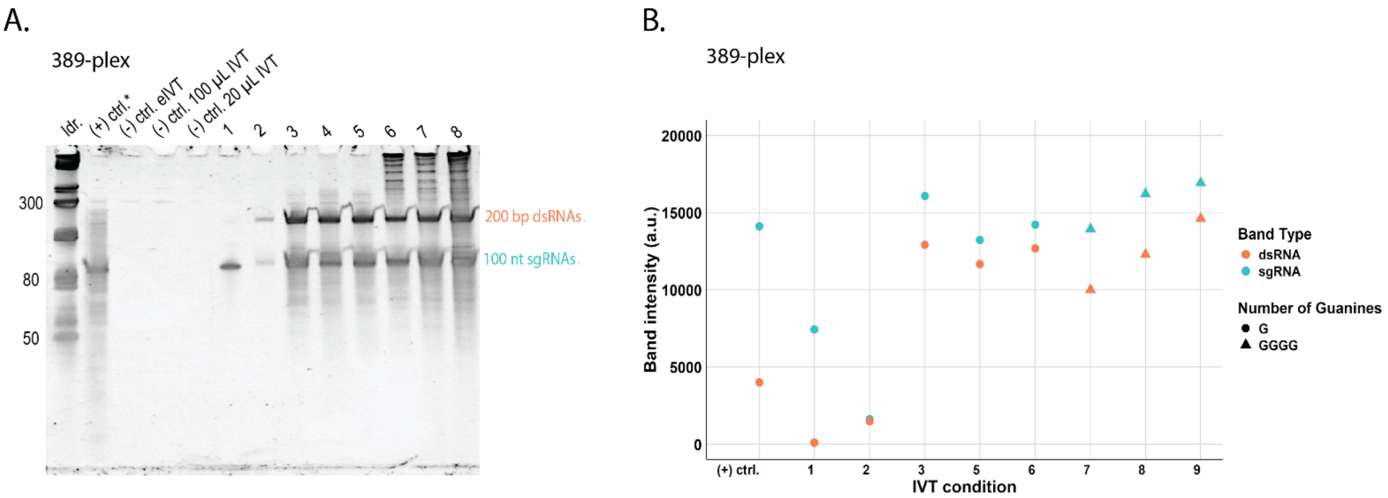


| IVT condition | IVT/eIVT | Input DNA (ng) | Aq. rxn vol (µL) | 5′ guanines (G) |
|---|---|---|---|---|
| 1 | eIVT | 10 | 100 | G |
| 2 | IVT | 10 | 20 | G |
| 3 | eIVT | 100 | 100 | G |
| 4 | IVT | 100 | 100 | G |
| 5 | IVT | 100 | 20 | G |
| 6 | eIVT | 100 | 100 | GGGG |
| 7 | IVT | 100 | 100 | GGGG |
| 8 | IVT | 400 | 20 | GGGG |

*(+) ctrl: Synthetic sgRNA control sequence

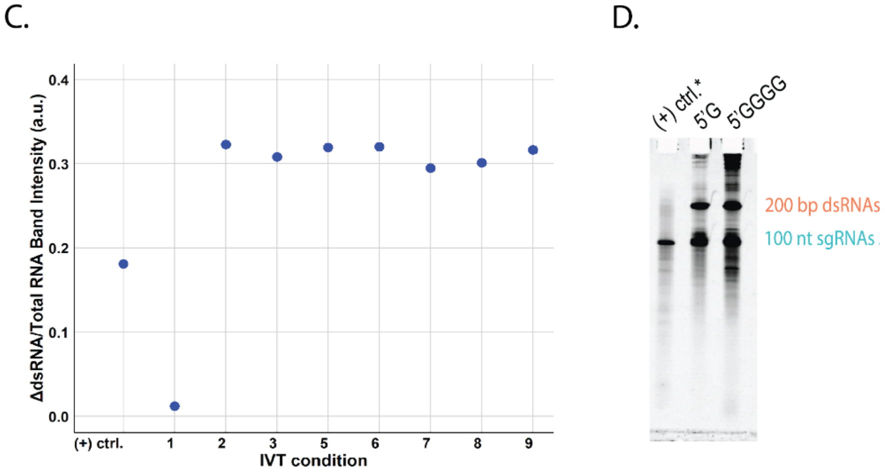


**Figure. S18. Assessment of single-stranded sgRNA (100 nt) products and high molecular weight (200 bp+) RNA byproducts between 5' G and 5' GGGG sgRNA libraries transcribed in emulsions or bulk. a.** Tris-borate-EDTA-urea denaturing gel electrophoresis (10% polyacrylamide) with SYBR Gold staining was used to analyze 389-plex sgRNA libraries transcribed under different conditions. These conditions included bulk IVT (IVT), emulsion IVT (eIVT), varying DNA input DNA amounts, and spacers starting with either 5' G or a 5' GGGG. The aqueous reaction volume (aq. rxn vol) was also varied, representing the total aqueous IVT volume emulsified in eIVT or the total reaction volume in standard IVT. **b.** Quantification of gel band intensities corresponding to sgRNA (100 nt) and dsRNA (200 bp) from (A), plotted in arbitrary units (A.U.). **c.** Ratios of dsRNA band intensities were calculated relative to the total RNA, defined as the sum of sgRNA and dsRNA intensities, for the synthetic sgRNA control and various IVT conditions. **d.** Tris-borate-EDTA-urea denaturing gel electrophoresis (10% polyacrylamide) with SYBR Gold staining was used to assess transcribed RNA products from libraries containing 12 identical spacers. These spacers started with either 5’ G or a 5' GGGG and were compared to a synthetic control sgRNA sequence (100 nt).

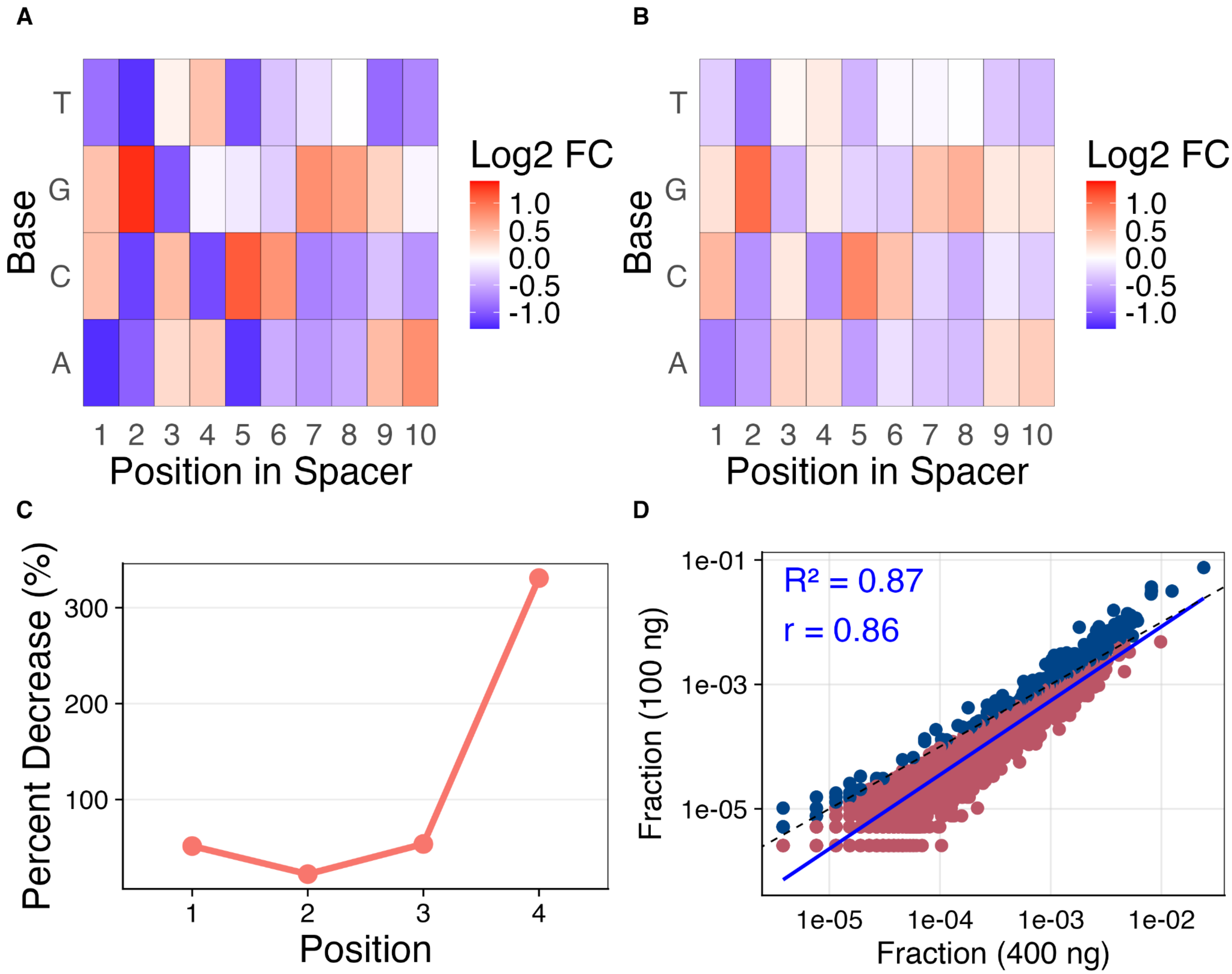


**Figure. S19. Spacer representation changes in 2,626-plex sgRNA libraries with 5' G transcribed with 100 ng or 400 ng input DNA in a 20 µL IVT reaction volume. a-b.** $Log_2$ fold change (FC) of observed vs. expected spacer abundance across all nucleotide identities within the first 10 nucleotides of 20-nt spacers for sgRNA libraries transcribed with **a.** 100 ng and **b.** 400 ng input DNA. "Observed" refers to the fraction of spacers identified from RNA-seq data, while "expected" represents the fraction assuming perfect uniformity across the population. Positive $Log_2$ FC values (orange) indicate increased spacer abundance compared to expected, while negative values (blue) indicate decreased abundance. **c.** Percent decrease in $Log_2$ FC of observed vs. expected abundance, calculated separately for IVT with 100 ng and 400 ng input DNA using the formula (FC_100ng - FC_400ng) / FC_100ng. The difference (ΔFC) between the two conditions was plotted to show changes in position-dependent guanine representation across spacers. **d.** Pearson correlation analysis of the fraction of spacer reads for sgRNA libraries transcribed with 100 ng and 400 ng input DNA ($R^2 = 0.87$, $r = 0.86$). The solid blue line represents the linear fit of spacer reads, while the dashed black line indicates the unity fit (1:1 relationship) between bulk IVT and eIVT. Blue dots represent overrepresented spacers that decrease in eIVT, while red dots represent underrepresented spacers that increase in eIVT.

**Table S1 – Pearson correlations among various sgRNA library metrics for 10 microarray-derived sgRNA libraries.** The Pearson correlation coefficient (*r*) is shown for each pairwise comparison. Selected variable name abbreviations include median percent perfects (medpperf), total PCR cycles (cyctotal), and sequencing depth (depth).

| Variable 1 | Variable 2 | Pearson coefficient (*r*) |
|---|---|---|
| Gini | Scale | -0.28674 |
| Gini | depth | 0.385077 |
| Gini | medpperf | -0.06484 |
| Gini | cyctotal | 0.022516 |
| Gini | coverage | 0.211416 |
| Scale | depth | -0.35841 |
| Scale | medpperf | 0.694729 |
| Scale | cyctotal | -0.55549 |
| Scale | coverage | -0.57874 |
| depth | medpperf | -0.80169 |
| depth | cyctotal | 0.063687 |
| depth | coverage | 0.865335 |
| medpperf | cyctotal | -0.39103 |
| medpperf | coverage | -0.89704 |
| cyctotal | coverage | 0.271306 |

**Table S2. Kolmogorov–Smirnov (KS) statistical analysis of spacer distribution changes between eIVT and bulk IVT sgRNA libraries.** KS test analysis was applied to normalized sgRNA sequencing data for 389- and 2,626-plex sgRNA libraries, each containing spacers starting with 5’ G, transcribed in bulk or emulsions (see **Fig. 4c** in the main text). Each sgRNA library was transcribed once ($n = 1$) with varying reaction scales and DNA input amounts. The KS test was used to assess the maximum discrepancy between the cumulative distribution functions of sgRNAs transcribed in emulsions compared to their corresponding bulk IVT control libraries. A significant KS statistic ($p < 0.05$) indicates a substantial shift in sgRNA distribution between the two conditions, suggesting changes in library composition or uniformity.

| DNA Amount (ng) | Library Scale | D value | p value |
|---|---|---|---|
| 1 | 389 | 0.20352 | 3.53E-07 |
| 10 | 389 | 0.097695 | 5.40E-02 |
| 100 | 389 | 0.36263 | 2.20E-16 |
| 1 | 2626 | 0.22022 | 2.20E-16 |
| 10 | 2626 | 0.098369 | 7.35E-10 |
| 100 | 2626 | 0.081096 | 3.19E-07 |

**Table S3. Kolmogorov–Smirnov (KS) analysis of spacer distribution changes in sgRNA libraries transcribed with bulk IVT using 1 ng, 10 ng, or 100 ng input DNA.** KS test analysis of normalized sgRNA sequencing data for libraries containing 389 and 2,626 spacers, all starting with 5' G. The KS test was used to evaluate the maximum discrepancy between the cumulative distribution functions of sgRNAs transcribed with 1-400 ng of input DNA, with IVT reaction volumes of 20 µL or 100 µL. A significant KS statistic ($p < 0.05$) indicates a substantial shift in sgRNA distribution between the two conditions, suggesting changes in library composition or uniformity.

| **Comparing DNA Input Amount (in ng)** | | | | | |
|---|---|---|---|---|---|
| Total IVT Volume (µL) | Library Scale | DNA Amount 1 | DNA Amount 2 | D value | p value |
| 20 | 2626 | 1 | 10 | 0.064611 | 0.000105 |
| 20 | 2626 | 10 | 100 | 0.08759 | 3.96E-08 |
| 20 | 2626 | 100 | 400 | 0.24361 | 2.20E-16 |
| 20 | 2626 | 1 | 400 | 0.21657 | 2.20E-16 |
| 100 | 2626 | 1 | 10 | 0.085496 | 1.59E-07 |
| 100 | 2626 | 10 | 100 | 0.06849 | 4.18E-05 |
| 100 | 2626 | 1 | 100 | 0.05971 | 0.000464 |
| 100 | 389 | 1 | 10 | 0.1675 | 4.69E-05 |
| 100 | 389 | 10 | 100 | 0.2289 | 3.30E-08 |
| 100 | 389 | 1 | 100 | 0.060506 | 0.5585 |
| 20 | 389 | 1 | 10 | 0.088837 | 0.112 |
| 20 | 389 | 10 | 100 | 0.18941 | 2.49E-06 |
| 20 | 389 | 100 | 400 | 0.23818 | 6.19E-10 |

**Table S4.** The 18-target single-stranded DNA oligonucleotide sequences for an 18-plex sgRNA target library. Each sequence includes a forward primer site, T7 promoter sequence, 20 nt spacer, and reverse primer site.

| Target oligo sequence name | Sequence |
|---|---|
| Pool_002_sgRNA_1 | GGGTCACGCGTAGGATTCTAATACGACTCACTATAGGCGTTCAATTTAGATTAGGGTTTTAGAGCTAGAGGAGACCGTTCTCGTGTGGCTGCGGAAC |
| Pool_002_sgRNA_2 | GGGTCACGCGTAGGATTCTAATACGACTCACTATAGGCTATTGTTTCTAAAATGCGTTTTAGAGCTAGAGGAGACCGTTCTCGTGTGGCTGCGGAAC |
| Pool_002_sgRNA_3 | GGGTCACGCGTAGGATTCTAATACGACTCACTATAGGAGTGAGTTTAATCTGGGCGTTTTAGAGCTAGAGGAGACCGTTCTCGTGTGGCTGCGGAAC |
| Pool_002_sgRNA_4 | GGGTCACGCGTAGGATTCTAATACGACTCACTATAGGGATGATAATGGCCCTGTTGTTTTAGAGCTAGAGGAGACCGTTCTCGTGTGGCTGCGGAAC |
| Pool_002_sgRNA_5 | GGGTCACGCGTAGGATTCTAATACGACTCACTATAGGAAACTTAGCGTAATATAAGTTTTAGAGCTAGAGGAGACCGTTCTCGTGTGGCTGCGGAAC |
| Pool_002_sgRNA_6 | GGGTCACGCGTAGGATTCTAATACGACTCACTATAGGTTCTGTGCTTAACAATGAGTTTTAGAGCTAGAGGAGACCGTTCTCGTGTGGCTGCGGAAC |
| Pool_002_sgRNA_7 | GGGTCACGCGTAGGATTCTAATACGACTCACTATAGGAGAGCGCCTCATTCATGTGTTTTAGAGCTAGAGGAGACCGTTCTCGTGTGGCTGCGGAAC |
| Pool_002_sgRNA_8 | GGGTCACGCGTAGGATTCTAATACGACTCACTATAGGTATTCGTATATGTGCGAAGTTTTAGAGCTAGAGGAGACCGTTCTCGTGTGGCTGCGGAAC |
| Pool_002_sgRNA_9 | GGGTCACGCGTAGGATTCTAATACGACTCACTATAGGACGGTGTCGCATTTGGAAGTTTTAGAGCTAGAGGAGACCGTTCTCGTGTGGCTGCGGAAC |
| Pool_002_sgRNA_10 | GGGTCACGCGTAGGATTCTAATACGACTCACTATAGGTTCGGATCATGTCAACTTGTTTTAGA |

| | |
|---|---|
| | GCTAGAGGAGACCGTTCTCGTGTGGCTGCG GAAC |
| Pool_002_sgRNA_11 | GGGTCACGCGTAGGATTCTAATACGACTCAC TATAGGATCTTAATGAGTATTGATGTTTTAGA GCTAGAGGAGACCGTTCTCGTGTGGCTGCG GAAC |
| Pool_002_sgRNA_12 | GGGTCACGCGTAGGATTCTAATACGACTCAC TATAGGTAGATAGACCTTTCACCGGTTTTAGA GCTAGAGGAGACCGTTCTCGTGTGGCTGCG GAAC |
| Pool_002_sgRNA_13 | GGGTCACGCGTAGGATTCTAATACGACTCAC TATAGGAAGAGGTTTGCGATCTGCGTTTTAG AGCTAGAGGAGACCGTTCTCGTGTGGCTGC GGAAC |
| Pool_002_sgRNA_14 | GGGTCACGCGTAGGATTCTAATACGACTCAC TATAGGTTATGAAAATTGTATGTCGTTTTAGA GCTAGAGGAGACCGTTCTCGTGTGGCTGCG GAAC |
| Pool_002_sgRNA_15 | GGGTCACGCGTAGGATTCTAATACGACTCAC TATAGGATCCGATTACGACAAATAGTTTTAGA GCTAGAGGAGACCGTTCTCGTGTGGCTGCG GAAC |
| Pool_002_sgRNA_16 | GGGTCACGCGTAGGATTCTAATACGACTCAC TATAGGTCTAATACCGCACTCTCTGTTTTAGA GCTAGAGGAGACCGTTCTCGTGTGGCTGCG GAAC |
| Pool_002_sgRNA_17 | GGGTCACGCGTAGGATTCTAATACGACTCAC TATAGGAACGCGTCCCGTTCATCGGTTTTAG AGCTAGAGGAGACCGTTCTCGTGTGGCTGC GGAAC |
| Pool_002_sgRNA_18 | GGGTCACGCGTAGGATTCTAATACGACTCAC TATAGGTGACTCTGCTTAGCCTATGTTTTAGA GCTAGAGGAGACCGTTCTCGTGTGGCTGCG GAAC |

**Table S5.** Primer pairs and conditions for subpooling and bulk-amplifying ten target oligo ssDNA libraries of various scales from microarray-synthesized chip9 oligonucleotides. Annealing temperatures (Tm) for each subpooling primer pair, using Kapa HiFi polymerase, were obtained by gradient qPCR. The table also includes the number of PCR cycles corresponding to the qPCR amplification plateau for subpooling and bulk amplification, before and after cycle number optimization. Each target oligo subpool was T7 *in vitro* transcribed to produce ten sgRNA libraries.

| Library scale | Subpool | Forward Primer Name | Forward Primer Sequence | Reverse Primer Name | Reverse Primer Sequence | Tm* (°C) | Subpool Cycles (Original) | Subpool Cycles (Optimized) | Bulk Amp Cycles (Original) | Bulk Amp Cycles (Optimized) |
|---|---|---|---|---|---|---|---|---|---|---|
| 342 | 1 | skpp15-1-F filt15-1651 | GGGTCACGCGTAGGA | skpp15-1-R filt15-1181 | GTTCCGCAGCCACAC | 51 | 24 | N/A | 17 | N/A |
| 206 | 4 | skpp15-4-F filt15-656 | GGTCGAGCCGGAACT | skpp15-4-R filt15-740 | GGATGCGCACCCAGA | 52 | 26 | N/A | 16 | N/A |
| 1,382 | 6 | skpp15-6-F | CGCAGGGTCCAGAGT | skpp15-6-R | GTTCGCGCGAAGGAA | 50 | 22 | 15 | 16 | 12 |
| 1,254 | 7 | skpp15-7-F filt15-1855 | AGTGACCCGTCCCTG | skpp15-7-R filt15-757 | AGTCGACCTCTGCCC | 51 | 24 | N/A | 17 | N/A |
| 1,753 | 9 | skpp15-9-F filt15-453 | CGATCGTGCCCACCT | skpp15-9-R filt15-1189 | GTGCGGGCTCCAACT | 58 | 24 | N/A | 17 | N/A |
| 1,237 | 13 | skpp15-13-F | GGGTTCGAGCGGGAG | skpp15-13-R | TAGCGCGCAGAGAGG | 56 | 20 | N/A | 16 | N/A |
| 389 | 23 | skpp15-23-F filt15-577 | AGCTGCTACACCGCC | skpp15-23-R filt15-1596 | GCGCGATGGTCACAG | 53 | 22 | 17 | 17 | 12 |
| 227 | 26 | skpp15-26-F | GCGGCACCACAAACT | skpp15-26-R | CGTGGCCTCTGTCCT | 52 | 23 | N/A | 17 | N/A |
| 2,626 | 27 | skpp15-27-F | ACCTTCACGCGTCCC | skpp15-27-R | GCCCACCGACTCCAC | 51 | 20 | 16 | 18 | 7 |
| 2,223 | 28 | skpp15-28-F | GACTGCGGCGTTGGT | skpp15-28-R | TACGCCCGGGACAGA | 52 | 22 | N/A | 16 | N/A |

*Annealing temperatures listed correspond to Kapa HiFi Polymerase

**Table S6.** Additional nucleic acid sequences that were not used for subpooling amplification or 5' RACE in this study.

| Sequence name | Sequence | Description |
|---|---|---|
| ds_sgRNA_scaffold_BsaI_YG | GAGAACGGTCTCCTAGAAAT AGCAAGTTAAAATAAGGCTAG TCCGTTATCAACTTGAAAAAG TGGCACCGAGTCGGTGCTTT T | Duplexed sgRNA scaffold DNA sequence from IDT |
| SgRNA_GGA_oligo_FWD_NV | GGGTCACGCGTAGGATTCTA ATACG | Forward primer for amplifying sgRNA templates (GGA products) for IVT |
| sgRNA_GGA_oligo_REV_NV | AAAAGCACCGACTCGGTGCC AC | Reverse primer for amplifying sgRNA templates (GGA products) for IVT<br>Also used for RSPE of of S2 and S4 oPools (1 PCR cycle) |
| *S. pyogenes* control sgRNA sequence | mC*mA*mU* rCrCrU rCrGrG rCrArC rCrGrU rCrArC rCrCrG rUrUrU rUrArG rArGrC rUrArG rArArA rUrArG rCrArA rGrUrU rArArA rArUrA rArGrG rCrUrA rGrUrC rCrGrU rUrArU rCrArA rCrUrU rGrArA rArArA rGrUrG rGrCrA rCrCrG rArGrU rCrGrG rUrGrC mU*mU*mU* rU | Control sgRNA sequence from the EnGen® sgRNA Synthesis kit |

**Table S7.** Primer pairs and conditions are provided for subpooling four 98 nt oligo libraries of varying sizes, with either one or three guanines downstream of the T7 promoter. These libraries were subpooled from the e13sgRNA oPool, which contains 389 unique target oligos. Each 98 nt oligo includes a library-specific forward primer site, T7 promoter, 20 nt spacer, and reverse primer site. Annealing temperatures (Tm) for each subpooling primer pair, using Kapa HiFi polymerase, were obtained by gradient qPCR. It also includes the number of PCR cycles corresponding to the qPCR amplification plateau for subpooling. Each target oligo subpool was *in vitro* transcribed to produce four sgRNA libraries.

| Subpool name (plexity, number of guanines) | Forward Primer Name | Forward Primer Sequence | Reverse Primer Name | Reverse Primer Sequence | Tm* (°C) | Subpool PCR Cycles |
|---|---|---|---|---|---|---|
| 12G | skpp15-6-F | CGCAGGGTCC AGAGT | skpp15-6-R | GTTCGCGCGA AGGAA | 50 | 11 |
| 12G4 | skpp15-13-F | GGGTTCGAGC GGGAG | skpp15-13-R | TAGCGCGCAG AGAGG | 56 | 12 |
| 60G4 | skpp15-23-F filt15-577 | AGCTGCTACA CCGCC | skpp15-23-R filt15-1596 | GCGCGATGGT CACAG | 53 | 10 |
| 389G4 | skpp15-27-F | ACCTTCACGC GTCCC | skpp15-27-R | GCCCACCGAC TCCAC | 51 | 7 |

*Annealing temperatures listed correspond to Kapa HiFi Polymerase

**Table S8.** Sequences used for the 5' RACE protocol in RNA-seq of sgRNA libraries, including RT primers, template-switching oligos, and gene-specific primers.

| Sequence Name | Sequence | Description |
|---|---|---|
| RT_primer_YG | AGCATATATCCCGGTCTGGA NNNNNNNNNNNNNNNNNN AAAAGCACCGA | Reverse transcription DNA primer |
| TSO_YG | GCTAATCATTGCAAGCAGTGGTATCAACGCAGAGTACATrGrGrG | Template Switching Oligo (TSO) |
| TSO_primer_YG | CATTGCAAGCAGTGGTATCAAC | DNA primer |
| gene_specific_primer_YG | AGCATATATCCCGGTCTGGA | DNA primer |
| sgRNA_cDNA_gibson_FWD_YG | TCCAGACCGGGATATATGCTtatgcggtgtgaaataccgcac | Forward DNA primer |
| sgRNA_cDNA_gibson_REV_YG | CACTGCTTGCAATGATTAGCcggtatcattgcagcactgg | Reverse DNA primer |
| sgRNA_cDNA_1kbp_ext_FWD_YG | AGAGTTCTTGAAGTGGTGGC | Forward DNA primer |
| sgRNA_cDNA_1kbp_ext_REV_YG | GTGTGGAATTGTGAGCGGAT | Reverse DNA primer |

To synthesize sgRNA libraries, we first computationally select spacer sequences from barcode protospacers within DropSynth gene libraries. We add a T7 promoter for IVT and BsaI restriction sites for golden gate assembly to these spacer sequences to generate target oligos. We order all target oligos as a single OLS pool. We subpool libraries from the OLS pool via PCR and perform golden gate assembly to join these target oligos to a conserved, duplexed sgRNA scaffold sequence. We add this sgRNA template to an IVT mix with T7 RNA polymerase to generate the sgRNA pools. These are verified via TBE-urea gel electrophoresis and RNA-seq.

Methods

1. **Subpool sgRNA target oligo libraries from an OLS pool.**
    a. Resuspend and dilute the chip pool with oligos produced via oligonucleotide library synthesis (OLS).
        i. Follow resuspension guidelines provided by the manufacturer.
        ii. Briefly centrifuge the tube opening and resuspend in nuclease free Tris-EDTA (TE) buffer, pH 8.0 to the desired concentration. A concentration of at least 10 ng/µL is recommended for the stock dilution.
        iii. Dilute the stock OLS chip pool 1:10.
        iv. Prepare 10 µM forward and reverse subpool amplification primers for each subpool.
    b. Amplify subpools
        i. For each subpool, run a qPCR to determine the number of cycles required for amplification. Amplifications are stopped several cycles before plateauing to prevent over-amplification of the libraries. Alternatively, qPCR can be run for 30 cycles solely for determination of cycles required and the amplification product can be discarded. Aim to obtain less than 15 cycles to prevent overamplification and biased propagation of specific spacer sequences.
        ii. Kapa HiFi or Kapa SYBR fast can be used for this step.
        iii. Prepare qPCRs in duplicate for each subpool (1,382-plex oligo library in this case) and corresponding primer pair (Table S5).

| **Component** | **1X reaction (25 µL final vol)** |
|---|---|
| Nuclease-free water to fill | 7.5 µL |
| KAPA HiFi HotStart ReadyMix (2X) | 12.5 µL |
| Fwd primer (10 uM)<br>skpp15-6-F | 1.25 µL |
| Rev primer (10 uM)<br>skpp15-6-R | 1.25 µL |
| Chip9 OLS 1:10 (0.43 ng/µL) | 2.5 µL (1 ng) |
| Thiazole green (100X) | 0.25 µL |

        iv. Run according to subpool PCR protocol. The table below shows the qPCR protocol for subpooling the 1,382-plex oligo library.

| **Cycles** | **Step** | **Temperature** | **Time** |
|---|---|---|---|
| 1 | Denaturation | 98°C | 30 s |
| 40 | Denaturation | 98°C | 10 s |
| | Annealing | 50°C (for skpp15-6 primers) | 15 s |
| | Extension | 72°C | 15 |

    c. Bulk amplify subpools. Use the OLS pool as the template for this step. The purpose of this step is to obtain sufficient subpooled DNA for downstream bulk amplification.
    d. Prepare 2-3X PCRs per subpool (1,382-plex oligo library in this case). Scale up master mixes as necessary.

| Component | 1X reaction (25 µL final vol) |
|---|---|
| Water to fill | 9.75 µL |
| KAPA HiFi HotStart ReadyMix (2X) | 12.5 µL |
| Fwd primer (10 µM)<br>skpp15-6-F | 1.25 µL |
| Rev primer (10 µM)<br>Skpp15-6-R | 1.25 µL |
| Chip9 OLS 1:10 (0.43 ng/µL) | 2.5 µL (1 ng) |

e. Run according to subpool PCR protocol. However, use the number of cycles determined in step 1, 2 (15 in this case) to denature, anneal, and extend the subpooled DNA. The table below shows the qPCR protocol for subpooling the 1,382-plex oligo library.

| Cycles | Step | Temperature | Time |
|---|---|---|---|
| 1 | Denaturation | 98°C | 30 s |
| 15 | Denaturation | 98°C | 10 s |
| | Annealing | 50°C | 15 s |
| | Extension | 72°C | 15 s |
| 1 | Final extension | 72°C | 1 min |
| | Hold | 10°C | infinite |

f. Elute each subpool using one 5 µg DNA clean-up column with 10 µL hot elution buffer.
g. Use five times the volume of binding buffer as the volume of your pooled PCRs.
h. Run PCR products on a gel and look for high molecular weight products that are indicative of overamplification. Also check for excessive low molecular weight products that may indicate chip synthesis issues.
i. Size select the correct PCR products using gel extraction if necessary.
j. Quantify yield amount using 1X hs dsDNA Qubit kit. Aim to obtain at least 80 ng total for the next step.

2. **Bulk amplify sgRNA target oligo subpools.** Use amplified subpools as the template for this step. The purpose of this step is to obtain sufficient subpooled DNA for downstream golden gate assembly to join scaffold sgRNA sequence.
   a. For each subpool, run a qPCR to determine the number of cycles required for bulk amplification. Amplification is stopped several cycles before plateauing to prevent over-amplification of the libraries. Alternatively, qPCR can be run for 30 cycles solely for determination of cycles required and the amplification product can be discarded. Aim to obtain less than 10 cycles to prevent overamplification and biased propagation of specific spacer sequences.
      i. Kapa HiFi or Kapa SYBR fast can be used for this step.
      ii. Prepare qPCRs in duplicate for each subpool and corresponding primer pair (Table S5). Scale up master mix as needed.

| Component | 1x reaction (25 µL reactions) |
|---|---|
| Nuclease-free water to fill | 9 µL |
| Kapa HiFi Master Mix | 12.5 µL |
| FWD primer (10 µM) | 1.25 µL |
| REV primer (10 µM) | 1.25 µL |
| Lib subpool PCR product<br>Add 10 ng total (1,382-plex lib) | 0.8 µL (10 ng) |
| Thiazole green (100X) | 0.25 µL |

iii. Run according to the subpooling qPCR protocol on step 1e.

b. Bulk amplify each subpool. Scale up master mixes and aim to prepare 8X PCRs per subpool.

| **Component** | **1x reaction (25 µL reactions)** |
|---|---|
| Water to fill | 9.25 µL |
| Kapa HiFi Master Mix | 12.5 µL |
| FWD primer (10 µM) | 1.25 µL |
| REV primer (10 µM) | 1.25 µL |
| Lib subpool PCR product<br>Add 10 ng total (1,382-plex lib) | 0.8 µL (10 ng) |

i. Run according to subpool PCR protocol. However, use the number of cycles determined in step 2,a, iii (6 cycles in this case) to denature, anneal, and extend the subpooled DNA. The table below shows the PCR protocol for bulk amplifying S2 lib 15.

| **Cycles** | **Step** | **Temperature** | **Time** |
|---|---|---|---|
| 1 | Denaturation | 98°C | 30 s |
| 6 | Denaturation | 98°C | 10 s |
| | Annealing | 50°C | 15 s |
| | Extension | 72°C | 15 s |
| 1 | Final extension | 72°C | 1 min |
| | Hold | 10°C | Infinite |

ii. Column-clean PCR products using Monarch DNA clean up kit.

1. Use five times the volume of binding buffer as the volume of your pooled PCRs.
2. Elute each subpool using one miniprep clean-up column (NEB Cat: (NEB Cat: T1017-2) with 30 µL hot elution buffer.
3. Measure concentration with Qubit kit and aim to obtain at least 315 ng of DNA for the next step.

3. **Prepare for Golden Gate Assembly (GGA) of sgRNA target oligo subpools and scaffold oligo.** Prepare 5X assemblies per subpool. Use a 2:1 scaffold oligo: target oligo subpool ratio.

a. Dilute scaffold duplex (100 µM) 1:100.

i. Add 1 µL scaffold to 100 µL water. This should be about 30 ng/µL.

b. Drop dialyze target oligo subpools to remove any residual salts that may reduce T4 ligase activity.

i. Place a 0.05 µm membrane filter on a 245 mm x 245 mm dish (Polystyrene Corning) filled with DI water.

ii. Pipette subpooled DNA onto the membrane filter.

iii. Allow DNA to dialyze for 20 minutes.

iv. Pipette DNA into clean tube.

c. Prepare GGAs according to the table below. Scale up master mixes as necessary and aliquot the reactions into PCR tubes.

| **Component** | **GGA 1x (25 µL total volume)** |
|---|---|
| Duplexed scaffold oligo (83 bp)<br>Add 2 pmol/rxn, | 4 µL (~111 ng) |
| Subpooled oligo library DNA (98 bp)<br>Add 1 pmol/rxn | ___ µL (~63 ng) |
| 10X T4 DNA ligase buffer (NEB B0202S)<br>Add 1:1 with supplemental 1 mM ATP (NEB P0756) | 5 µL buffer mix |

| | |
|---|---|
| T4 DNA ligase (NEB M0202) | 0.25 µL |
| BsaI-HFv2 (NEB R3733) | 0.75 µL |
| Nuclease-free water to fill | To fill |

d. Perform the GGA protocol shown on the table below.

| Step | Temp | Time |
|---|---|---|
| | 37°C | 5 min |
| 100 cycles | 16°C | 5 min |
| Heat inactivation | 80°C | 20 min |
| | 12°C | hold |

e. Column-clean GGA products using Monarch DNA clean up kit.
   i. Use five times the volume of binding buffer as the volume of your pooled GGAs.
   ii. Elute each subpool using one 5 µg DNA clean-up column with 20 µL hot elution buffer.
   iii. Measure concentration with Qubit kit and aim to obtain at least 200-300 ng of DNA for the next step (2-3 pmol).

4. **Perform *in vitro* sgRNA synthesis**. Use GGA products from step 3 as the template.
   a. Prepare one *in vitro* transcription reaction per subpool according to the table below. Avoid preparing master mixes and add each reagent in the order listed to PCR tubes that are on a cooling rack on ice. Include a 0 pmol (no DNA) control.

| **Component** | **Volume** | **0 pmol control** | **Library IVT** |
|---|---|---|---|
| NEB Murine RNase inhibitor (NEB M0314) | 0.5 µL | 0.5 µL | 0.5 µL |
| Nuclease-free water | To fill | 11.5 µL | To fill |
| GGA products (full-length sgRNA oligos) 135 bp<br>2-3 pmol if possible | Up to 11.5 µL | -- | Up to 11.5 µL |
| DTT (100 mM) | 2 µL | 2 µL | 2 µL |
| NEB 10x reaction mix (NEB B9012SVIAL) | 2 µL | 2 µL | 2 µL |
| Ribonucleotide mix NEB (NEB N0466L) | 2 µL | 2 µL | 2 µL |
| NEB T7 RNA polymerase (NEB M0251L) | 2 µL | 2 µL | 2 µL |
| Total volume | 20 µL | 20 µL | 20 µL |

   b. Mix reactions well and spin them down. Incubate at 37°C for 2 hours.
   c. Treat each reaction with DNase I (NEB M0303) according to the protocol on the table below. Add water first, then buffer, then DNase while samples are on a cooling rack on ice.

| Component | Per 50 µL reaction |
|---|---|
| Transcribed sgRNA | 20 µL reaction from 3 |
| DNase I reaction buffer (10x) | 5 µL |
| DNase I (RNase-free) | 0.5 µL (1 unit) |
| Nuclease-free water | 24.5 µL |

i. Mix reactions well and spin them down. Incubate at 37°C for 10 minutes.
ii. Column-clean the transcribed sgRNAs with Monarch RNA kit (NEB Cat: T2040). Elute with 25 µL nuclease-free water into RNase-free 1.5 mL tubes.
iii. Quantify RNA yield with Qubit HS RNA kit (ThermoFisher Cat: Q32852)

d. Run sgRNAs on a TBE-UREA (10% polyacrylamide) denaturing gel (Bio-Rad 4566036) to confirm sgRNA size. Load each sample with 2X RNA loading dye containing 47.5% Formamide (NEB Cat: B0363S) and run the gel alongside a low range ssRNA ladder (NEB Cat: N0364S). Post-stain gel with SYBR gold nucleic acid stain (ThermoFisher Cat: S11494) and compare size of pooled sgRNAs to the NEB sgRNA control sequence. Expected sgRNA size is 100 bases.
e. Check for mutated spacers and sequence biases by performing RNA-seq of sgRNA pools.

**RNA-seq Master Protocol**

1. **Anneal the RT primer with RNA templates**
   a. Briefly centrifuge the Template Switching RT Enzyme Mix (NEB Product number: M0466S) to collect the solution to the bottom of tube, then place on ice.
   b. Thaw the Template Switching RT Buffer at room temperature completely. Vortex and centrifuge briefly to collect the solution to the bottom of tube, then place on ice.
   c. Prepare the reaction as follows (on ice) in a 0.2 mL nuclease free PCR tube (also make NTC):

| Reagent | Volume | Final Concentration |
|---|---|---|
| sgRNA | ___ µL | 10 ng – 1 µg |
| RT_primer_YG (10 µM) | 1 µL | 1 µM |
| dNTP (10 mM) | 1 µL | 1 mM |
| Murine RNase Inhibitor | 0.5 µL | - |
| Nuclease-free Water | To fill | - |
| Total Volume | 6 µL | - |

   d. Mix thoroughly by gently pipetting up and down at least 10 times, then centrifuge briefly to collect the solution to the bottom of the tube.
   e. Incubate for 5 minutes at 70°C in a thermocycler with the lid temperature set at ≥85°C, then hold at 4°C until the next step.

2. **Reverse Transcription (RT) and Template Switching**
   a. During the primer annealing reaction, vortex the Template Switching RT Buffer briefly followed by a quick spin Prepare the RT reaction mix as follows (adding RT Enzyme Mix last):
   b. Prepare the reactions as follows:

| Reagent | Volume | Final Concentration |
|---|---|---|
| Template Switching RT Buffer (4X) | 2.5 µL | 1X |
| TSO_YG (75 µM) | 0.5 µL | 3.75 µM |
| RT Enzyme mix (10X) | 1 µL | 1X |
| Total Volume | 4 µL | - |

c. Mix thoroughly by gently pipetting up and down at least 10 times, then centrifuge briefly to collect solution to the bottom of the tube.
d. Combine 4 µL RT reaction mix (above) with 6 µL of the annealed mix from step 3, mix well by pipetting up and down gently at least 10 times, then centrifuge briefly to collect the solution to the bottom of the tube.
e. Incubate the 10 µL combined reaction in a thermocycler.
f. Samples can be stored at 4°C overnight or at -20°C for up to a week.

| |
|---|
| 90 minutes at 42°C |
| 5 minutes at 85°C |
| Hold at 4°C |

3. **qPCR and PCR**
   a. Dilute the RT reaction from step 2 above by 2-fold with water and use 1 µL of the diluted cDNA in a subsequent 25 µL PCR reaction. For low abundant RNA targets, up to 2.5 µL of undiluted cDNA can be used in a 25 µL PCR reaction.
   b. Assemble the **qPCR** reaction on ice as follows:

| Components | Volume | Final Concentration |
|---|---|---|
| Diluted template switching cDNA product | 1 µL | - |
| Q5 Hot Start High-Fidelity Master Mix (2X) (NEB #M0494) | 12.5 µL | 1X |
| TSO_primer_YG (10 µM) | 1.25 µL | 0.5 µM |
| Gene_specific_primer_YG (10 µM) | 1.25 µL | 0.5 µM |
| H2O | 8.75 µL | - |
| Thiazole green (100X) | 0.25 | - |
| Total Volume | 25 µL | - |

   c. Mix gently by pipetting up and down at least 10 times, then centrifuge briefly to collect solution to the bottom of the tube.
   d. Incubate the reaction in a thermocycler with the lid temperature set at ≥100°C, and perform **qPCR** with the following cycling condition:

| Step | Temperature | Time | Cycles |
|---|---|---|---|
| Initial Denaturation | 98°C | 30 sec | 1 |

| | | | |
|---|---|---|---|
| Denaturation | 98 °C | 10 sec | 5 |
| Annealing & Extension | 72 °C | 5 sec (30 sec/kb) | |
| Denaturation | 98 °C | 10 sec | 5 |
| Annealing & Extension | 70 °C | 5 sec (30 sec/kb) | |
| Denature | 98 °C | 10 sec | 40 (25-35 recommended) |
| Annealing | **65 °C** | 15 sec | |
| Extension | 72 °C | 5 sec (30 sec/kb) | |
| Final Extension | 72 °C | 5 min | 1 |
| Hold | 4 °C | ∞ | |

e. Assemble the **PCR** reaction on ice as follows:

| Components | Volume | 10x | Final Concentration |
|---|---|---|---|
| Diluted template switching cDNA product | 1 µL | 10 µL | - |
| Q5 Hot Start High-Fidelity Master Mix (2X) (NEB #M0494) | 12.5 µL | 125 µL | 1X |
| TSO_primer_YG (10 µM) | 1.25 µL | 12.5 µL | 0.5 µM |
| Gene_specific_primer_YG (10 µM) | 1.25 µL | 12.5 µL | 0.5 µM |
| H2O | 8.75 µL | 87.5 µL | - |
| Total Volume | 25 µL | 250 µL | - |

f. Mix gently by pipetting up and down at least 10 times, then centrifuge briefly to collect solution to the bottom of the tube.

g. Incubate the reaction in a thermocycler with the lid temperature set at ≥100°C, and perform **PCR** with the following cycling condition:

| Step | Temperature | Time | Cycles |
|---|---|---|---|
| Initial Denaturation | 98°C | 30 sec | 1 |
| Denaturation | 98 °C | 10 sec | 5 |
| Annealing & Extension | 72 °C | 5 sec (30 sec/kb) | |
| Denaturation | 98 °C | 10 sec | 5 |
| Annealing & Extension | 70 °C | 5 sec (30 sec/kb) | |

| | | | |
|---|---|---|---|
| Denature | 98 °C | 10 sec | Cycle number determined by qPCR |
| Annealing | **65 °C** | 15 sec | |
| Extension | 72 °C | 5 sec (30 sec/kb) | |
| Final Extension | 72 °C | 5 min | 1 |
| Hold | 4 °C | ∞ | |

h. Store the PCR product at -20°C (no cleanup) for up to one week.
   i. Measure yield using HS DNA Qubit

4. Run samples on a 4% E-gel to check products (expect 186bp product). Load the samples according to the following gel key:
   a. NEB 50 bp ladder (100 ng/µL)
      i. Load 2 µL (200 ng) + 18 µL water
   b. sgRNA cDNAs
      i. Load 100 ng

5. **Linearize pUC19 and add overlaps for Gibson assembly via PCR**
   a. Dilute pUC19 from 1000 ng/µL to 100 ng/µL
   b. Assemble and run qPCR reaction (babysit for ~25 cycles):

| **Component** | **25 µl Reaction** | **Final Concentration** |
|---|---|---|
| Q5 Hot Start High-Fidelity 2X Master Mix | 12.5 µl | 1X |
| 10 µM sgRNA_cDNA_gibson_FWD_YG | 1.25 µl | 0.5 µM |
| 10 µM sgRNA_cDNA_gibson_REV_YG | 1.25 µl | 0.5 µM |
| Bayou Biolabs pUC19 (100 ng/µL) | 2 µµL | <5 ng |
| Nuclease-Free Water | 8 µl | |

| **STEP** | **TEMP** | **TIME** |
|---|---|---|
| Initial Denaturation | 98°C | 30 seconds |
| **25** Cycles | 98°C | 10 seconds |
| | **67°C** | 30 seconds |
| | 72°C | 45 seconds (20-30 sec/kb) |
| Final Extension | 72°C | 2 minutes |
| Hold | 4–10°C | |

   c. Clean up DNA using Monarch DNA clean-up kit.
   d. Elute in 26 µL nuclease-free water.
   e. Measure yield using HS DNA Qubit
   f. Assemble and run DpnI digestion of pUC19 template:

| Component | 50 µl Reaction |
|---|---|
| PCR product (1 µg) | 25 µL |
| 10X rCutSmart Buffer | 5 µl (1X) |
| DpnI | 1.0 µl (20 units) |
| Nuclease-free Water | 19 µl |

g. Incubate at 37°C for 15 minutes.
h. Clean up DNA using Monarch DNA clean-up kit.
   i. Elute in 26 µL of nuclease-free water.
i. Measure yield using the HS DNA Qubit kit.
j. Run a 1% E-gel (expect a 1.5 kb fragment)
   i. NEB 1 kb ladder (100 ng/µL)
      1. Load 2 µL (200 ng) + 18 µL water
   ii. 100 ng pUC19 fragment
      1. Load 100 ng

6. **Gibson assembly of digested pUC19 and sgRNA cDNA insert**
   a. Assemble Gibson assembly reaction as follows:

| | Assembly | Positive Control | Negative Control |
|---|---|---|---|
| pUC19 fragment (100 ng) | ___ µL * | - | ___ µL |
| sgRNA cDNA (42.07 ng) | ___ µL * | - | - |
| NEBuilder® Positive Control | - | 10 µl | - |
| Gibson Assembly Master Mix (2X) | 10 µL | 10 µL | 10 µL |
| Deionized H2O | To fill | - | To fill |
| Total Volume | 20 µL | 20 µL | 20 µL |

*Optimized cloning efficiency is 50–100 ng of vector with 2-3-fold molar excess of each insert.

   b. Incubate samples in a thermocycler at 50°C for 15 minutes.
   c. Clean up DNA using Monarch DNA clean-up kit.
   d. Elute in 12 µL elution buffer.
   e. Measure yield using HS DNA Qubit.

7. **PCR amplification of a 1.049 kbp fragment for sequencing**
   a. Assemble and run PCR reaction:

| **Component** | **25 µl Reaction** | **Final Concentration** |
|---|---|---|
| Q5 Hot Start High-Fidelity 2X Master Mix | 12.5 µl | 1X |
| 10 µM sgRNA_cDNA_1kbp_ext_FWD_YG | 1.25 µl | 0.5 µM |
| 10 µM sgRNA_cDNA_1kbp_ext_REV_YG | 1.25 µl | 0.5 µM |
| Gibson Assembly Product | ___ µL | 1ng-1µg |
| Nuclease-Free Water | To fill | |

| STEP | TEMP | TIME |
|---|---|---|
| Initial Denaturation | 98°C | 30 seconds |
| 25 Cycles | 98°C | 10 seconds |
| | **65°C** | 30 seconds |
| | 72°C | 25 seconds (20-30 sec/kb) |
| Final Extension | 72°C | 2 minutes |
| Hold | 4–10°C | |

b. Clean up DNA using Monarch DNA clean-up kit.
    i. Elute in 21 µL nuclease-free water or elution buffer.
c. Measure yield using HS DNA Qubit.
d. Run 2 % E-gel:
    i. NEB 1 kb plus ladder (100 ng/µL)
        1. Load 2 µL (200 ng) + 18 µL water
    ii. pUC19 fragment (expect 1,576 bp fragment)
    iii. Load 50 ng
    iv. cDNA (expect 186 bp product)
    v. Load 50 ng
    vi. Gibson Assembly product (expect 1,766 bp circular product)
    vii. Load 20 ng
    viii. PCR product (expect a 1,049 bp product)
    ix. Load 50 ng

8. **Submit for sequencing**
   Vendor: Plasmidsaurus (Oxford Nanopore Sequencing)
   1. Submission details:
       1. 1.014 kb, requires ≥10 uL of 30 ng/uL sample
       2. Request 2GB data.
